# Conditional and marginal SNP-heritability to leverage ancestral and environmental diversity

**DOI:** 10.64898/2026.05.28.728536

**Authors:** Anubhav Nikunj Singh Sachan, David Azriel, Armin Schwartzman

## Abstract

SNP-heritability is defined as the fraction of variance of a trait that is explained by the SNPs in a genome-wide association study. Several methodologies have been proposed to estimate this quantity. More recent methods aim to do so with ancestrally diverse datasets and yet obtain a single heritability for an entire dataset, which we refer to as marginal heritability. However, the different underlying subpopulations that compose a genetically diverse dataset might have different environmental and genetic exposures, and thus may have different heritabilities. In order to address this, we propose a conditional SNP-heritability approach that allows to estimate multiple SNP-heritabilities on a dataset corresponding to different ancestral compositions and environmental exposures. We take a careful statistical approach, including estimation of conditional genetic and environmental variances, and calculation of standard errors via a combination of the delta method with bootstrapping. We validate our method via extensive simulations. We then apply it to an ancestrally and socio-economically diverse dataset of 6603 subjects aged around 9 to 11 from the Adolescent Brain Cognitive Development study, and illustrate how the SNP-heritability of intelligence scores can change due to differing extrinsic variances in different socio-economic groups, which coincides with previous work in the literature. This conditional estimation approach can be a valuable tool for understanding differences in risks across subpopulations. Our work here improves on existing methodology and allows us to leverage the heterogeneity of the data to obtain new insights.

## 1 Introduction

SNP-heritability is defined as the fraction of variance of a trait that is explained by the SNPs in a genome-wide association study (GWAS). Several methodologies have been proposed to estimate this quantity [38]. The most popular is LD-score regression (LDSC) [3]. Another relevant estimator to our work is the GWAS heritability estimator (GWASH) [33], which is similar to LD-score regression with fixed intercept (LDSC-1) but with a closed form estimate for heritability and standard error. These methods require samples from homogenous populations [18]. More recent methods aim to estimate SNP-heritability with ancestrally diverse datasets. An example is covariate adjusted LD-score regression (cov-LDSC) [18], which is an extension of LD-score regression and was recently released to better handle samples with admixed individuals [18]. The method works by regressing out principal components from each of the SNPs included in the analysis (creating covariate-adjusted SNPs) and regressing out the same principal components from the phenotypes, and then applying LDSC where the LD scores are obtained from the LD matrix of the covariate-adjusted SNPs.

Although cov-LDSC and similar methods address some of the issues present when estimating SNP-heritability for ancestrally diverse populations, such as LD matrices with long-range correlations, they only obtain single estimates of heritability for an entire sample. However, due to varying environmental effects and differences in the frequency of different genetic markers, the heritability of a trait might not be the same across different ancestries and sub-populations present in a sample. An example of this can be seen in the study conducted in [39] that found that the heritability of IQ-scores in children was modified by socioeconomic status, with children in lower income households experiencing increased environmental variance. Since heritability measures the fraction of variance explained by genetic factors and the total variance has both genetic and environmental components, then an increased environmental variance results in a decreased heritability, even if the genetic component remains constant.

Therefore, when estimating heritability with ancestrally diverse datasets, one should account for co-variates that can affect it. These covariates may be genetic, such as ancestry, or environmental, such as socio-economic income. In this article we directly address estimation of SNP-heritability when working with datasets where such covariates are present.

First we give a proper definition of a single SNP-heritability in the presence of covariates, which we call *marginal heritability*, and then define a *conditional heritability* which is a function of the covariates, allowing the SNP-heritability value to change from individual to individual depending on their ancestry or environmental characteristics. We propose formal statistical estimators of these quantities. The estimator for marginal heritability works similarly to cov-LDSC, by regressing out covariates from phenotypes and SNPs. However, as opposed to cov-LDSC, which only works with LD-score regression, our method allows any valid heritability estimator such as GWASH or LDSC-1 on the covariate adjusted phenotypes and SNPs. The estimator for conditional heritability breaks down the estimand into five parameters, some of which are functions of covariates, and estimates each of these. We also provide a framework for the calculation of standard errors for conditional heritability estimates via a combination of the delta method with bootstrapping.

We then apply our methods to an ancestrally and socio-economically diverse dataset of 6603 subjects aged around 9 to 11 from the Adolescent Brain Cognitive Development (ABCD) study (https://abcdstudy.org/about/), and illustrate how the SNP-heritability of intelligence scores can change due to differing extrinsic variances in different socio-economic groups, which coincides with previous work with twin studies in the literature [39], as well as a recent study with the ABCD dataset that studied the SNP-heritability of cortical structure and cognitive ability [25].

To validate our estimation framework we run an extensive simulation study, varying different parameters such as marginal heritability, number of ancestries and ancestry effects. We show that the estimated marginal and conditional heritability values match their theoretical values under our framework. We also compare our calculation of standard errors of the conditional heritability estimates to Monte Carlo standard error values, and show that they are close to each other. All of these simulations demonstrate the robustness of our heritability estimation methods to a variety of scenarios.

To the best of our knowledge, our estimation framework for conditional heritabilities is not present in the current literature. Our proposed approach can be a valuable tool for understanding differences in risks across subpopulations. Our work here improves on existing methodology and allows us to leverage the heterogeneity of the data to obtain new insights.

## 2 Methods

### 2.1 Underlying polygenic model

We shall describe a conditional approach for taking population stratification and potentially other covariates into consideration when calculating heritability. To justify our approach, we focus on the following phenotype model that takes into account polygenic and covariate effects:

#### Assumption 1

(Phenotype model with covariates). *Assume that* ***x*** *is a random vector of length m that represents the sequenced SNPs and that y is a continuous trait or phenotype of interest. Also assume that there are K covariates to account for and take* ***s*** *to be a random vector of length K that represents the covariates for that particular person. Suppose that the relationship between y*, ***x*** *and* **s** *can be modelled as a linear regression:*

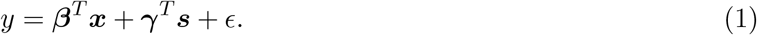

*where* ***β*** *and* ***γ*** *are fixed. Let* Var(***β***^*T*^ ***x***|***s***) = *τ* ^2^(***s***) *and* Cov(***x***|***s***) = **Σ**(***s***), *and assume E*(*ϵ*|***s***) = 0, Var(*ϵ*|***s***) = *σ*^2^(***s***) *and that conditional on* ***s*** *we have that ϵ is independent of* ***x***.

The vector ***s*** can contain ancestry variables, but also other covariates that are potentially associated with the trait of interest such as age, income and sex. We also note that under Assumption 1, the conditional variance of a phenotype given covariates of interest can be decomposed as

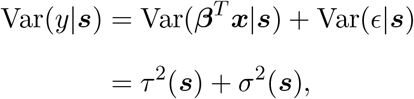

where *τ* ^2^(***s***) = Var(***β***^*T*^ ***x***|***s***) captures the genetic variance component and *σ*^2^(***s***) = Var(*ϵ*|***s***) captures the extrinsic non-genetic variance component. Moreover, under this same model, the genetic component can be expressed as

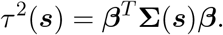

### 2.2 Covariate-adjusted marginal heritability

One way to obtain a heritability estimate from the model described in Equation (1) would be to apply cov-LDSC [18]. This method works by regressing out the covariates of interest from the observed phenotype and SNPs of interest and then running LDSC [3] using these residualized inputs. If we were to generalize this approach, we could apply any typical heritability estimator such as GWASH instead of LDSC. More details about this are provided in the “Estimation setup” of Section 2.5 below on and in Appendix B. This process leads to an estimate of what we shall refer to as covariate-adjusted marginal heritability 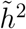 defined as follows:

#### Definition 1

(Covariate-adjusted marginal SNP heritability). *Assume the model described in Assumption 1. We define covariate-adjusted marginal SNP-heritability as:*

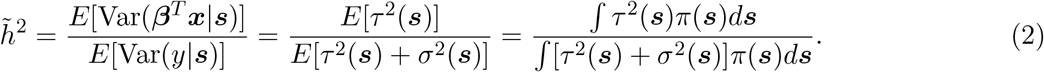

*where π*(***s***) *represents the joint probability density function for* ***s*** *and the expectation is taken over the distribution of* ***s***.

Appendix B has a derivation to justify the use of residualized inputs to estimate the covariate-adjusted marginal heritability 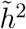 from Definition 2. We refer to this quantity as covariate-adjusted because its definition accounts for the presence of the covariates in Equation (1), and we refer to it as marginal because the expected values in the numerator and denominator reduce the estimation to a single quantity for an entire population. That said, for simplifying terminology we shall also refer to 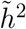 as simply **marginal SNP-heritability** or **marginal heritability**.

### 2.3 Covariate-adjusted conditional heritability

Although a marginal heritability estimator allows one to obtain a heritability estimate for an ancestrally diverse sample, it still only obtains a single heritability estimate for an entire dataset. However, the different underlying subpopulations that compose an ancestrally diverse sample might have different environmental and genetic exposures. This motivates us to focus on a conditional framework.

To understand this better, let us say that there are only two ancestral populations in the data, for example, East Asian and European (assume for now these cannot be further subdivided and that there are no individuals with mixed ancestry). Our model from Equation (1) reduces to

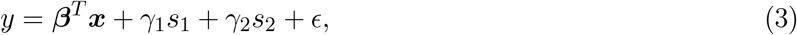

where *s*_1_ takes a value of 0 if the random individual is East Asian and 1 if the random individual is European. Analogously, *s*_2_ takes a value of 0 if the random individual is European and 1 if the random individual is East Asian. We take the covariance of ***x*** for the European population to be **Σ**_1_ and the covariance of ***x*** for the East Asian population to be **Σ**_2_. Under Assumption 1, the conditional variance for the European sub-cohort reduces to 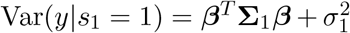 and the conditional variance for the East Asian sub-cohort reduces to 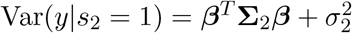.

We can then define two heritabilities - one for the European sub-cohort 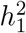 and another one for the East Asian sub-cohort 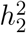,

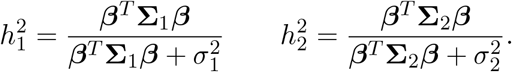

We can also denote this as

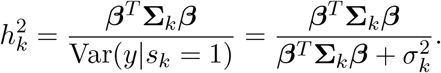

This is an example of what we will call **covariate-adjusted conditional SNP-heritability** - an individual’s heritability estimate is dependent on their ancestry and potentially other covariates. Referring back to Equation (1), we can extend the definition of this more formally to allow for more than two ancestries and admixture as follows:

#### Definition 2

(Covariate-adjusted conditional SNP heritability). *Assume the model described in Assumption 1*. *We define covariate-adjusted conditional SNP-heritability as:*

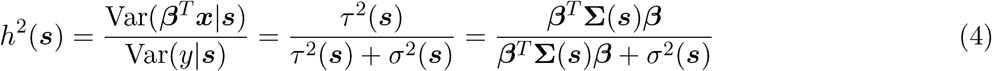

To simplify terminology, we shall proceed to also refer to *h*^2^(***s***) as just **conditional SNP-heritability** or **conditional heritability**. We note that the covariate-adjusted marginal heritability from Definition 1 takes an expectation with respect to ***s*** of the numerator and denominator and thus differs from Definition 2.

For the two-population case with no admixture described above, we would have ***s*** = (1, 0) for European ancestry and ***s*** = (0, 1) for East Asian ancestry, and we would take **Σ**(0, 1) = **Σ**_1_ and **Σ**(1, 0) = **Σ**_2_. Defining conditional heritability as in Definition 2 has the advantage of being able to also account for admixed individuals. For example, if someone had 30% European ancestry and 70% East Asian ancestry then we could represent this as ***s*** = (0.3, 0.7). Moreover, we can also use this definition if there are covariates other than ancestry - these would be part of vector ***s*** in Equation (1). For example, we could use covariates such as age, income and sex; and we could use principal components in lieu of genetic ancestry proportions - a common practice since [27] suggested their use to account for stratification in genome-wide association studies.

It is worth clarifying that the term “conditional heritability” exists with a different meaning in [41]. Their work only provides a procedure for estimating a single heritability which does not vary with covariates of interest, and thus in our context, it is understood as a type of a marginal heritability.

Similar to Definition 2, we also define the **conditional extrinsicality** as 1 − *h*^2^(***s***). This is the complement of the conditional heritability and represents the fraction of variance of the outcome that is **not** explained by genetics, conditional on the covariates ***s***.

### 2.4 Simplifying assumption for conditional SNP-heritability

In order to estimate conditional heritability we make a simplifying assumption that allows us to approximate the conditional heritability from Equation (4) as a function of five estimable parameters. Our assumption is as follows:

#### Assumption 2.

*[Simplifying assumption] Let t*(***s***) = tr(**Σ**(***s***)) *and a* = ||***β***||^2^*/m where m represents the number of sequenced SNPs. Assume that in addition to the framework described in Assumption 1, we have that* 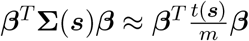 *and so:*

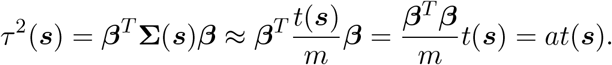

Then, following the details in Appendix D, we can obtain the following approximation:

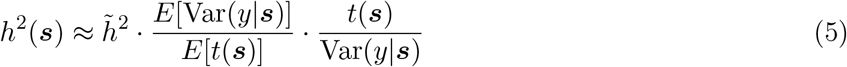

where the expectations are taken with respect to the distribution of ***s*** and 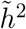 represents the marginal heritability from Equation (2).

This is an expression with the following five parameters: 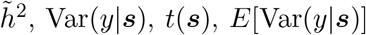 and 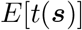.

We can obtain conditional heritability estimates by plugging in estimates for each of these, and we will describe the procedure to do so in the following subsection. We note that there is a relationship between conditional and marginal heritability, as one of the parameters required to estimate conditional heritability is the marginal heritability defined in Equation (2).

It is worth mentioning that Assumption 2 is already needed for existing methods that estimate heritability for homogeneous populations in the absence of covariates. For example the GWAS heritability (GWASH) estimator developed in [33] carries this assumption from the Dicker estimator [9] that it is based on. Moreover, although it is not obvious that LD-Score regression makes this assumption, it is hinted in [33] due to an asymptotic relationship to the GWASH estimator. From biological perspective, one way that this assumption can be satisfied is if the causal SNPs are considered random and areindependent from each other.

### 2.5 Parameter estimates

Suppose we have *n* individuals, *m* SNPs and *K* covariates of interest. Our observations of a phenotype of interest are stored in a column vector of size *n* which we denote with **y**. We also store our genotype observations in a *n* × *m* matrix denoted by **X**, where the element *X*_*ij*_ represents the value of the *j*-th SNP for the *i*-th individual. Similarly, our covariates are stored in a *n* × *K* matrix denoted by **S**, where the element *S*_*ik*_ represents the value of the *k*-th covariate for the *i*-th individual. We detail below the estimation of the five parameters of interest that appear in the conditional heritability approximation in Equation (5).

#### 2.5.1 Estimating 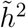

To obtain an estimate for 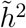 we first obtain pre-residualized phenotypes and genotypes 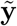 and 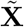 as follows:

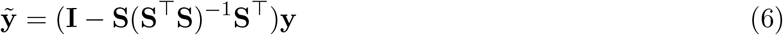

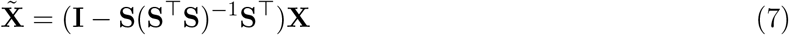

where **I** is an identity matrix. We then simply apply a heritability estimator such as GWASH or LDSC-1 to 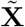 and 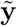. We denote this as 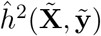.

Appendix B has a derivation to justify the use of 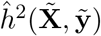 as an estimator for the marginal heritability defined in Definition 1. We note that we can equivalently obtain 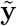 by setting it to the residuals after fitting a linear regression of **y** on covariates **S** using ordinary least squares (OLS). An analogous process can be used to obtain each column of 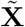.

In Section 2.6 we discuss the use of cov-LDSC as an estimator of marginal heritability. However in this article we also make use of the GWASH estimator because it has a closed-form formula for standard error estimates, which can be leveraged for the variance estimations of the conditional heritability estimates defined later on. Moreover, asymptotically there is a relationship between LDSC-1 and GWASH as described in [2] and [33], and from the simulations in Section F.2 we show that GWASH and LDSC-1 perform similarly for estimating marginal heritability. Because of this close relationship, it is sufficient to run GWASH and the results are likely to apply to LDSC-1. To coincide with the term used by [18] for their estimator, we shall sometimes refer to 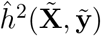 as **covariate-adjusted GWASH** when 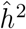 is taken to be the GWASH estimator.

#### 2.5.2 Estimating Var(*y*|s)

Let 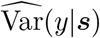 be the estimator for Var(*y*|***s***) and 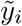 be the *i*-th element of 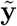. Note that due to the residualization from Equation (6), the vector 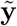 has a sample mean of 0. We define a vector **u** such that 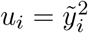, and fit a model for **u** using the covariate values stored in **S** as predictors. We then define *û*(***s***) to be the predicted value from this model for a subject with covariates equal to ***s***. We estimate 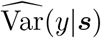 as

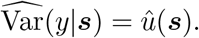

The model to use depends on the type of relationship that Var(*y*|***s***) has with ***s***. If the relationship is linear, then we can use OLS to fit a linear model of **u** on **S**. Otherwise, we can use a more flexible model such as a Generalized Additive Model (GAM) [15] [43] [44]. In practice, we recommend fitting both OLS and a GAM and comparing results.

#### 2.5.3 Estimating *t*(s)

Let 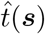 denote the estimator for *t*(***s***). We set **v** to be the diagonal of 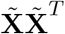 and, similar to above, we fit a model for **v** using the covariates values stored in **S** as predictors. Analogous to before, we define 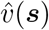 to be the predicted value from this model for a subject with covariates equal to ***s***. Our estimate 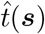 is

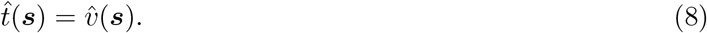

To understand the reasoning behind this estimate, note that *t*(***s***) can be expressed as *t*(***s***) = *E*[||***x*** − *E*[***x***|***s***]||^2^|***s***]. Moreover, 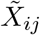 can be re-expressed as 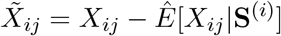 where **S**^(*i*)^ is the *i*-th row of **S** and *Ê*[*X*_*ij*_|**S**^(*i*)^] is a linear regression estimate for *E*[*X*_*ij*_|**S**^(*i*)^]. Thus 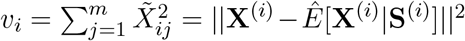 and so our proposed 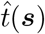 is a natural estimator for *t*(***s***).

Similar to before, we recommend fitting both OLS and GAM and comparing results.

#### 2.5.4 Estimating *E*[Var(*y*|s)]

Using our derivation in Appendix B, we can estimate *E*[Var(*y*|***s***)] as 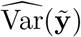 where 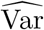 represents the sample variance. In other words,

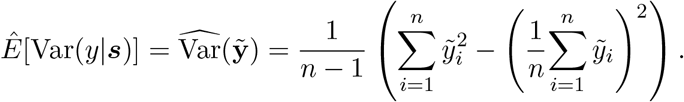

Due to residualization properties, 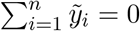 so:

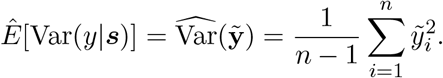

#### 2.5.5 Estimating *E*[*t*(s)]

We can obtain *Ê*[*t*(***s***)], an estimate for *E*[*t*(***s***)], as

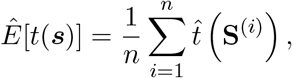

where **S**^(*i*)^ is the *i*-th row of **S** and 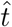 is a functional that has been estimated above in Equation (8).

### 2.6 Additional considerations

#### 2.6.1 Cov-LDSC as an estimator of covariate-adjusted marginal heritability

We mentioned previously that we can use GWASH applied to transformed SNPs and phenotypes to obtain an estimator for the covariate-adjusted marginal heritability 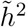. We note that covariate-adjusted LD score regression from [18] can also be used to estimate 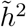 since it follows the process of regressing out covariates from **X** and **y** to obtain 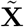 and 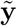, and then applies LDSC to these. This leads to an estimator of the form 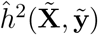 which estimates 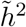 as discussed in Section 2.5.1.

This is interesting because our proposed theoretical framework and derivation is different from that discussed in [18]. In contrast to our proposed framework, the LD score regression paper [18] takes the effects ***β*** = (*β*_1_, …, *β*_*m*_)^⊤^ to be random and assumes a constant non-genetic variance 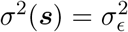. Under this latter framework, [18] derives the covariate-adjusted LD score regression as an estimator for

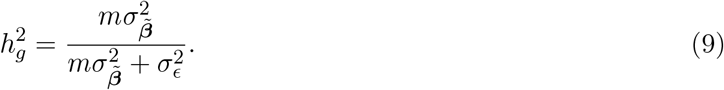

Here 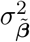 represents the variance of the vector 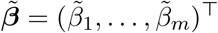 with each entry 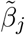 equal to 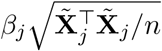 where 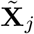 is the *j*-th column of 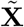. Apart from the difference in frameworks, we also note that unlike the marginal heritability 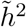 defined in Equation (2), 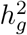 does not take an expectation of the numerator and denominator with respect to ***s***. Nonetheless, our simulation in Appendix F.2 confirms that covariate-adjusted LD-score regression using a fixed interecept remains a good estimator of our proposed covariate-adjusted marginal SNP heritability 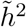 from Equation (2). This coincides with results from [2] and [33] that show an asymptotic equivalence between traditional GWASH and LDSC-1.

#### 2.6.2 Relationship between conditional heritability and adjusted marginal heritability estimates

Note that there is a relationship between our conditional heritability estimates and the covariate-adjusted marginal heritability estimates - when estimating conditional heritability, we need to estimate the covariate adjusted marginal heritability as one of its parameters.

We note that the conditional heritability estimator will be a functional estimate that can change depending on the values ***s***. We can also use the resulting functional to obtain estimates for values of ***s*** not observed in our sample. Changing the estimator used for calculating marginal heritability, will have the effect of transforming the conditional heritability estimates estimates by a multiplicative constant.

#### 2.6.3 Estimates for *τ* ^2^(s) and *σ*^2^(s)

When analyzing conditional heritabilities it can be of interest to understand whether any differences arise from the genetic component *τ* ^2^(***s***) or the non-genetic component *σ*^2^(***s***). Recalling Equation (5) and the first part of Equation (4), it is easy to show that:

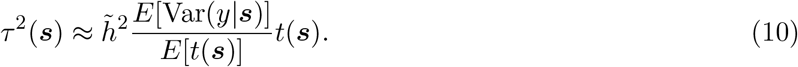

Thus, we can obtain the estimate 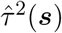 by plugging in the estimates we obtain for 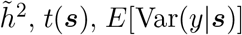 and *E*[*t*(***s***)] into Equation (10). Then we can obtain 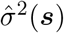 as

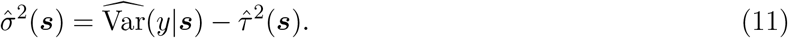

#### 2.6.4 Using summary statistics for marginal heritability estimation

Both the traditional LDSC and GWASH estimators can make use of summary statistics from a genome-wide association study. If using these for covariate-adjusted marginal heritability estimation with GWASH, some slight adjustments need to be made to the formulas presented in [33]. These are detailed in Appendix C.

### 2.7 Estimating the variance of conditional heritability estimates

One can estimate the variance of the conditional heritability estimator 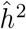(***s***) using a combination of the Delta Method and bootstrapping. Similar to 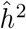(***s***), its variance estimate will also be a function of covariates ***s*** and takes the following form:

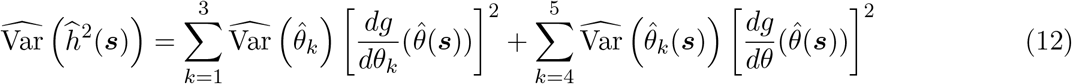

where 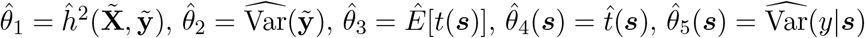 and

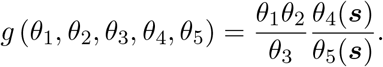

Full details are provided in Appendix E.

## 3 Results

### 3.1 Simulation study

We run simulation studies with 100 replications to examine the performance of the estimation of marginal and conditional heritabilities detailed above. A SNP matrix **X** with *n* = 1000 subjects and *m* = 2000 SNPs, and a corresponding phenotype vector **y** are generated during each replication. The ancestry proportions are assumed to be observed and the covariate matrix **S** is of dimensions *n* × *K* where each row corresponds to the ancestry proportions of each subject in the sample. We use the model from Equation (1) and take the number of ancestries to be *K* = 4, since this corresponds to the number of observed ancestries available for our real data analysis in Section 3.2. The ancestry proportions in **S** are generated only once at the beginning of the simulation study and kept fixed for the 100 replications. Since conditional heritabilities are dependent on the observed covariates, keeping the ancestry proportions fixed throughout the replications allows us to make sure that the conditional heritabilities do not change throughout replications.

To simulate the SNP effects ***β*** = (*β*_1_, …, *β*_*m*_), we randomly select 10 percent of them to be drawn from standard normal distributions and set the rest of them to be 0. This is done only once at the beginning of the simulation study and kept fixed for the 100 replications in accordance with Assumption 1. Furthermore, we generate different conditional heritabilities for each ancestry composition ***s*** by allowing the distribution of the SNPs ***x*** to be conditional on the ancestry composition ***s***, and model *σ*^2^(***s***) as a linear function of ***s***. Full details about the simulation setup, as well as simulations where additional parameters are varied, can be found in Appendix F.

#### 3.1.1 Estimation of marginal heritability

For each of the replications we calculate 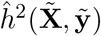 as described in Section 2.5 to estimate a pre-defined covariate-adjusted marginal heritability 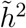 by setting 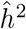 as the GWASH estimator. We explore the effect of varying 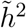 between 0.1 and 0.9, and summarize the results in Table 1. More details and simulations can be found in Appendix F. The naive GWASH reported in Table 1 is calculated using summary statistics from a GWAS that has accounted for covariates (and thus obtaining adjusted t-statistics and chi-square statistics), but using LD-scores where the SNPs have not been residualized with covariates. Since adjusting for principal components is standard for modern GWAS studies [21] [27], but covariate-adjusted LD scores have not been available before being discussed by [18], one can see such issues arising in practice and it is important to evaluate the effect of this discrepancy.

**Table 1:** Summary of the marginal heritability estimation simulation study for varying levels of 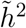 (target marginal heritability). We report the mean and standard deviation in parenthesis across the 100 repetitions. Any mean that is within one standard deviation of the target heritability is colored in blue, otherwise it is colored in red.

| Estimator | $\tilde{h}^2 = 0.1$ | $\tilde{h}^2 = 0.3$ | $\tilde{h}^2 = 0.5$ | $\tilde{h}^2 = 0.7$ | $\tilde{h}^2 = 0.9$ |
| --- | --- | --- | --- | --- | --- |
| Covariate-adjusted GWASH | 0.089<br>(0.048) | 0.283<br>(0.057) | 0.508<br>(0.065) | 0.725<br>(0.070) | 0.878<br>(0.067) |
| Naive GWASH | 0.033<br>(0.018) | 0.105<br>(0.021) | 0.190<br>(0.024) | 0.256<br>(0.025) | 0.303<br>(0.023) |

From Table 1 it can be seen that all mean estimates using covariate-adjusted GWASH are within one standard deviation of the corresponding target marginal heritability. Conversely, the naive approach underestimates the target marginal heritabilities for the scenarios reported here. Similar findings are described in [18] when comparing cov-LDSC with LDSC.

#### 3.1.2 Estimation of conditional heritability

Since ancestry proportions are continuous and kept fixed at the beginning of the simulation, each of the *n* = 1000 subjects will have a corresponding conditional heritability that will not vary across replications. In each of the 100 replications we calculate conditional heritability estimates for each subject as described in Section 2.5. We use these to get an average estimated conditional heritability for each of the *n* = 1000 subjects and compare these to the true values. Empirical standard deviations for the estimates are calculated too. Our results are summarized in Figure 3.1 which shows that all true conditional heritabilities are within one standard deviation of their average estimates. More details and simulations can be found in Appendix F.

**Figure 3.1:**
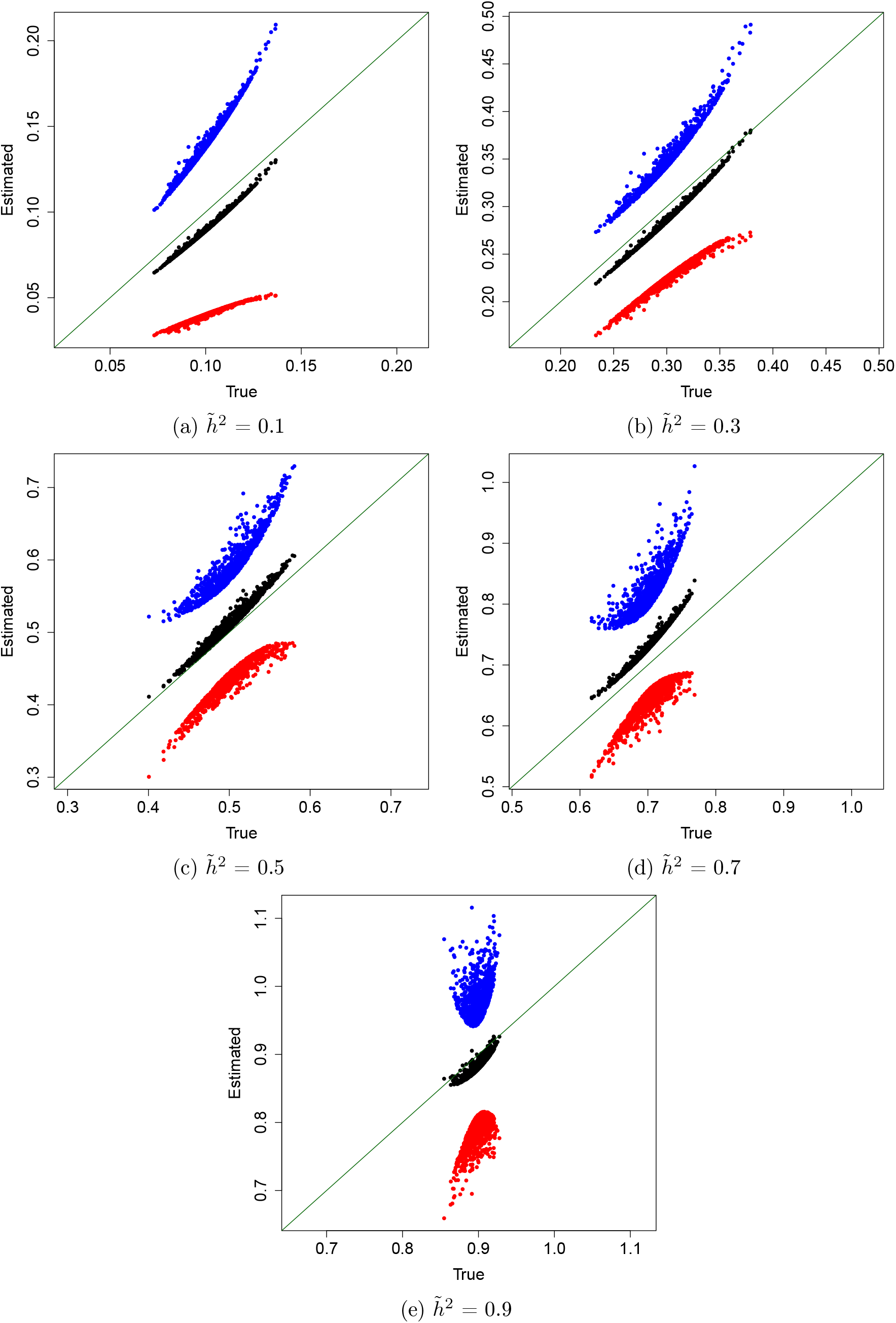
Comparisons of true conditional heritabilities vs their average estimates (black). Plus and minus one empirical standard deviations of the 100 simulations are plotted in the blue and red dots, respectively, for each of the *n* = 1000 subjects with varying values of 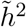. The true conditional heritabilities are all within 1 empirical standard deviation of the average estimates.

#### 3.1.3 Estimation of conditional heritability standard errors

We also calculate estimates for the conditional heritability standard errors by taking the square-root of Equation (12) in each replication. This is done for the simulation setting where we take the marginal heritability to be 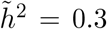. We then calculate an average of this for each subject and compare to the Monte Carlo standard errors calculated from the 100 replications. We can see that the average standard error estimates using Equation (12) correlate with the Monte Carlo standard errors as shown in Figure 3.2a. There seems to be a bias upwards, but it is preferable for practical purposes to over-estimate standard errors than underestimate them. That said, as illustrated in Figure 3.2b, the over-estimation is not too high, with the highest bias being under 0.008 which is small considering that the conditional heritability ranges from 0.23 to 0.38. Moreover, there are only three average estimated standard errors that are lower than the Monte Carlo standard errors, and these only seem to be slightly underestimated.

**Figure 3.2:**
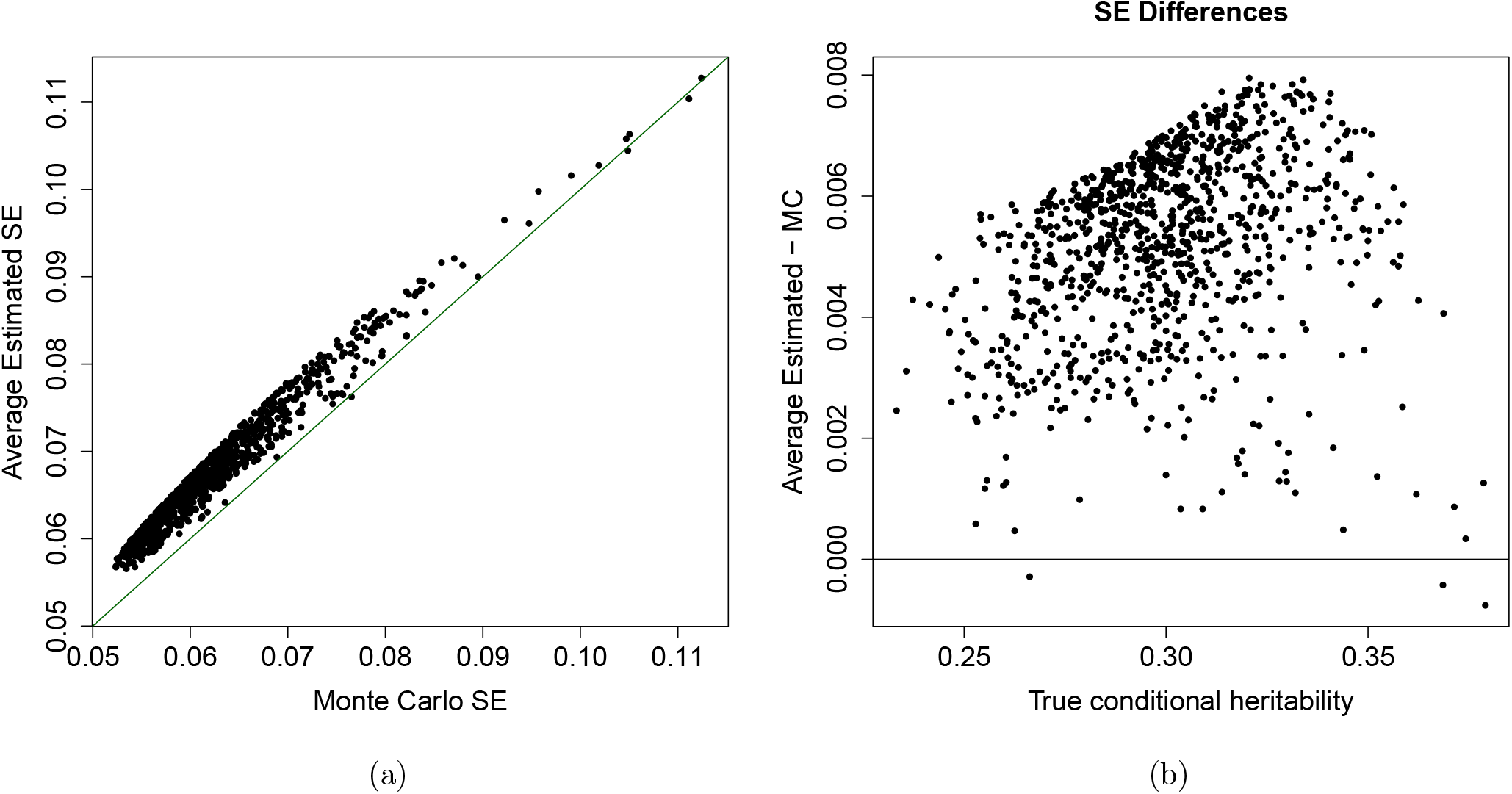
(a) Comparison of Monte Carlo standard errors and average estimated standard errors using Equation (12) for the simulation study from Section 3.1.2. The two are very close to each other, with the average estimated standard errors tending to be only slightly higher. (b) Comparison of the bias of the estimated standard errors across differing values of true conditional heritability for the simulation study from Section 3.1.2. The highest bias is under 0.008, which is small considering that the true conditional heritability ranges from 0.23 to 0.38.

#### 3.1.4 Additional simulation studies

We have mentioned previously that covariate-adjusted GWASH and covariate-adjusted LDSC perform similarly in simulations. The results from this simulation are summarized in Table F.1 in Appendix F. Covariate-adjusted GWASH and LDSC-1 result in very similar estimates for marginal heritability with a mean estimate of 0.283 and 0.284. Both mean estimates are within one standard deviation of the true simulated marginal heritability of 0.3. Conversely, the naive framework presented in Table 1 also underestimates the marginal heritability when using LDSC. This coincides with a trend described in [18]. These similarities between covariate-adjusted LDSC and covariate-adjusted GWASH are to be expected since the authors of [2] show that GWASH and LDSC-1 share similar asymptotic properties.

Furthermore, to test the robustness of the marginal and conditional heritability estimation frameworks we conducted a comprehensive simulation study where we also varied more parameters such as the number of covariates, the correlation between SNPs, the effect of ancestry on SNPs and the effect of ancestry on the phenotype. In all cases the covariate-adjusted GWASH our conditional estimation framework provided unbiased estimates of the target heritabilities. More details about this comprehensive simulation can be found in Appendix F.

Moreover, we also conduct a simulation study to analyze the effect of using principal components instead of ancestry proportions. In that study we find that our covariate-adjusted GWASH and conditional heritability framework alsolead to unbiased estimates of the target heritability as long as enough principal components are used to capture the number of underlying ancestries in the dataset. More details about this can also be found in Appendix F.

Finally, in Appendix G we present supplementary simulations small-scale simulations. Although these focus on a two population case and use a different simulation framework, our framework for estimating covariate-adjusted marginal and conditional heritabilities is still shown to be valid. Moreover, even though these simulations are less comprehensive in comparison to what those we have presented above, they also illustrate a specific scenario with admixed individuals in which *t*(***s***) and Var(*y*|***s***) are non-linear in ***s*** and use GAMs used to adequately estimate conditional heritabilities.

All of these simulations demonstrate the robustness of our heritability estimation methods to a variety of scenarios.

### 3.2 Application to IQ scores in the ABCD dataset

#### 3.2.1 Data description

We were interested in applying the marginal and conditional heritability estimation procedure on an ancestrally diverse dataset. For this, we used data from the Adolescent Brain Cognitive Development^SM^ Study (ABCD) (https://abcdstudy.org/about/). Details about the genetic data that we accessed can be found in [13]. We restricted ourselves to autosomal SNPs that were also present in a GWAS analysis carried out by Sniekers et al. for IQ-scores [36] because we were interested in comparing our results. We used PLINK 1.9 [4] [30] [28] to conduct quality control steps described by [21] and randomly selected 750,000 of these SNPs to continue to use for further analysis leading to a final dataset with *n* = 6623 individuals (consisting of 3148 female participants and 3475 male participants) and *m* = 750, 000 SNPs. This random selection of *m* = 750, 000 SNPs was done keeping in mind a comprehensive analysis with different traits carried out in [33] that showed that GWASH heritability estimates tend to converge after *m* = 750, 000 SNPs which can save computational time and resources. A full description about the analysis used to obtain the individuals and SNPs for our analysis can be found in Appendix H.

Recall that for our modelling framework we are using the underlying model described in Equation (1). For our data application our phenotype *y* corresponds to an individual’s IQ-score and the vector ***x*** corresponds to an individual’s SNP values for the 750,000 SNPs kept in the data analysis. Recall that ***s*** represents a vector with covariates of interest that can affect either the phenotype or genotype distribution. For this data analysis there are fifteen covariates of interest captured in ***s***: an intercept term, age, sex (taking a value of 0 for females and 1 for males), two indicators for household income (one variable representing household income greater than or equal to $50K but less than $100K, and another variable representing household income greater than or equal to $100K) and 10 principal components that had been previously calculated on a larger version of our dataset using PC-Air [6]. Principal components act as proxy variables that capture underlying ancestry, as described in [19], [20], [23] and [27]. We note that to avoid issues with multicollinearity we do not include the variable representing household income less than $50K since it can be inferred from the values of the other income variables. For IQ scores we used the crystallized cognition composite scores described in [16].

### 3.3 Justification of principal components for ancestry covariates

It is worth mentioning that the ABCD dataset also has other variables that are associated with ancestry - namely self-reported race/ethnicity (with 3719 participants reported as “White”, 1317 as “Hispanic”, 807 as “Black”, 95 as “Asian”, and 685 as “Other”) and proportion of European, American, East Asian and African ancestry calculated using “fastStructure” [32]. Using principal components as the proxies for ancestry has advantages over the self-reported race/ethnicity variables and the ancestry proportions. For example the article that introduces cov-LDSC [18] has already shown that regressing out principal components helps obtain estimates of adjusted marginal heritability for admixed populations and it recommends using 10 of these in practical settings. We only have estimates for four ancestry proportions and there might be more ancestral variability that might need to be captured (e.g. differences in individuals from North European and South European ancestry). Using principal components might capture these more granular differences.

Furthermore, self-reported race/ethnicity does not accurately reflect complete genetic ancestry. For example, one interesting observation from Figures H.20 and H.21 is that there is a fair amount of admixture within individuals self-reported as Asian since we see several individuals with more than 40 percent European ancestry in this category.

Moreover, historically described admixture within subjects self-reported as Hispanic can be observed with individuals in this category having a wide range of European and American ancestry proportions (Figures H.20 and H.22), as well as in individuals self-reported as Black who have a large range of African and European ancestry proportions (Figures H.19 and H.20).

Finally, individuals self-reported as “Other” have a scattered range and combination of ancestry proportions, but most seem to have at least 40 percent European ancestry (Figure H.20). That said, although self-reported race/ethnicity will not be used in the covariate vector ***s***, it can help summarize findings and relationships between variables. For example, as illustrated in Figure H.23, there is confounding between household income and self-reported race/ethnicity with certain groups reporting a higher household income than others. We shall also use this variable for graphical summaries for results obtained from the application of our conditional heritability framework. More details regarding our covariate assessment can be found in Appendix H.

#### 3.3.1 GWAS summary statistics and LD-scores

To obtain GWAS summary statistics for implementing the marginal heritability estimation methods, we ran PLINK 2.0 [29] to conduct a GWAS for the IQ scores with our subset of 750,000 SNPs while covarying for the fifteen covariates of interest defined in Section 3.2.1. We also downloaded the GWAS summary statistics provided by Sniekers et al. [36] for comparative purposes. To calculate adjusted and unadjusted LD-scores, we first took **X** to be the matrix of the 750,000 SNP values for the subjects in the ABCD dataset and obtained 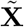 as discussed in Section 2.5 by residualizing on the 15 covariates of interest. We calculated the unadjusted LD-scores using the approximation from Equation (A.2) taking 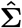 to be the sample correlation matrix obtained from **X** and *Q*_*j*_ to be the set of all SNPs that are in the same chromosome as SNP *j*. We also used this equation to calculate the adjusted LD-scores, but setting 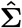 to be the sample correlation matrix obtained from 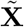 instead of **X**.

#### 3.3.2 Marginal heritability

We estimated covariate-adjusted marginal heritability using covariate-adjusted GWASH and covariate-adjusted LDSC-1 and contrasted the result with the naive approach described earlier in Section 3.1.1. We also repeated this process using GWAS summary statistics from the Sniekers et al. analysis [36].

Naive LDSC estimates for marginal heritability were obtained by running LDSC-1 with the unadjusted LD-scores and GWAS summary statistics from above, whereas cov-LDSC estimates were obtained by using the adjusted LD-scores instead. As mentioned previously, naive estimates are the most likely to be computed in practice if no further adjustment is made. The software provided by [3] was used for both of these LDSC runs after the LD-scores were calculated. We also made use of the LD-scores and GWAS summary statistics to obtain naive and covariate-adjusted GWASH estimates. Standard errors for the GWASH estimates were obtained as described by Equation (A.4) and for the LDSC-1 estimates were obtained from the output generated by the software pertaining to [3]. Full details are described in Appendix H. We summarize our results in Table 2.

**Table 2:** Covariate adjusted and naive marginal heritability estimates for the ABCD and Sniekers et al. IQ data. GWASH and LDSC-1 give similar estimates, but the estimated heritability is higher in the ABCD dataset than the Sniekers et al. dataset.

|  | ABCD |  | Sniekers et al. |  |
| --- | --- | --- | --- | --- |
|  | Adjusted | Naive | Adjusted | Naive |
| GWASH | 0.295<br>(0.052) | 0.070<br>(0.034) | 0.165<br>(0.007) | 0.039<br>(0.006) |
| LDSC-1 | 0.302<br>(0.049) | 0.071<br>(0.012) | 0.187<br>(0.008) | 0.049<br>(0.002) |

As can be seen from Table 2, the GWASH and LDSC-1 estimates are similar to each other. In the literature, the heritability of intelligence is estimated to be between 25% to 50%, with twin studies estimating it closer to 50% and SNP-heritability studies closer to 25% [26]. Our covariate-adjusted marginal heritability with the ABCD dataset is within this ballpark, whereas the corresponding Sniekers et al. estimate is below this. This is expected because Sniekers et al. acknowledge that heritability estimated from their study is understimated [36], and the crystalized metric used to measure intelligence for the ABCD dataset has also been previously shown to be strongly associated with polygenic predictors [16].

One important characteristic of the GWASH estimator is its ability to summarize the correlation between SNPs in a single parameter estimate 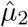. We obtained an “unadjusted” 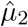 by plugging in the unadjusted LD-scores into the approximation from Equation (A.7). Analogously, an “adjusted” 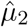 was obtained by plugging in the adjusted LD-scores into this equation. The resulting unadjusted and adjusted 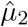 were 68.27 and 16.29 respectively. The difference between the unadjusted and adjusted 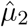 can be attributed to the residualization on covariates removing correlation between SNPs that is caused by underlying population structure and admixture in the sample. In the case of the ABCD dataset, this inflated correlation between SNPs leads to an underestimated heritability when using the naive framework, similar to the simulation study results presented in Section 3.1.1.

Although we use the same adjusted and unadjusted LD-scores for the ABCD and Sniekers et al. dataset to generate Table 2 for comparative purposes, it is worth keeping in mind that these datasets come from distinct population types with the ABCD dataset being more heterogenous and admixed, and so the underlying SNP correlation and LD-scores could differ between the two. Therefore we would caution about the interpretation of the naive heritabilities presented in the table for the Sniekers et al. dataset. In practice, if we had access to covariates and genotypes from the Sniekers et al. study we would expect the covariate-adjusted and naive frameworks to give similar estimates for the Sniekers dataset due to it being more ancestrally homogenous. That said, Sniekers et al. [36] estimate the SNP-heritability for their dataset using LDSC and obtain a heritability of 0.20 (S.E.= 0.01) with 12,104,294 SNPs which is within two standard errors of the estimated heritability of 0.187 that we obtain from using the adjusted LD-scores from the ABCD dataset restricting to 750,000 SNPs. Furthermore, when we use an alternative 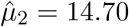 corresponding to 750,000 SNPs from the 1000 Genomes European sample [7] we obtain an estimated GWASH heritability of 0.183 for the Sniekers et al. dataset which is closer to the covariate-adjusted heritabilities in Table 2. Details to obtain this recalibrated 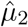 can be found in Appendix I.

#### 3.3.3 Conditional heritability

We estimated conditional heritabilities as described in Section 2.5. Since covariate adjusted GWASH and LDSC-1 give similar estimates for marginal heritability (Table 2), we used covariate adjusted GWASH as the estimate for 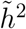. We tested the sensitivity of our approach to using using GAM instead of OLS for obtaining the estimated trace parameter 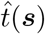 and variance parameter 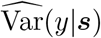. Both types of models were fit on the fifteen covariates of interest defined above. This was done using the “gam” function with default parameters from package “mcgv” in R version 4.0.2 (details can be found in [43] [44]) and the “lm” function in R version 4.0.2. We proceeded to use OLS estimates for these parameters since these make use of a more parsimonious modeling framework and generally coincide with GAM estimates too. More details about this discernment can be found in Appendix H.

Since our covariates included principal components, which are continuous variables that differ for each individual, we obtained conditional heritability estimates for each subject in our dataset. There was no discernible difference in the distribution of the estimated conditional heritabilities obtained when comparing male and female subjects. The distribution of estimates for 1−*h*^2^(***s***), *τ* ^2^(***s***) and *σ*^2^(***s***) separated out by income and ethnicity are illustrated in Figure 3.3. The estimates for parameters *τ* ^2^(***s***) and *σ*^2^(***s***) are obtained as described in Section 2.6. We decided to plot the extrinsicalities estimated as 1 − *ĥ*^2^(***s***) instead of the estimated heritabilities 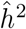(***s***) since non-genetic factors are more pertinent to the differences in subjects in our dataset. This is because, as shown in Figure 3.3, we observe an inverse trend between the estimated non-genetic variance *σ*^2^(***s***) and income, which seems to capture most of the differences between individuals. This matches the trend of a previous twin study which showed that the heritability of intelligence scores can change due to differing extrinsic variances in different socio-economic groups [39], as well as a recent study with the ABCD dataset that studied the SNP-heritability of cortical structure and cognitive ability [25].

**Figure 3.3:**
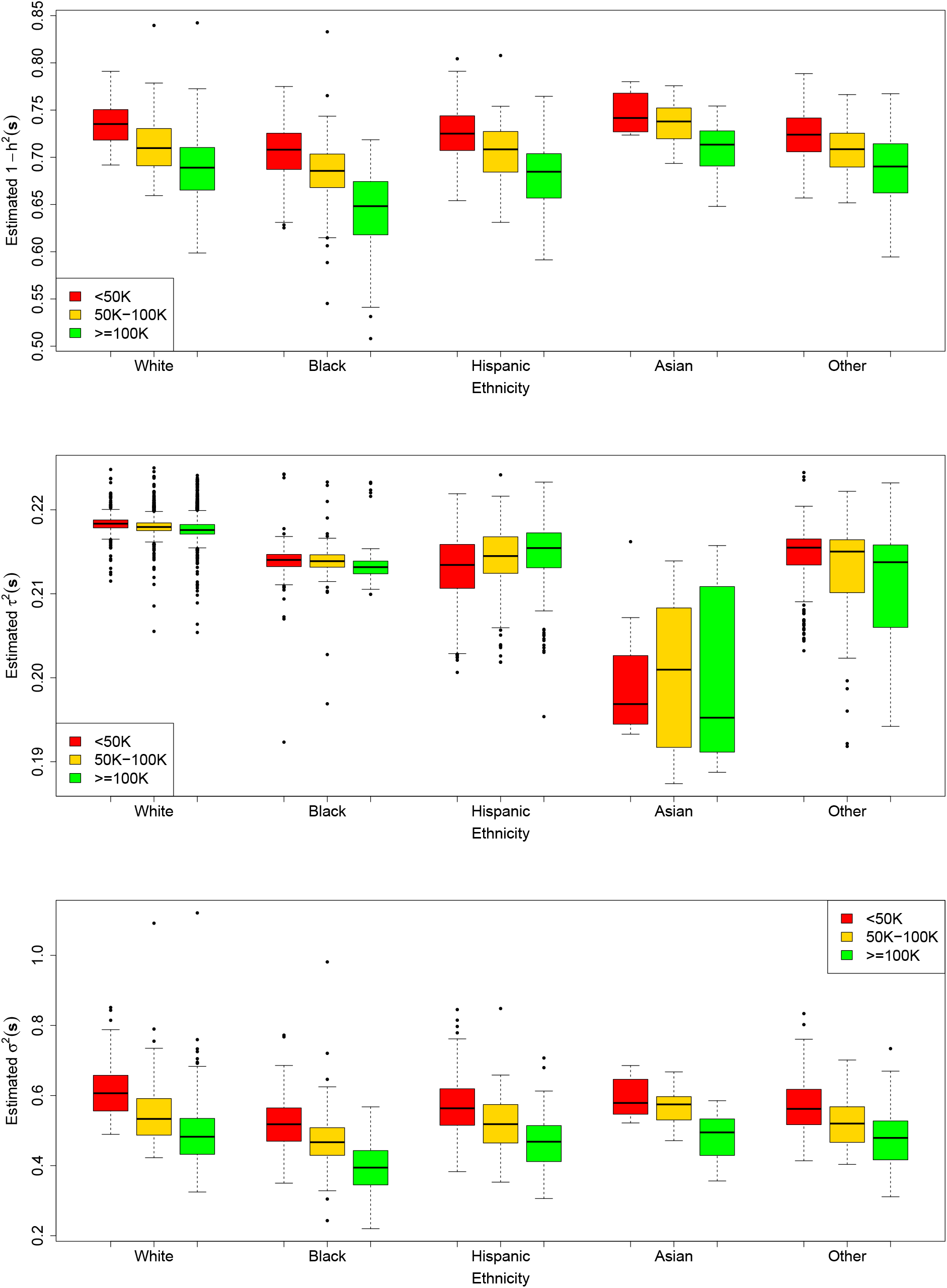
Boxplots of estimated extrinsicalities 1 − *h*^2^(***s***), *τ* ^2^(***s***) and *σ*^2^(***s***) for IQ scores in the ABCD dataset conditional on the fifteen covariates from Section 3.2.1. We observe an inverse trend between the non-genetic variance *σ*^2^(***s***) and income, which drives the same trend for the estimated extrinsicality 1 − *h*^2^(***s***). Self-reported race/ethnicity are used to display different sub-populations.

To further assess the precision of our conditional heritability estimates, we extracted the subjects whose extrinsicalities corresponded to the median for each income-ethnicity group and plotted their corresponding estimates for extrinsicalities and standard errors using the square-root of Equation (E.1). This corresponds to Figure 3.4a. We also plotted extrinsicalities plus/minus one estimated standard error for each subject in the ABCD data analysis in a scatterplot as shown in Figure 3.4b. From these figures, we notice that that the standard errors are highest when the estimated extrinsicality is close to 0.5 and then decrease with increasing extrinsicality. This is similar to the behaviour of the GWASH estimator for homogenous samples [33]. Moreover, the estimated standard errors seem to be generally higher in the Asian subgroup and lowest in the White subgroup, which can be explained by the difference in number of subjects for each of these (Figure H.23). Although there is an overlap in the standard error bars for the median conditional extrinsicalities for each income-ethnicity group in Figure 3.4a, when looking at all subjects in Figure 3.4b we notice that lower extrinsicality estimates differ by more than one standard error than higher estimates. For example, the highest conditional extrinsicality estimate is 0.84 (S.E. = 0.05) whereas the lowest one is 0.51 (S.E. = 0.14). These results suggest that a single estimate of marginal heritability and its standard error cannot capture the full behavior within our sample. Instead, conditional heritability estimates and their standard errors vary among subjects depending on their particular values of the covariates.

**Figure 3.4:**
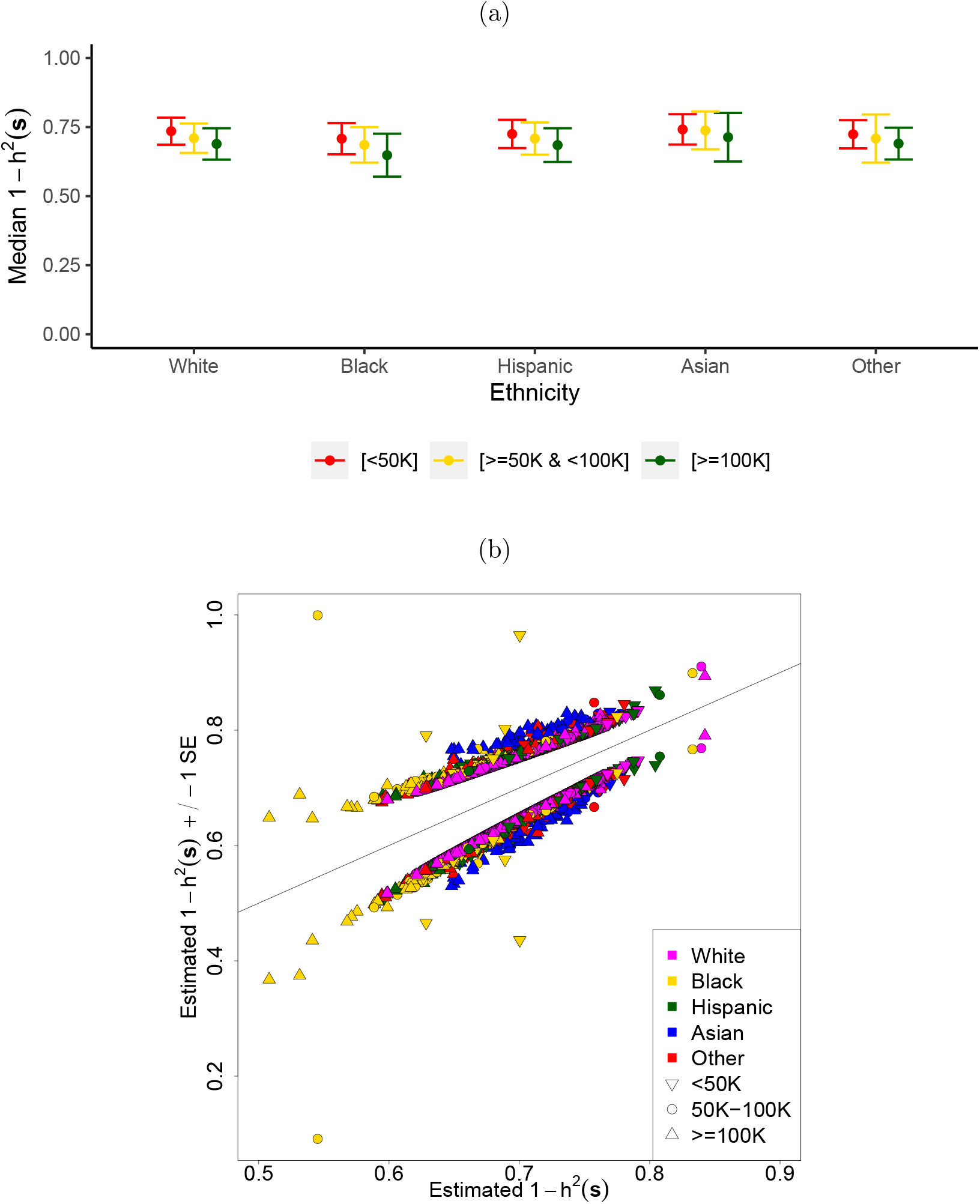
(a) Median estimated extrinsicalities for each income-ethnicity group along with their corresponding estimated standard errors. We observe a similar trend as that illustrated in Figure 3.3. (b) Estimated extrinsicalities plus/minus one estimated standard error for each subject in the ABCD data analysis (vertical axis) against the estimated extrinsicalities themselves (horizontal axis). We observe differences in estimated extrinsicalities and standard errors at an individual level.

Additional details and plots for this section can be found in Appendices H and J.

## 4 Discussion

### 4.1 Summary

In this work, we developed new theory and methods for covariate-adjusted marginal heritability, but also proposed an estimation procedure for conditional SNP-heritability. Our conditional heritability estimation procedure is a novel approach to the best of our knowledge, and can be used to estimate heritabilities for the different covariate combinations of interest in a sample, as well as their corresponding standard errors. We carried out a comprehensive simulation study where we tested our procedures for estimating covariate adjusted marginal and conditional heritabilities, which showed that our estimates were close to their corresponding values. We also derived a procedure to estimate standard errors for our conditional heritability estimates and showed through simulations that these were only slightly higher than their respective Monte Carlo estimates. Finally we ran a study on the ABCD study for estimating the conditional and marginal heritability of IQ scores where we observed a decreasing trend in conditional extrinsicalities with increasing household income which coincides with other studies in the literature.

Our proposed estimation approach can be a valuable tool for understanding differences in risks across different populations, as our conditional heritability estimates can also be conditioned on other covariates such as lifestyle factors to understand how genetics can come into play in people with different lifestyles. As datasets become increasingly diverse in nature, new methods need to be developed in order to analyze these. Our work here improves on traditional methodology that was originally designed to analyze homogeneous datasets and allows us to leverage the heterogeneity of our data to obtain new insights. Below we discuss some additional considerations regarding our methods and application.

### 4.2 Extrapolation of simulation study results

In order to carry out simulations we had to define an underlying SNP-correlation parameter as detailed in Appendix F. The value of this parameter could vary depending on the population and how many SNPs were sequenced, but our simulations in Appendix F do not show much change across different values of this parameter.

Furthermore, there is a “breakdown” point where the naive methods stop working and the covariate adjusted estimators become necessary to estimate marginal heritability. This is explored in the simulations presented in Appendix F. It is important to note that for different values of the simulation parameters, the breakdown point for the naive framework could be different. Nonetheless, in practice, the breakdown would be observed if the naive framework and the covariate adjusted framework resulted in considerably different estimates of marginal heritability, which happens to be the case for the ABCD dataset. In such cases our simulations suggest to adopt the covariate adjusted framework.

For the estimation of marginal heritability and consequentially conditional heritability, we need to make sure we have residualized enough ancestry-related covariates from the SNPs as shown in Appendix F. Moreover, the resulting residualized SNPs still need to satisfy the assumptions required for the consistency of the heritability estimator of interest used to calculate marginal heritability. Deviations such as the presence of family-related individuals and large individual SNP effects, among other factors, can lead to inaccurate estimates of marginal and conditional SNP heritabilities.

### 4.3 Estimating *t*(s) and Var(*y*|s)

In our implementation, the quantities *t*(**s**) and Var(*y*|**s**) were estimated using OLS and GAM. When applying our methods in practice we would recommend testing out both a GAM and OLS model for estimating *t*(**s**) and Var(*y*|**s**) since there can be scenarios where a GAM is more appropriate as shown in Appendix G. That said, at least for our data analysis of IQ scores for the ABCD dataset, both models gave similar results and thus we opted for OLS models due to parsimony. Other models apart from GAM and OLS can also be used if deemed appropriate to capture the behavior of these parameters.

### 4.4 Residualization

Residualizing SNPs on covariates for our methods can be a computationally expensive process. The cov-LDSC paper [18] suggests a slightly less computationally expensive way to do so if the principal components obtained are orthogonal and no other covariates affect the distribution of SNPs. In our case we could not follow this procedure since the principal components for the ABCD dataset had been previously calculated on a larger sample with PC-Air [6], and were not orthogonal for the sub-sample used.

Moreover, our framework can be applied to situations where we have a sample from a homogenous ancestry but differing non-genetic covariates such as socioeconomic variables. In such cases it is only necessary to pre-residualize the phenotype and not the genotype in order to estimate the marginal heritability 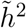.

### 4.5 A note on the use of standard errors

Although we have developed a framework for estimating standard errors for our conditional heritability framework, these should only be used for exploratory purposes since we have not developed any theory for the asymptotic distribution of our estimates. Developing a hypothesis testing framework for conditional heritability estimates is left for future work.

### 4.6 Obtaining marginal heritabilities for a new population

Recall the definition of covariate adjusted marginal heritability from Equation (2). If we know or have an approximation for the distribution of covariates in a new population of interest, we can plug in the corresponding probability density function *π*(***s***) along with the current sample estimates for *τ* ^2^(***s***) and Var(*y*|***s***) to obtain a new marginal heritability. We shall leave implementing standard errors for this situation for future work.

### 4.7 Implications for polygenic risk scores

Several studies such as [22] have drawn attention to the fact that the accuracy of polygenic risk scores decreases in non-European cohorts. Recently, [10] showed that the accuracy of polygenic scores can vary accross the continuum of ancestry compositions in all populations, even within European cohorts conventionally thought of as homogeneous. Though an increased diversity of participants is crucial to address these discrepancies in accuracy [22], applying a conditional heritability estimation framework can further help assess if variations in the performance of polygenic risk scores can be attributed to differences in genetic and non-genetic variance accross the continuum.

### 4.8 Comparison to Norbom et al. [25]

Although our conditional heritability estimation framework shows that non-genetic extrinsic variance decreases with higher socioeconomic status, similar to Norbom et al. [25], important differences exist in our methods and results.

Norbom et al. include related individuals (*n* = 9080), whereas we follow the conventional GWAS approach [21] and remove related participants, yielding *n* = 6623 subjects. They analyze total cognition composite scores, while we focus on crystallized cognition scores [1, 16, 17]. Socioeconomic indicators also differ: they use a composite of income, education, and neighborhood deprivation, while we use categorical household income. Methodologically, they use an underlying mixed-effects model for the phenotype, whereas we employ linear regression. Moreover, our heritability estimates are a function of fifteen covariates (Section 3.2.1), whereas in their case they are only a function of socioeconomic status.

All the above can lead to differences in heritability definitions and estimates. While both studies show reduced non-genetic variance with higher socioeconomic status, Norbom et al. additionally report that the genetic contribution to heritability increases with socioeconomic status, whereas we do not observe this trend in our estimated *τ* ^2^(***s***) in Figure 3.3, possibly due to the reasons described above.

## 5 Data and code availability

The ABCD data repository grows and changes over time. The ABCD data used in this report came from 10.15154/z563-zd24 for SNPs, and 10.15154/1519007 and 10.15154/1523041 for other variables used in the analysis. DOIs can be found at http://dx.doi.org/10.15154/z563-zd24, http://dx.doi.org/10.15154/1519007 and http://dx.doi.org/10.15154/1523041.

Links to additional data and code will be made available at the time of journal publication.

## 6 Author contributions

A.N.S.S: Conceptualization, data curation, formal analysis, methodology, software, validation, visualization, writing and editing.

D.A: Conceptualization, methodology, writing, editing and supervision.

A.S: Conceptualization, methodology, funding acquisition, project administration, writing, editing, supervision.

## 7 Acknowledgments

We would like to acknowledge UCSD Professors Chun Chieh Fan, Wesley Thompson, Tiffany Amariuta, Steven Edland, Karen Messer and Xinlian Zhang for the insights they have provided for this work.

Anubhav Nikunj Singh Sachan would like to acknowledge the Mexican Consejo Nacional de Ciencia y Tecnología (CONACYT) for providing funding for his PhD studies.

This work was funded in part by the NIMH grant R01MH128923.

Data used in the preparation of this article were obtained from the Adolescent Brain Cognitive Development^SM^ (ABCD) Study (https://abcdstudy.org), held in the NIMH Data Archive (NDA). This is a multisite, longitudinal study designed to recruit more than 10,000 children age 9-10 and follow them over 10 years into early adulthood. The ABCD Study^®^ is supported by the National Institutes of Health and additional federal partners under award numbers U01DA041048, U01DA050989, U01DA051016, U01DA041022, U01DA051018, U01DA051037, U01DA050987, U01DA041174, U01DA041106, U01DA041117, U01DA041028, U01DA041134, U01DA050988, U01DA051039, U01DA041156, U01DA041025, U01DA041120, U01DA051038, U01DA041148, U01DA041093, U01DA041089, U24DA041123, U24DA041147. A full list of supporters is available at https://abcdstudy.org/federal-partners.html. A listing of participating sites and a complete listing of the study investigators can be found at https://abcdstudy.org/consortium_members/. ABCD consortium investigators designed and implemented the study and/or provided data but did not necessarily participate in the analysis or writing of this report. This manuscript reflects the views of the authors and may not reflect the opinions or views of the NIH or ABCD consortium investigators.

## 9 Declaration of generative AI and AI-assisted technologies in the writing process

During the preparation of this work the authors used ChatGPT in order to improve the readability of some sections. After using this tool, the authors reviewed and edited the content as needed and take full responsibility for the content of the publication.

## A Background

### A.1 SNP-heritability

Assume that ***x*** is a random vector of length *m* that represents the sequenced SNPs for a particular person with covariance Cov(***x***) = **Σ**, and that *y* is a continuous trait or phenotype of interest. Assume that the relationship between *y* and ***x*** can be modelled as a linear model *y* = *α* + ***β***^*T*^ ***x*** + *ϵ* where *ϵ* is a normal random variable with mean *E*(*ϵ*|***x***) = 0 and variance *E*(*ϵ*^2^|***x***) = *σ*^2^. We can define SNP-heritability as:

#### Definition 3

(SNP-heritability). 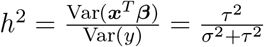 *where τ* ^2^ = ***β***^*T*^ **Σ*β***.

This definition is intuitive because the variance of *y*, Var(*y*) can be decomposed as Var(*y*) = *σ*^2^ + *τ* ^2^ where *τ* ^2^ can be seen to represent the portion of variance arising from the randomness of the SNPs and *σ*^2^ can be seen to represent the portion of the variance arising from other unobserved factors.

We shall assume from this point that ***x*** and *y* are mean-centered and scaled.

### A.2 Linkage disequilibrium

Linkage disequilibrium (LD) can be generically be defined as “the nonrandom association of alleles of different loci” [35]. The statistical interpretation of this is that high LD refers to two SNPs having high dependence with each other and low LD refers to two SNPs having low dependence with each other. There are various ways one can define “dependence between SNPs” and [35] reviews various of these. For our purposes we will assume that LD refers to “linear dependence” and thus quantify it with the Pearson correlation matrix between SNPs which we will call LD-matrix. We note that there is a difference between the population LD-matrix which we will denote with **Σ** and the estimated or “in-sample” LD-matrix which we will denote with 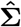. This concept of linkage disequilibrium and the LD-matrix plays a key role in heritability estimators, as well as when accounting for population stratification present in ancestrally diverse datasets.

### A.3 LD-score regression

One commonly used method for estimating SNP-heritability in the setting where *m* >> *n* is LD-score regression (LDSC) [3]. One reason for its popularity is the ability to apply this method starting out with the following summary statistics:

1. The p-values/*t*-statistics from a GWAS.
2. An estimated LD-matrix. One can also use pre-computed LD-scores from a reference panel that is similar to the sample in order to reduce computational time.

LD-scores captures the amount of linkage disequilibrium that a SNP has with all others, and are calculated from an in-sample LD-matrix 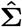 as:

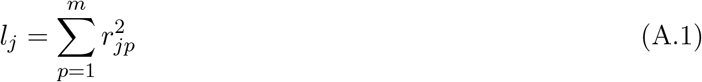

where 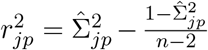 and 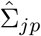 corresponds to the in-sample correlation between the *j*-th and *p*-th SNP. To improve the computational efficiency of calculating LD-scores, it can be assumed that SNPs located on different chromosomes are independent from each-other, and get the approximation:

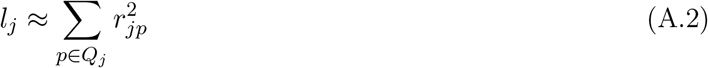

where *Q*_*j*_ is the set of all SNPs that are in the same chromosome as SNP *j*. By using this approximation, we only need to calculate the elements of 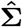 that correspond to SNPs in the same chromosome. Moreover, SNPs that are located far from each other within a certain chromosome can also be assumed to be independent, in which case *Q*_*j*_ represents the set of indices that are within a certain distance of SNP *j*. This distance can be pre-defined by the user, but in the absence of admixture [3] recommends using a 1 centimorgan window.

LD-score regression assumes each SNP effect *β*_*j*_ is a random variable drawn from a distribution with mean 0 and variance 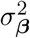, obtaining SNP-heritability as:

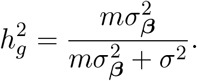

Let **X**_*j*_ represent the *j*-th column of SNP-matrix **X**. The estimation of 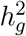 with LD-Score regression relies on the following claimed relationship between 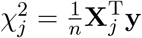 and the LD-score *l*_*j*_:

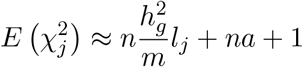

where *a* = 0 under assumptions of large *n* and no structure or relatedness between subjects.

The heritability 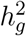 is estimated with a weighted regression of 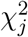 on *l*_*j*_ and its standard error is estimated with a block jackknife procedure. If *a* = 0 then the intercept term in the regression is constrained to 1.

More details about this derivation can be found in [3] and the follow-up [18]. It is worth remembering that we are assuming for all of these that the matrix of SNPs and vector phenotypes have been previously centered and scaled.

The *χ*^2^ statistics can be calculated from the *t*-statistics from a GWAS as follows:

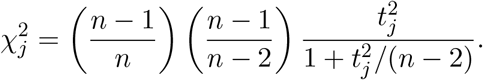

Note that for large enough *n*:

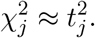

This approximation for 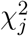 is implemented in the LD-score regression software.

### A.4 GWASH

Another way to estimate SNP-heritability is the GWAS heritability estimator (GWASH) proposed by Schwartzman et al [33]. This method is a modification of Dicker’s estimator with unestimable covariance [9] which allows one to use summary statistics from a GWAS similar to LD-score regression. It takes the following form:

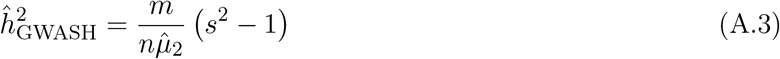

where:

- 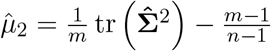
- 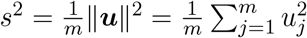
- 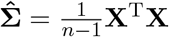
- 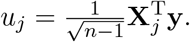

It is worth remembering again that we are assuming that the matrix of SNPs (**X**) and vector phenotypes (**y**) have been previously centered and scaled.

If one breaks down Equation (A.3) one can explain it is form in an intuitive manner. The quantity *s*^2^ represents the sum of squared sample correlations between each SNP and the outcome, and thus we would expect heritability to be larger if *s*^2^ is large. Moreover, 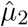 is a measure of the correlation between SNPs and is an estimator for *µ*_2_ = tr(**Σ**^2^). If there is higher correlation between SNPs, we would expect *s*^2^ to be inflated and so we need to divide by 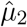 to account for this inflation. The reason for the expected inflation in *s*^2^ is because if a SNP *j* is not associated with the outcome of interest, but has high correlation with a SNP that is, then *u*_*j*_ will overestimate the association between SNP *j* and the outcome.

Schwartzman et al. [33] show that 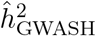 is a consistent estimator for *h*^2^ defined in Definition 3 under certain conditions as illustrated by the following assumptions and theorem:

#### Assumption 3.

*Suppose the variance components σ*^2^ *and τ* ^2^, *as well as the spectral moments m*_*k*_ = tr (**Σ**^*k*^) */m, k* = 1, …, 4, *are bounded. Let* 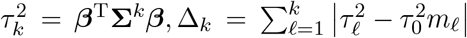, *and suppose* 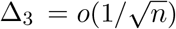.

#### Theorem 1.

*Under Assumption 3, as m and n get large such that m/n converges to a constant (which may be zero), the GWASH estimator* (A.3) *satisfies the central limit theorem* 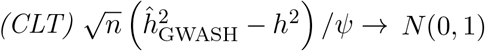, *where*

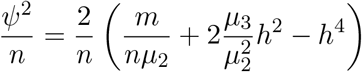

*and µ*_*k*_ = tr(**Σ**^*k*^).

Thus, Theorem 1 implies that GWASH is consistent for large *m* and *n*. It is also valid for both *m* > *n* and *m* < *n*.

The GWASH estimator is advantageous because it can be used to conduct inference. If *m* and *n* are large, this estimator is approximately normally distributed with mean *h*^2^ and variance defined in 1.

An estimate for the standard error of 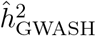 is 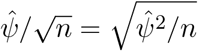, where

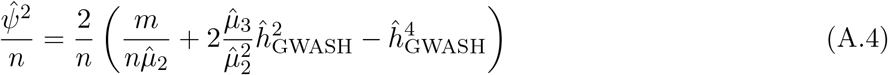

and 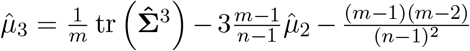 is an estimator of *µ*_3_, and 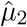 and 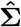 are taken to be the same as in Equation (A.3).

Using the asymptotic normality of 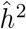 described in Theorem 1 one can construct an approximate two-sided confidence intervals for *h*^2^ of level 1 − *α* as 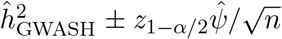 where *z*_1−*α*/2_ is the 1 − *α*/2 quantile for a standard normal distribution. For example, if one wanted an approximate two-sided 95% confidence intervals for *h*^2^ this would be of the form 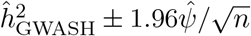.

Moreover, one can test the null hypothesis *H*_0_ : *h*^2^ = 0 against *H*_*A*_ : *h*^2^ *>* 0 using a Wald test statistic of the form 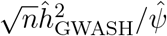 and rejecting the null hypothesis at level *α* if it exceeds *z*_1−*α*_.

Now we discuss how one can apply GWASH using similar summary statistics as LD score regression.

To do so, the following are needed:

1. The p-values/*t*-statistics from a GWAS.
2. An estimated LD-matrix. One can also use pre-computed estimates for 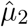 and 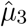 from a reference panel that is similar to the sample in order to reduce computational time.

If one has the *t*-statistics from a GWAS one can obtain the square of the correlation scores as

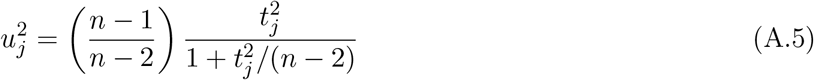

where *t*_*j*_ represents the *t*-statistic for SNP *j*. In case of just having p-values, one can map these to the corresponding *t*-statistic and then obtain the square of the correlation scores as above. Then these scores can be used to compute the *s*^2^ in A.3.

For the estimates of 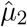 and 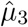 one can use the LD-matrix from a publicly available dataset such as the 1000 Genomes project [7] in case the studies in-sample LD-matrix is not available assuming the population the sample comes from and the sequenced SNPs are similar enough to that of the publicly available dataset. We also note that if we have access to LD-scores (*l*_1_, *l*_2_, …, *l*_*m*_) then it is easy to show that we can calculate 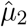 as:

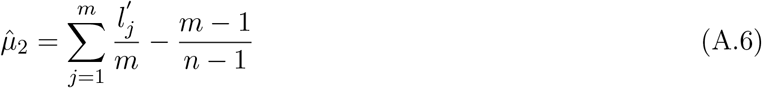

where 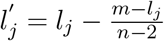.

Furthermore if the LD-scores are approximated using Equation (A.2) with *Q*_*j*_ representing all the SNPs in the same chromosome as SNP *j*, one can approximate 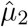 as follows:

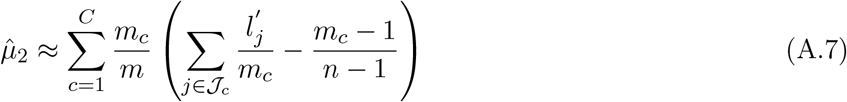

where 𝒥_*c*_ is the set of SNPs in the *c*-th chromosome, *m*_*c*_ is the number of SNPs in the *c*-th chromosome included in the analysis, and 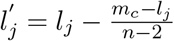 with *l*_*j*_ approximated using Equation (A.2),

Moreover, note that 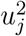 and 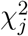 are asymptotically equivalent for large *n* since:

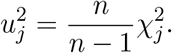

Although GWASH and LD-score regression with intercept 1 are derived under different frameworks, [33] shows that a stylized version of LD-score regression (which involves conducting a linear regression instead of a weighted regression) is asymptotically equivalent to GWASH. Moreover, applying GWASH and LD-score regression with intercept 1 to different datasets in [33] results in similar heritability estimates, and in a follow-up paper [2] it is further shown that GWASH and LD-score regression share similar properties and behaviors.

### A.5 Principal components

One issue present in ancestrally diverse datasets is population stratification. Although its definition can vary depending on the context one studies it in, for our purposes we will define population stratification as similarities in some of our sample’s genetics due to a common shared ancestry. This common shared ancestry can be an issue because it is often associated with hidden variables (cultural, social, environmental, etc.) that can have an impact on the trait of interest. The presence of diverse ancestries can also lead to LD-matrix structures that violate assumptions of commonly used heritability estimators.

In the context of GWAS studies, a common method used to control for population stratification involves using principal components. The usage of this method gained popularity after being described in [27]. For our simulation studies, these are calculated as follows.

Suppose as before that we start with a SNP matrix **X** where each column has been scaled and centered. We first define the in-sample kinship matrix 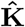 as:

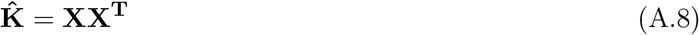

We note that this differs slightly from other definitions of the in-sample kinship matrix that might multiply it by a factor such as 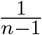. We then obtain the eigenvectors of this matrix 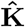 by applying singular value decomposition on **X**:

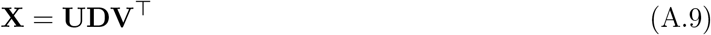

where **V** is an *m* × *n* matrix, **U** is an *n* × *n* matrix, and **D** is a diagonal matrix. The columns of **U** are the eigenvectors of 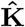 that we are interested in obtaining. We refer to the *u*_*ik*_ entry of the matrix **U** as the *k*-th principal component for individual *i*. Extensions to this such as PC-Air [6] address the presence of potential family structure.

## B Supplementary derivations for sections 2.2 and 2.5.4

In this section we will provide derivations to justify the use of 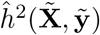 to estimate the adjusted marginal heritability 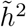 from Definition 1 in Section 2.2. Moreover, the first part of this section can also be used to justify our estimation procedure for the denominator in our proposed conditional heritability estimation framework in Section 2.5.4.

As before, we shall use the model described in Defintion 2. Let 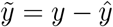 where *ŷ* = *Ê*[*y*|***s***]. If *Ê*[*y*|***s***] is estimated using an OLS framework, it will be unbiased and so *E*[*ŷ*|***s***] = *E*[*y*|***s***]. Thus:

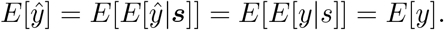

Therefore:

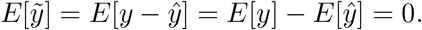

Moreover, 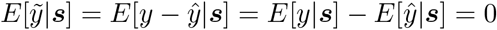. This implies:

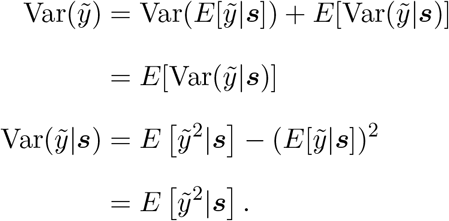

Under the OLS framework, for a large enough sample size *ŷ* ≈ *E*[*y*|***s***]. Thus 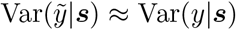 and so:

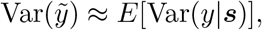

where the expectation is taken with respect to the distribution of ***s***. Note that this corresponds to the numerator of the covariate adjusted marginal heritability from Definition 1.

Now, let 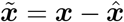 where 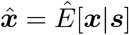. From this we obtain:

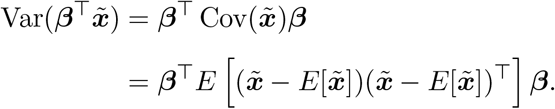

Using an analogous reasoning as before, 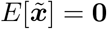 and so:

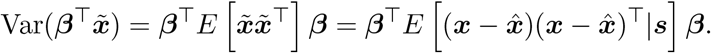

For large enough sample sizes 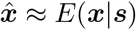 and so:

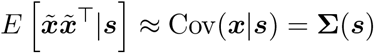

Plugging this back into the expression for 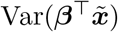:

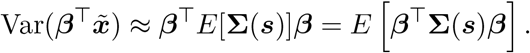

Since *τ* ^2^(***s***) = ***β***^⊤^**Σ**(***s***)***β*** we get:

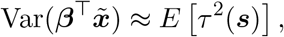

where the expectation is taken with respect to the distribution of ***s***.

Note that this corresponds to the numerator of the covariate adjusted marginal heritability from Definition 1.

Therefore the covariate adjusted marginal heritability 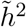 from Definition 1 can be estimated as 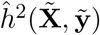. It is important to mention, that this requires the observed sample of phenotypes **y**, SNP-matrix **X** and covariate matrix **S** to be representative of a target population.

## C Using summary statistics for marginal heritability estimation

Recall that in the absence of covariates, one can use LD-scores to obtain 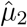 using Equation (A.6), and *t*-statistics from a GWAS to obtain correlation scores using Equation (A.5). These can then be plugged into Equation (A.3) to obtain the GWASH estimate for heritability. We can carry out a similar procedure to obtain a GWASH estimate for marginal heritability in the presence of covariates.

To do so, we first use Equation (A.6) to obtain an adjusted estimate for 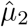 but use covariate-adjusted LD-scores instead of regular LD-scores. The covariate-adjusted LD-scores are calculated as in Equation (A.1) but using 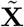 to obtain 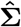 instead of **X**.

Next, if the GWAS accounts for covariates in the linear regressions that it fits, and one has access to the *t*-statistics from it, then one can obtain the square correlation scores with the following modified transformation:

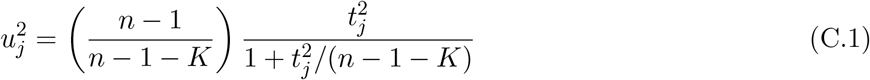

where *t*_*j*_ represents the *t*-statistic for SNP *j* and *K* represents the number of covariates including a possible intercept term. Once calculated, these correlation scores can be plugged into the GWASH estimator defined in Equation (A.3). We note that if the SNPs, phenotypes and covariates are centered, then this process implicitly adds an intercept term to the underlying model that needs to be accounted for in *K*. The justification for this comes from the Frisch-Waugh-Lovell theorem.

## D Derivation of the Conditional Heritability Estimator from Section 2.4

In this section we derive the conditional heritability estimator presented in Section 2.4.

Under Assumption 2, we can express Equation (4) as:

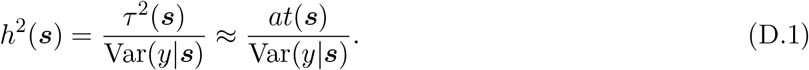

Also notice that under this same Assumption 2:

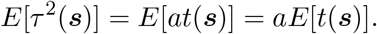

and so:

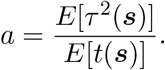

Furthermore, from the definition of covariate-adjusted marginal heritability:

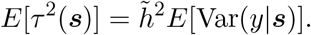

Thus, we can express *a* as:

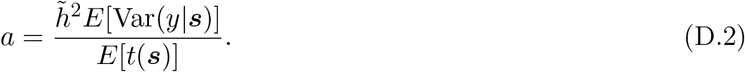

Plugging in Equation (D.2) into Equation (D.1), we obtain the following approximation:

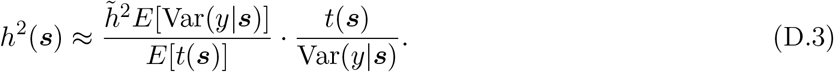

where the expectations are taken with respect to the distribution of ***s***.

Therefore, we can express conditional heritability as a function of five parameters: 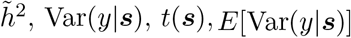 and *E*[*t*(***s***)].

## E Estimating the variance of conditional heritability estimates

We shall describe here a framework using the Delta Method and bootstrapping to estimate the variance of the conditional heritability estimator *ĥ*^2^(***s***). This variance will also be a function of covariates ***s***.

First, notice that the conditional heritability approximation from Equation (5) can be expressed as:

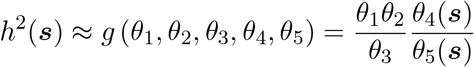

where:

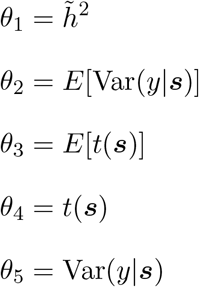

The derivatives with respect to each parameter are as follows:

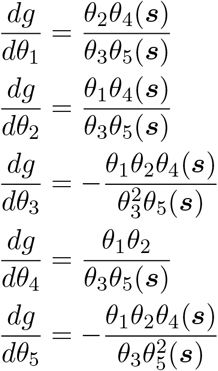

Now, we plugin our previously described parameter estimates to obtain the condtional heritability estimator:

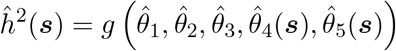

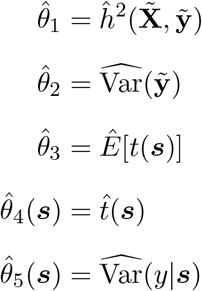

Let 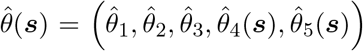. Assuming that the parameter estimates are approximately independent from each other, we apply the Delta Method to obtain the following estimate for the variance of the estimator:

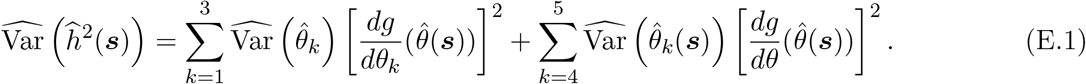

Note that the first three parameter estimates are not a function of ***s***, but the fourth and fifth are. If using GWASH, 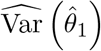 can be obtained using the result from Theorem 1 using 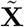 to obtain the covariance matrix to estimate the spectral moments and plugging in the covariate adjusted marginal heritability estimates instead of the regular heritability estimates. If using covariate adjusted LD-Score regression, 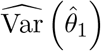 can be obtained through the software provided in [18] which uses a block jackknife estimation procedure [3].

In the case of the variance of 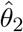 we can bootstrap 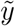 and recalculate its variance *B* times to create the bootstrap sample 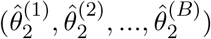 We then calculate the sample variance of this to obtain 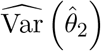.

For obtaining the variances of 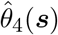 and 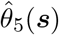 we can bootstrap our sample covariates and fit the corresponding models with these to obtain the set of fitted functionals 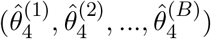 and 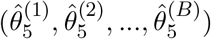.

To obtain 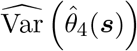 and 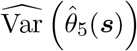 we evaluate these functionals at ***s*** and obtain the sample variance.

For example,

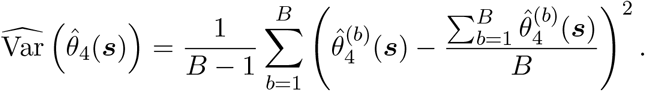

To obtain 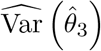 we also boostrap our covariates *B* times to obtain the corresponding bootstrap estimates 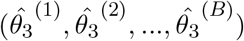. We then calculate the sample variance of this to obtain 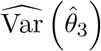.

## F Comprehensive simulation study

In this section we shall run a comprehensive simulation study to show that the estimated covariate-adjusted marginal and conditional heritability values match their theoretical values under our framework. We shall also compare our proposal to calculate standard errors of the conditional heritability estimates from the previous section to Monte Carlo standard error values, and show that these correlate with each other. All of these simulations demonstrate the robustness of our heritability estimation methods to a variety of scenarios.

### F.1 Simulation setup

Our simulation study uses mechanisms to generate ancestry proportions, SNPs based on transformed AR(1) processes, and phenotypes based on the model from Equation (1). These are detailed below.

#### F.1.1 Simulation of ancestry proportions

Let *K* be the number of underlying ancestries. To simulate a vector of ancestry proportions ***s*** for a subject, we generate *K* i.i.d. Uniform(0, 1) random variables (*q*_1_, *q*_2_, …, *q*_*K*_) and then set 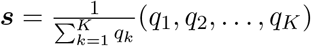. We note that the expected value for ***s*** simulated as such is *E*[***s***] = (1*/K*, 1*/K*, …, 1*/K*).

The ancestry proportions for each of the *n* subjects are generated using this mechanism before the start of the simulation study and kept fixed for each replication.

#### F.1.2 Simulation of AR(1) process

We simulate an AR(1) process with a predefined correlation *ρ* as follows:

- First generate a vector of i.i.d. standard Normal variables: 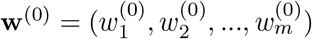.
- Next, transform **w**^(0)^ into **w** = (*w*_1_, *w*_2_, …, *w*_*m*_) by setting 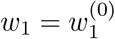 and 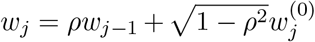 for *j* = 2, … *m*. It is simple to show that the resulting vector will have an AR(1) structure with E(*w*_*j*_) = 0, Var(*w*_*j*_) = 1 and Cor(*w*_*j*_, *w*_*j*−*l*_) = *ρ*^*l*^.

It is worth mentioning that the pre-residualizing ***x*** to obtain vector 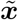 as described in Appendix B can be seen as a process to obtain a proxy for **w**.

#### F.1.3 Simulation of SNPs

To simulate correlated SNPs ***x*** = (*x*_1_, *x*_2_, …, *x*_*m*_) that have a distribution dependent on ancestry we generate an AR(1) process **w** as above and set 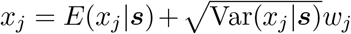 where *E*(*x*_*j*_|***s***) and Var(*x*_*j*_|***s***) represent the expectation and variance of SNP *x*_*j*_ conditional on a subject’s ancestry composition. This construction results in SNPs with Cor(*x*_*j*_, *x*_*j*−1_|***s***) = *ρ* and 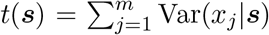. We detail the choice of *E*(*x*_*j*_|***s***) and Var(*x*_*j*_|***s***) below.

The conditional expectation *E*(*x*_*j*_|***s***) uses the fact that *“the expected allele frequency in the admixed population is the linear combination of the allele frequencies in the parental populations with weights determined by the sample admixture proportion”* [34] and is generated using the model:

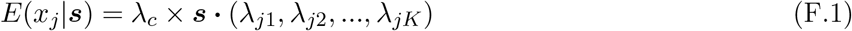

where × represents the multiplication between two scalars, · represents the vector dot product, *λ*_*j*1_, *λ*_*j*2_, …, *λ*_*jK*_ are drawn from standard normal distributions, and *λ*_*c*_ is a constant that controls the strength of ancestry on SNP means. Since different SNPs have different allele frequencies in real data, drawing random *λ*_*j*1_, *λ*_*j*2_, …, *λ*_*jK*_ once at the beginning of the simulation study setting allows us to mimic this characteristic. As an illustrative example, if our sample only consisted of European, East Asian and admixed individuals then the mean of the *j*-th SNP for an individual with 30 percent European ancestry and 70 percent East Asian ancestry would be *E*(*x*_*j*_|***s***) = 0.3*λ*_*c*_*λ*_*j*1_ + 0.7*λ*_*c*_*λ*_*j*2_ where *λ*_*c*_*λ*_*j*1_ and *λ*_*c*_*λ*_*j*2_ would correspond to the expected SNP value for someone with only European and only East Asian ancestry respectively.

To allow for a *t*(***s***) that has a linear relationship with ancestry composition so it can be estimated as described in Section 2.5, the conditional SNP variance Var(*x*_*j*_|***s***) is generated using the model:

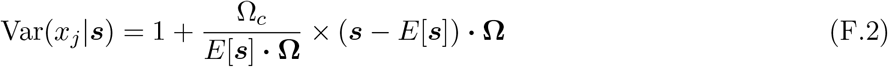

where *E*[***s***] = (1*/K*, 1*/K*, …, 1*/K*) as mentioned above, and **Ω** is taken to be **Ω** = (0, 1, 2, …, *K* − 1) which allows for a range of ancestry effects on SNP variance. The parameter Ω_*c*_ is a constant that can be varied to control the effect of ancestry on SNP variance, for example when Ω_*c*_ = 0 the SNP variance equals 1 regardless of ancestry. Note that plugging in different values for **Ω** which only differ by non-zero multiplicative constants will result in the same modelled conditional SNP variance. For example plugging in 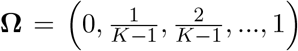 or **Ω** = (0, 1000, 2000, …, 1000 × (*K* − 1)) will result in the same value for Var(*x*_*j*_|***s***).

Moreover, this model ensures positive SNP variances for any Ω_*c*_ in [0, 1). To see this, note that since ***s*** represents ancestry proportions, all of its entries are non-negative not exceeding 1, and that the smallest entry of the defined **Ω** is 0 then inf 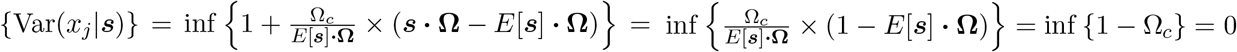 which occurs at ***s*** = (0, 1, …, 1) when Ω_*c*_ approaches 1.

This results in 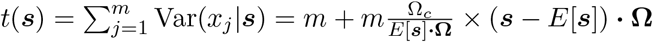 which is linear in ***s***.

#### F.1.4 Simulation of phenotype

To simulate *y* we pre-define the SNP effect sizes ***β*** and the marginal heritability of interest 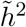. We then go through the following steps:

- Model *τ* ^2^(***s***) as:

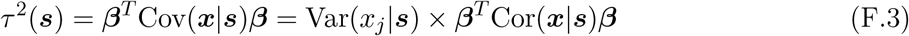

where Var(*x*_*j*_|***s***) is obtained from Equation (F.2) and Cor(***x***|***s***) corresponds to a correlation matrix for an AR(1) process with correlation *ρ*.
- Get the marginal 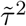 as 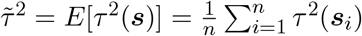 where ***s***_*i*_ is the vector of ancestry proportions for the *i*-th subject in the sample. Note that the 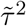 remains constant across each of the simulation replications since the ancestry proportions are generated only once before beginning the simulation study.
- Get the marginal 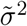 by using the formula 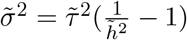.
- Model *σ*^2^(***s***) as:

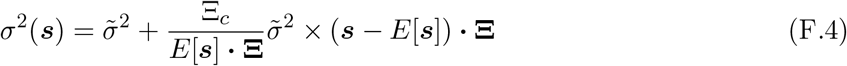

where **Ξ** = (*K* − 1, *K* − 2, …, 0) and Ξ_*c*_ is a constant that takes a value in [0, 1) and can be varied to control the strength of ancestry on *σ*^2^(***s***). This model shares similar characteristics to the model for Var(*x*_*j*_|***s***) from Equation (F.2). Note that the elements in **Ξ** are reversed from **Ω**. We opt for this since it generates a larger range of conditional heritabilities in our simulations as opposed to setting **Ξ** equal to **Ω**.
- Set 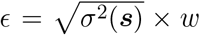 where *w* is drawn from a standard normal distribution. This results in an *ϵ* such that *E*(*ϵ*) = 0 and Var(*ϵ*|***s***) = *σ*^2^(***s***).
- Generate *y* as using the model from Equation (1) by setting

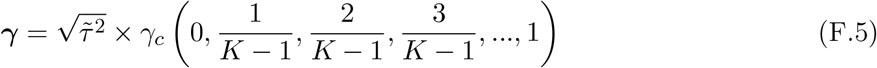

where *γ*_*c*_ is a constant that can be varied to control the strength of ancestry on *y*. By using this choice of ***γ*** we make sure that the weakest and strongest effect of ancestry are 0 and 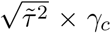 regardless of the choice of the number of parental ancestries *K*. Moreover, it is easy to show that the resulting Var(***γs***) will be proportional to 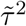, and thus the proportion of variance of *y* attributable to ***s***, 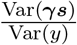, does not change if the value of 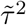changes, and only changes in our simulation when *γ*_*c*_ is varied.
- This process results in a phenotype *y* with the desired marginal heritability 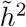that satisfies Equation (2) and a conditional heritability 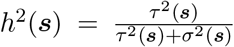 where *τ* ^2^(***s***) and *σ*^2^(***s***) are obtained from Equations (F.3) and (F.4) respectively.

### F.2 Comparison between GWASH and LD-score regression estimates of marginal heritability

To compare the use of LD-score regression with intercept 1 (LDSC-1) and GWASH for estimating the marginal heritability 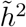, we conduct a simulation study with 100 replications as follows:

- To simulate the SNP effects ***β*** = (*β*_1_, …, *β*_*m*_), we randomly select 10 percent of them to be drawn from standard normal distributions and set the rest of them to be 0. This is done only once at the beginning of the simulation study and kept fixed for the 100 replications.
- We take the number of ancestries to be *K* = 4, since this corresponds to the number of observed ancestries available for our real data analysis in Section H. The ancestry proportions are generated only once at the beginning of the simulation study and kept fixed for the 100 replications.
- The marginal heritability is taken to be 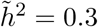 which is close to the estimates obtained in the real data analysis described in Section H.
- We take *ρ* = 0.5 to allow some correlation between SNPs. Although the value we have chosen could be considered mild, later simulations presented in Table F.3 and Figure F.8 show that this parameter does not have much effect on our heritability estimates.
- The constants *λ*_*c*_, Ω_*c*_, Ξ_*c*_ and *γ*_*c*_ are taken to be 1, 0.25, 0.25 and 5 respectively because the combination of these values results in a strong enough ancestry effect on SNPs and phenotypes to have a detrimental effect on naive estimation with GWASH and LDSC-1. One can see this in the later simulations presented in Table F.4 where the naive procedures work fine with a small value of *λ*_*c*_ when the other parameter values are kept the same.

A SNP matrix **X** with *n* = 1000 subjects and *m* = 2000 SNPs, and a corresponding phenotype vector **y** are generated during each replication. The ancestry proportions are assumed to be observed, and so the covariate matrix **S** is of dimensions *n* × *K* where each row corresponds to the ancestry proportions of each subject in the sample. Moreover since 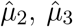 and LD-scores remain stable for large *m* and *n*, they are calculated only calculated once and then kept fixed across the rest of the replications.

For each replication we calculate 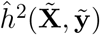 as described in Section 2.5 to estimate the pre-defined covariate-adjusted marginal heritability 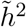by setting 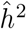 to GWASH and LD-score regression with intercept 1 (LDSC-1). We note that 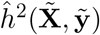 is equivalent to cov-LDSC described in [18] when 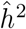 is taken to be the LD-score regression estimator. We use LDSC-1 instead of the unconstrained intercept version because the authors of [2] show that LDSC-1 is a better heritability estimator than the unconstrained version. We use the software provided in [3] to fit LDSC-1 estimates.

Recall that both GWASH and LD-score regression can be calculated from *t*-statistics and LD-scores as detailed in Sections A.3 and A.4. We say that the *t*-statistics are adjusted if they are calculated from 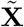 and 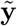, and unadjusted if calculated from **X** and **y**. Similarly, we say that the LD-scores are adjusted if they are calculated from an LD-matrix obtained from 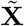 and unadjusted if they are calculated from **X**.

We compare estimates from our proposed framework to estimates obtained from three naive frameworks:

- Naive 1: Calculating GWASH and LDSC-1 estimates with unadjusted LD-scores and unadjusted *t*-statistics.
- Naive 2: Calculating GWASH and LDSC-1 estimates with unadjusted LD-scores and adjusted *t*-statistics.
- Naive 3: Calculating GWASH and LDSC-1 to adjusted LD-scores and unadjusted *t*-statistics.

The first two naive frameworks are also discussed by [18]. The first naive framework represents a case where population structure is not accounted for at all. The second naive framework represents a case where one might use summary statistics obtained from a GWAS that accounts for population structure, but not do any adjustments for LD-scores. Since adjusting for principal components is standard for modern GWAS studies [21], but covariate-adjusted LD scores have not been available before being discussed by [18], one can see such issues arising in practice and it can be of interest to evaluate the effect of this discrepency. Finally, the third naive framework represents a case where a GWAS does not account for population structure, but the LD-scores do. Although this case is not as expected to be observed in practice, we add it in our analysis as an additional comparison that can help understand the performance of our proposed framework and the other two naive frameworks. This becomes more apparent in later sections.

**Table F.1:** Summary of marginal heritability estimation results for the simulation study from Section F.2. We report the mean and standard deviation in parenthesis across the 100 repetitions. The target marginal heritability is 0.3. Any mean that is within one standard deviation of the target heritability is colored in blue, otherwise it is colored in red.

| Estimator | Adjusted | Naive 1 | Naive 2 | Naive 3 | $\hat{\mu}_2$ adjusted | $\hat{\mu}_2$ unadjusted |
| --- | --- | --- | --- | --- | --- | --- |
| GWASH | 0.283<br>(0.057) | 0.951<br>(0.161) | 0.105<br>(0.021) | 2.571<br>(0.437) | 1.681 | 4.546 |
| LDSC-1 | 0.284<br>(0.058) | 0.877<br>(0.139) | 0.113<br>(0.025) | 2.616<br>(0.446) |  |  |

The results from this simulation are summarized in Table F.1. Covariate-adjusted GWASH and LDSC-1 result in very similar estimates for marginal heritability with a mean estimate of 0.283 and 0.284. Both mean estimates are within one standard deviation of the true simulated marginal heritability of 0.3. Conversely, the first and second naive frameworks overestimate and underestimate the marginal heritability respectively. This coincides with a trend described in [18]. Finally, the third naive framework overestimates the marginal heritability even more than the first since it makes use of LD-scores of a lower magnitude due to their adjustment whilst still using inflated *t*-statistics. In the case of the GWASH estimator this leads to a larger numerator and a smaller denominator in Equation (A.3), and we can expect a similar behaviour with LDSC-1 since the authors of [2] show that LDSC-1 and GWASH share similar asymptotic properties. Moreover, we do not analyze the free-intercept version of LD-Score regression since [2] also shows that this estimator does not remedy bias due to population stratification.

### F.3 Estimation of conditional heritability and standard errors

We now explore the performance of our proposed framework to estimate conditional heritabilities and their standard error. Due to the similarity of LDSC-1 and GWASH results in the previous simulation and their asymptotic equivalence discussed in [33], we restrict ourselves to using GWASH for the marginal heritability estimation needed to estimate conditional heritabilities in this section.

We use the same settings as Section F.2, and since ancestry proportions are continuous and kept fixed at the beginning of the simulation, each of the *n* = 1000 subjects will have a corresponding conditional heritability that will not vary across replications. In each of the 100 replications we calculate conditional heritability estimates for each subject as described in Section 2.5. We use these to get an average estimated conditional heritability for each of the *n* = 1000 subjects and compare these to the true values. Empirical standard deviations for the estimates are calculated too. Our results are summarized in Figure F.5 which shows that all true conditional heritabilities are within one standard deviation of their average estimates.

**Figure F.5:**
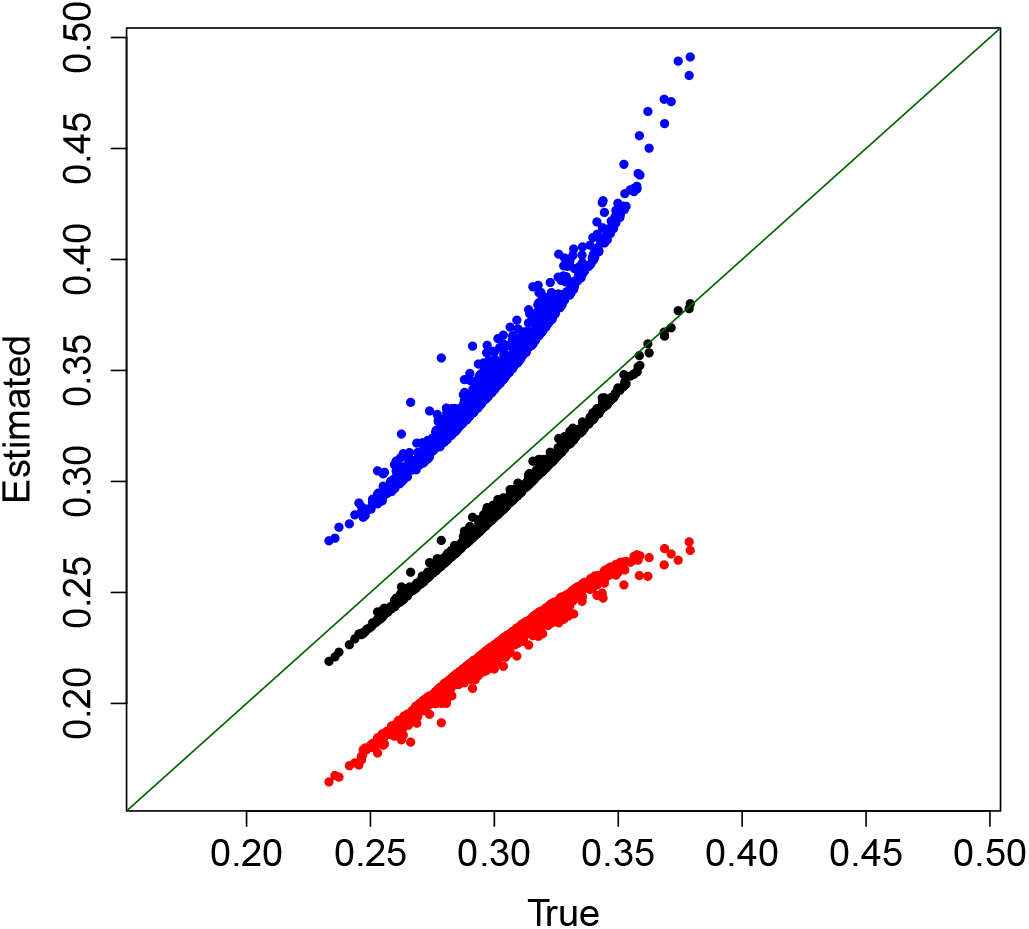
Comparisons of true conditional heritabilities vs their average estimates (black) +/ −1 empirical S.D. (blue/red) across 100 simulations for each of the *n* = 1000 for the simulation study from Section F.3. The true conditional heritabilities are all within 1 empirical S.D. of the average estimates.

We also calculate estimates for the conditional heritability standard errors by taking the square-root of Equation (E.1) in each replication. We then calculate an average of this for each subject and compare to the Monte Carlo standard errors calculated from the 100 replications. We can see that the average standard error estimates using Equation (E.1) correlate with the Monte Carlo standard errors as shown in Figure F.6a. There seems to be a bias upwards, but it is preferable for practical purposes to over-estimate standard errors than underestimate them. That said, as illustrated in Figure F.6b, the over-estimation is not too high, with the highest bias being under 0.008 which is small considering that the conditional heritability ranges from 0.23 to 0.38. Moreover, there are only three average estimated standard errors that are lower than the Monte Carlo standard errors, and these only seem to be slightly underestimated.

**Figure F.6:**
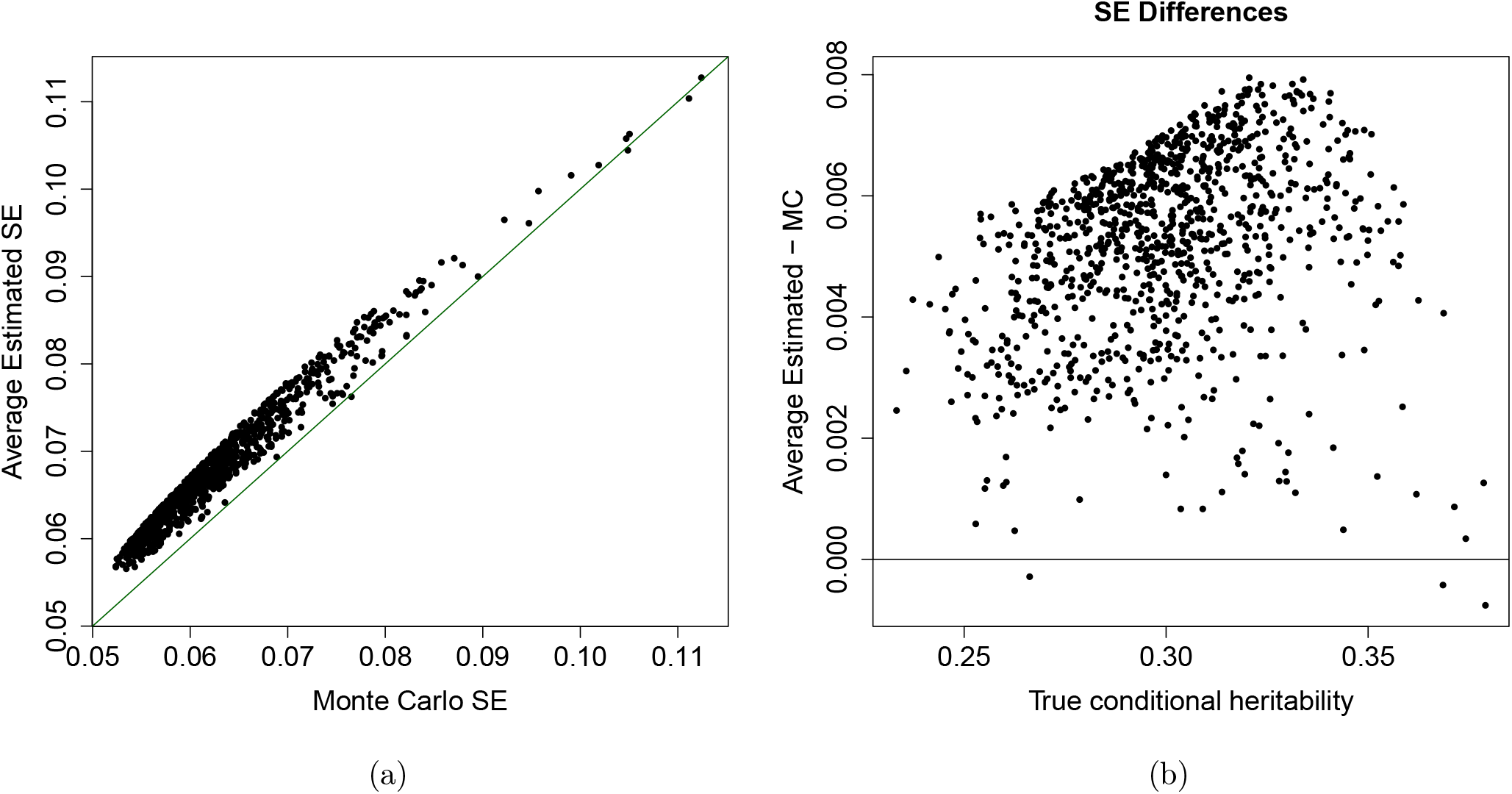
Comparison of Monte Carlo standard errors and average estimated standard errors using Equation (E.1) for the simulation study from Section F.3. The two are correlated with each other, with the average estimated standard errors tending to be only slightly higher.

### F.4 Effect of varying simulation parameters

We now conduct more simulations to explore the sensitivity of our methods to varying simulation parameters. The target marginal heritability 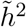 is varied from 0.1 to 0.9 in increments of 0.2. We use this range for the conditional SNP correlation *ρ* too. The strength of ancestry effects on SNP means and the phenotype, *λ*_*c*_ and *γ*_*c*_ respectively, are evaluated at 0.01, 1, 5, 10 and 100 to examine our methods at a very low ancestry effect, a very high ancestry effect and three intermediate cases. Similarly, we evaluate the impact of changing Ω_*c*_ and Ξ_*c*_, the strength of ancestry effects on SNP variance and *σ*^2^(***s***) respectively, by examining the behaviour near the boundaries of their parameter spaces at 0.01 and 0.99, and three intermediate values: 0.25, 0.5, and 0.75. Finally, the number of ancestries *K* is varied from 2 to 10 in increments of 2.

Tables F.2 - F.8 summarize the estimated marginal heritabilities from these simulations. We conduct each simulation analogous to Section F.2 and only change the value of the parameter of interest whilst the others are kept to their values from before. Since GWASH and LDSC-1 gave similar results in a previous analysis, we restrict ourselves to just using GWASH. The covariate-adjusted marginal heritabilities are shown to be within one standard deviation of the target in all cases.

Although Naive 1 and Naive 3 generally overestimate the target heritability, and Naive 2 generally underestimates it, there are some cases where this does not hold true. There are certain cases where some of the naive frameworks also lead to estimates that are within one standard deviation of the target. This is the case for Naive 1-3 with *λ*_*c*_ = 0.01, Naive 3 with *γ*_*c*_ values of 0.01 and 1, and Naive 2 with *K* = 10. The behaviour observed with *γ*_*c*_ can partially be explained by unadjusted *t*-statistics being similar to adjusted *t*-statistics when the impact of ancestry on the phenotype is low. That said, we can see from Table F.5 that the standard deviation of Naive 3 is also larger than that of the covariate-adjusted estimator for the corresponding values of *γ*_*c*_. To understand the behaviour with *λ*_*c*_ we can see from Table F.4 that at *λ*_*c*_ = 0.01, the adjusted and unadjusted 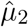 are very similar due to there being few differences in allele frequencies between populations. Regarding the behaviour at *K* = 10, we can see from Table F.8 that the unadjusted 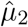 decreases as *K* increases, and with *K* = 10 it is close enough to its adjusted form for Naive 2 to be close to the target heritability. It is possible that there is a limit to how different the genotypes from each population can be from each other in order to keep all other parameters and settings fixed under our simulations, and thus with a large *K* the population genotypes start becoming more similar to each other which is reflected in the increasing convergence between the adjusted and unadjusted 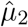. It is also interesting to see that even though they do not get within one standard deviation of the target heritability, Naive 1 and Naive 2 also seem to get closer to the desired value as *K* increases. Finally, there are also some cases where Naive 1 can underestimate heritability. For example, from Tables F.4 and F.5 we can see that Naive 1 underestimates the target heritability when *λ*_*c*_ is large and also when *γ*_*c*_ is low. This suggests that unaccounted high population heterogeneity can lead to underestimated heritability when it affects allele frequencies more than the phenotype.

We also calculate estimates for conditional heritabilities, and summarize our results in Figures F.7 - F.13. We conduct each simulation analogous to Section F.3 and only change the value of the simulation parameter of interest. The true conditional heritabilities are within one standard deviation of their average estimates for all examined cases.

**Table F.2:** Summary of the marginal heritability estimation simulation study for varying levels of 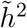 (target marginal heritability). The other parameter values are specified in Section F.2. We report the mean and standard deviation in parenthesis across the 100 repetitions. Any mean that is within one standard deviation of the target heritability is colored in blue, otherwise it is colored in red.

| $\tilde{h}^2$ | Adjusted | Naive 1 | Naive 2 | Naive 3 | $\hat{\mu}_2$ adjusted | $\hat{\mu}_2$ unadjusted |
| --- | --- | --- | --- | --- | --- | --- |
| 0.1 | 0.089<br>(0.048) | 0.342<br>(0.113) | 0.033<br>(0.018) | 0.924<br>(0.307) | 1.681 | 4.546 |
| 0.3 | 0.283<br>(0.057) | 0.951<br>(0.161) | 0.105<br>(0.021) | 2.571<br>(0.437) | 1.681 | 4.546 |
| 0.5 | 0.508<br>(0.065) | 1.431<br>(0.160) | 0.190<br>(0.024) | 3.820<br>(0.428) | 1.684 | 4.495 |
| 0.7 | 0.725<br>(0.070) | 2.015<br>(0.198) | 0.256<br>(0.025) | 5.698<br>(0.560) | 1.675 | 4.737 |
| 0.9 | 0.878<br>(0.067) | 2.642<br>(0.197) | 0.303<br>(0.023) | 7.643<br>(0.570) | 1.677 | 4.850 |

**Table F.3:** Summary of the marginal heritability estimation simulation study for varying levels of *ρ* (correlation between contiguous SNPs conditional on ancestry). The other parameter values are specified in Section F.2. We report the mean and standard deviation in parenthesis across the 100 repetitions. The target marginal heritability is 0.3. Any mean that is within one standard deviation of the target heritability is colored in blue, otherwise it is colored in red.

| $\rho$ | Adjusted | Naive 1 | Naive 2 | Naive 3 | $\hat{\mu}_2$ adjusted | $\hat{\mu}_2$ unadjusted |
| --- | --- | --- | --- | --- | --- | --- |
| 0.1 | 0.287<br>(0.062) | 1.060<br>(0.186) | 0.074<br>(0.016) | 4.092<br>(0.717) | 1.036 | 4.002 |
| 0.3 | 0.302<br>(0.063) | 0.971<br>(0.156) | 0.089<br>(0.019) | 3.283<br>(0.527) | 1.211 | 4.094 |
| 0.5 | 0.283<br>(0.057) | 0.951<br>(0.161) | 0.105<br>(0.021) | 2.571<br>(0.437) | 1.681 | 4.546 |
| 0.7 | 0.298<br>(0.050) | 0.919<br>(0.129) | 0.147<br>(0.025) | 1.860<br>(0.260) | 2.918 | 5.905 |
| 0.9 | 0.291<br>(0.045) | 0.561<br>(0.075) | 0.243<br>(0.038) | 0.672<br>(0.090) | 9.522 | 11.406 |

**Table F.4:** Summary of the marginal heritability estimation simulation study for varying levels of *λ*_*c*_ (strength of ancestry effects on SNP means). The other parameter values are specified in Section F.2. We report the mean and standard deviation in parenthesis across the 100 repetitions. The target marginal heritability is 0.3. Any mean that is within one standard deviation of the target heritability is colored in blue, otherwise it is colored in red.

| $\lambda_c$ | Adjusted | Naive 1 | Naive 2 | Naive 3 | $\hat{\mu}_2$ adjusted | $\hat{\mu}_2$ unadjusted |
| --- | --- | --- | --- | --- | --- | --- |
| 0.01 | 0.283<br>(0.057) | 0.257<br>(0.058) | 0.284<br>(0.058) | 0.256<br>(0.058) | 1.681 | 1.674 |
| 1 | 0.283<br>(0.057) | 0.951<br>(0.161) | 0.105<br>(0.021) | 2.571<br>(0.437) | 1.681 | 4.546 |
| 5 | 0.306<br>(0.061) | 0.151<br>(0.025) | 0.002<br>(0.000) | 18.873<br>(3.139) | 1.684 | 210.292 |
| 10 | 0.308<br>(0.055) | 0.115<br>(0.021) | 0.001<br>(0.000) | 29.802<br>(5.338) | 1.675 | 432.662 |
| 100 | 0.295<br>(0.057) | 0.103<br>(0.016) | 0.001<br>(0.000) | 40.571<br>(6.445) | 1.677 | 661.587 |

**Table F.5:** Summary of the marginal heritability estimation simulation study for varying levels of *γ*_*c*_ (strength of ancestry effects on *y*). The other parameter values are specified in Section F.2. We report the mean and standard deviation in parenthesis across the 100 repetitions. The target marginal heritability is 0.3. Any mean that is within one standard deviation of the target heritability is colored in blue, otherwise it is colored in red.

| $\gamma_c$ | Adjusted | Naive 1 | Naive 2 | Naive 3 | $\hat{\mu}_2$ adjusted | $\hat{\mu}_2$ unadjusted |
| --- | --- | --- | --- | --- | --- | --- |
| 0.01 | 0.283<br>(0.057) | 0.094<br>(0.031) | 0.105<br>(0.021) | 0.255<br>(0.083) | 1.681 | 4.546 |
| 1 | 0.306<br>(0.061) | 0.143<br>(0.042) | 0.115<br>(0.023) | 0.383<br>(0.113) | 1.684 | 4.495 |
| 5 | 0.283<br>(0.057) | 0.951<br>(0.161) | 0.105<br>(0.021) | 2.571<br>(0.437) | 1.681 | 4.546 |
| 10 | 0.308<br>(0.055) | 2.812<br>(0.201) | 0.109<br>(0.019) | 7.951<br>(0.569) | 1.675 | 4.737 |
| 100 | 0.295<br>(0.057) | 9.541<br>(0.086) | 0.102<br>(0.020) | 27.598<br>(0.249) | 1.677 | 4.850 |

**Table F.6:** Summary of the marginal heritability estimation simulation study for varying levels of Ω_*c*_ (strength of ancestry effects on SNP variance). The other parameter values are specified in Section F.2. We report the mean and standard deviation in parenthesis across the 100 repetitions. The target marginal heritability is 0.3. Any mean that is within one standard deviation of the target heritability is colored in blue, otherwise it is colored in red.

| $\Omega_c$ | Adjusted | Naive 1 | Naive 2 | Naive 3 | $\hat{\mu}_2$ adjusted | $\hat{\mu}_2$ unadjusted |
| --- | --- | --- | --- | --- | --- | --- |
| 0.01 | 0.286<br>(0.058) | 0.953<br>(0.162) | 0.106<br>(0.021) | 2.576<br>(0.438) | 1.674 | 4.522 |
| 0.25 | 0.283<br>(0.057) | 0.951<br>(0.161) | 0.105<br>(0.021) | 2.571<br>(0.437) | 1.681 | 4.546 |
| 0.5 | 0.302<br>(0.061) | 0.912<br>(0.139) | 0.115<br>(0.023) | 2.404<br>(0.367) | 1.702 | 4.489 |
| 0.75 | 0.300<br>(0.054) | 0.964<br>(0.156) | 0.109<br>(0.020) | 2.649<br>(0.428) | 1.731 | 4.758 |
| 0.99 | 0.287<br>(0.056) | 1.042<br>(0.157) | 0.104<br>(0.020) | 2.864<br>(0.432) | 1.783 | 4.902 |

**Table F.7:** Summary of the marginal heritability estimation simulation study for varying levels of Ξ_*c*_ (strength of ancestry effects on *σ*^2^(***s***)). The other parameter values are specified in Section F.2. We report the mean and standard deviation in parenthesis across the 100 repetitions. The target marginal heritability is 0.3. Any mean that is within one standard deviation of the target heritability is colored in blue, otherwise it is colored in red.

| $\Xi_c$ | Adjusted | Naive 1 | Naive 2 | Naive 3 | $\hat{\mu}_2$ adjusted | $\hat{\mu}_2$ unadjusted |
| --- | --- | --- | --- | --- | --- | --- |
| 0.01 | 0.286<br>(0.057) | 0.953<br>(0.162) | 0.106<br>(0.021) | 2.579<br>(0.438) | 1.681 | 4.546 |
| 0.25 | 0.283<br>(0.057) | 0.951<br>(0.161) | 0.105<br>(0.021) | 2.571<br>(0.437) | 1.681 | 4.546 |
| 0.5 | 0.305<br>(0.061) | 0.916<br>(0.141) | 0.114<br>(0.023) | 2.444<br>(0.376) | 1.684 | 4.495 |
| 0.75 | 0.303<br>(0.055) | 0.972<br>(0.156) | 0.107<br>(0.019) | 2.750<br>(0.441) | 1.675 | 4.737 |
| 0.99 | 0.287<br>(0.057) | 1.056<br>(0.164) | 0.099<br>(0.020) | 3.054<br>(0.476) | 1.677 | 4.850 |

**Table F.8:** Summary of the marginal heritability estimation simulation study for varying levels of *K* (number of underlying ancestries). The other parameter values are specified in Section F.2. We report the mean and standard deviation in parenthesis across the 100 repetitions. The target marginal heritability is 0.3. Any mean that is within one standard deviation of the target heritability is colored in blue, otherwise it is colored in red.

| $K$ | Adjusted | Naive 1 | Naive 2 | Naive 3 | $\hat{\mu}_2$ adjusted | $\hat{\mu}_2$ unadjusted |
| --- | --- | --- | --- | --- | --- | --- |
| 2 | 0.279<br>(0.059) | 3.141<br>(0.217) | 0.030<br>(0.006) | 29.138<br>(2.013) | 1.698 | 15.750 |
| 4 | 0.283<br>(0.057) | 0.951<br>(0.161) | 0.105<br>(0.021) | 2.571<br>(0.437) | 1.681 | 4.546 |
| 6 | 0.320<br>(0.064) | 0.562<br>(0.105) | 0.215<br>(0.043) | 0.838<br>(0.157) | 1.675 | 2.496 |
| 8 | 0.296<br>(0.054) | 0.407<br>(0.087) | 0.240<br>(0.044) | 0.502<br>(0.107) | 1.685 | 2.075 |
| 10 | 0.304<br>(0.066) | 0.389<br>(0.080) | 0.278<br>(0.060) | 0.426<br>(0.087) | 1.686 | 1.847 |

**Figure F.7:**
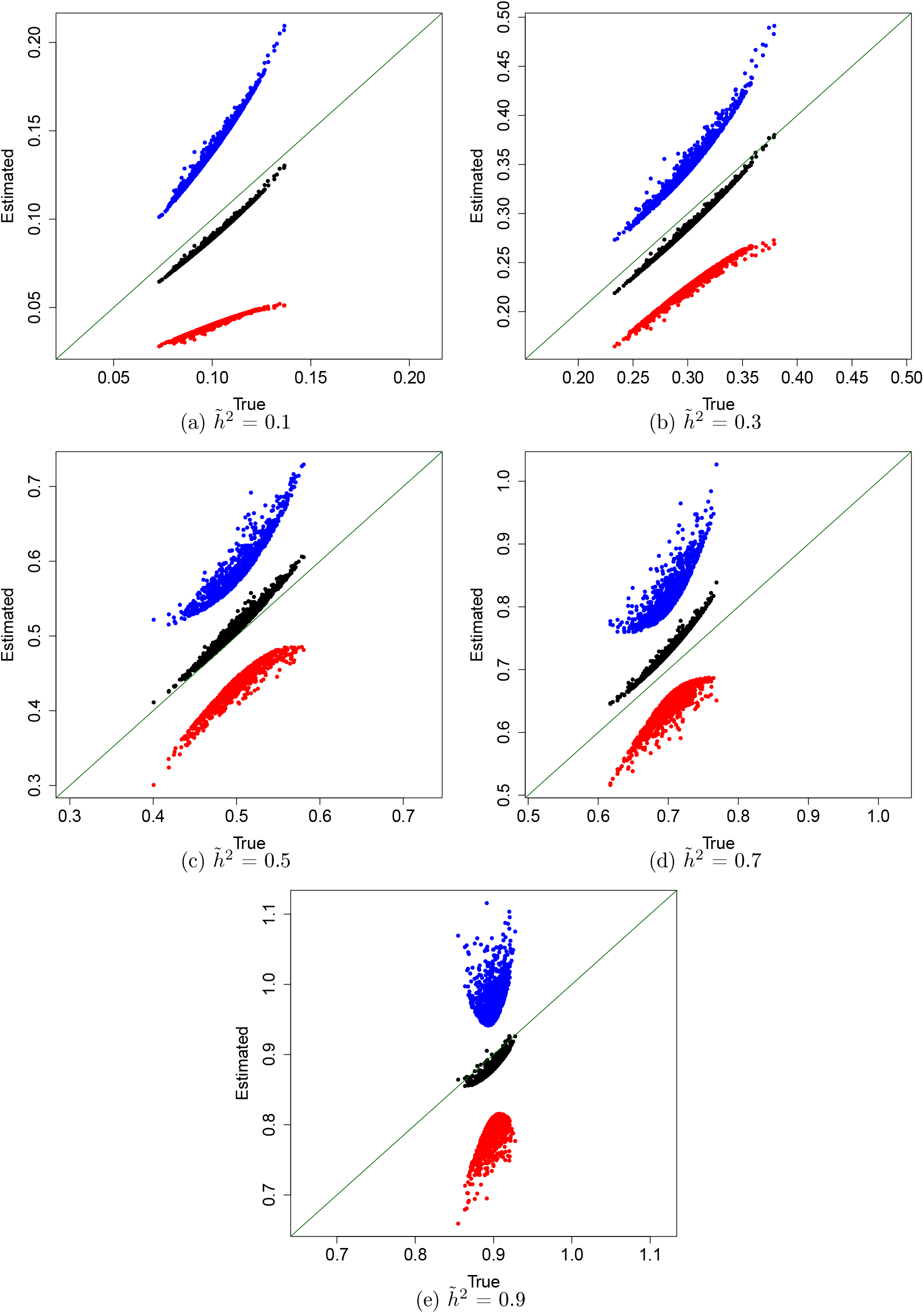
Comparisons of true conditional heritabilities vs their average estimates (black) +/ −1 empirical S.D. (blue/red) across 100 simulations for each of the *n* = 1000 subjects with varying values of 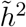. The other parameter values are specified in Section F.2. The true conditional heritabilities are all within 1 empirical S.D. of the average estimates.

**Figure F.8:**
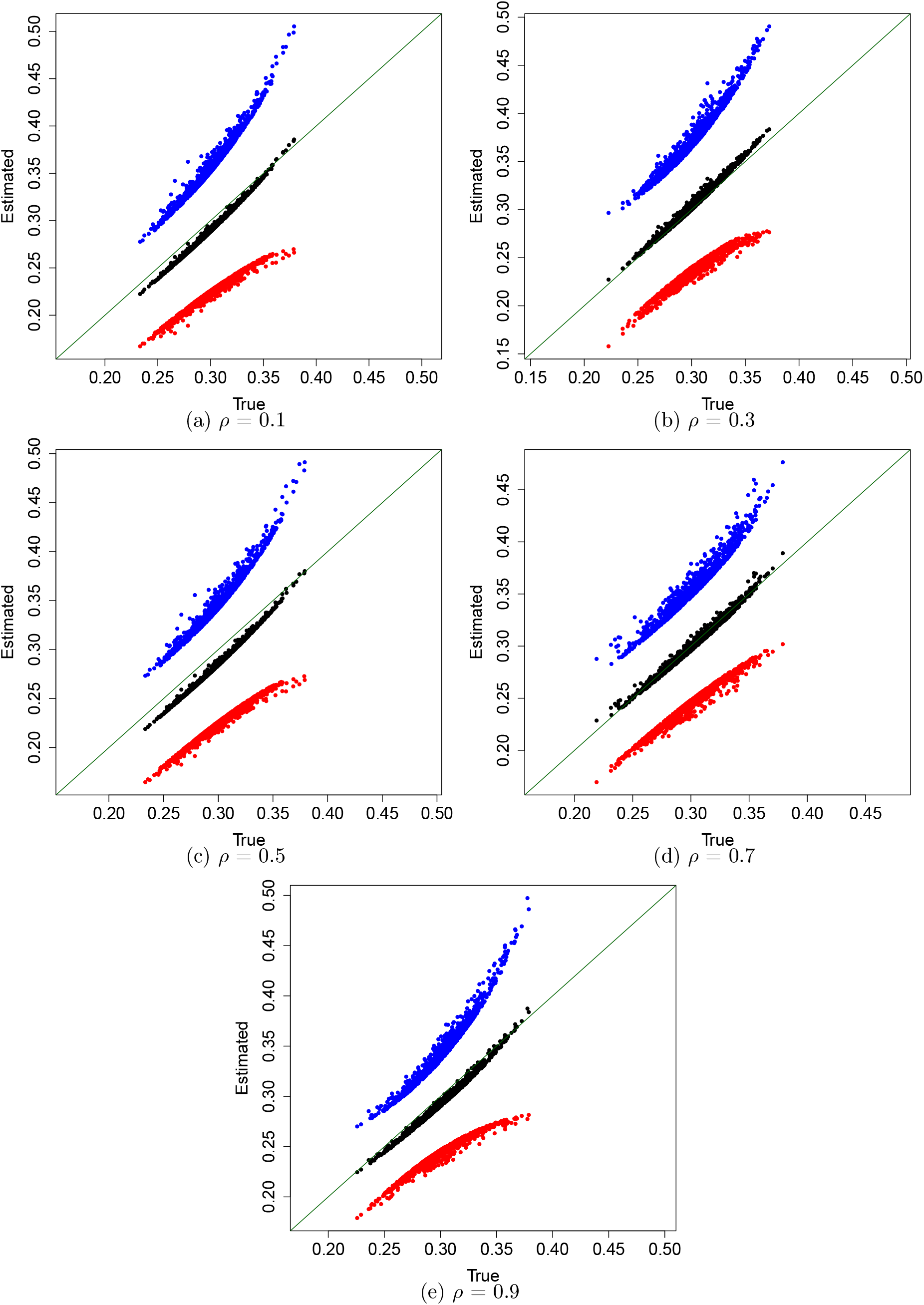
Comparisons of true conditional heritabilities vs their average estimates (black) +/ −1 empirical S.D. (blue/red) across 100 simulations for each of the *n* = 1000 subjects with varying values of *ρ*. The other parameter values are specified in Section F.2. The true conditional heritabilities are all within 1 empirical S.D. of the average estimates.

**Figure F.9:**
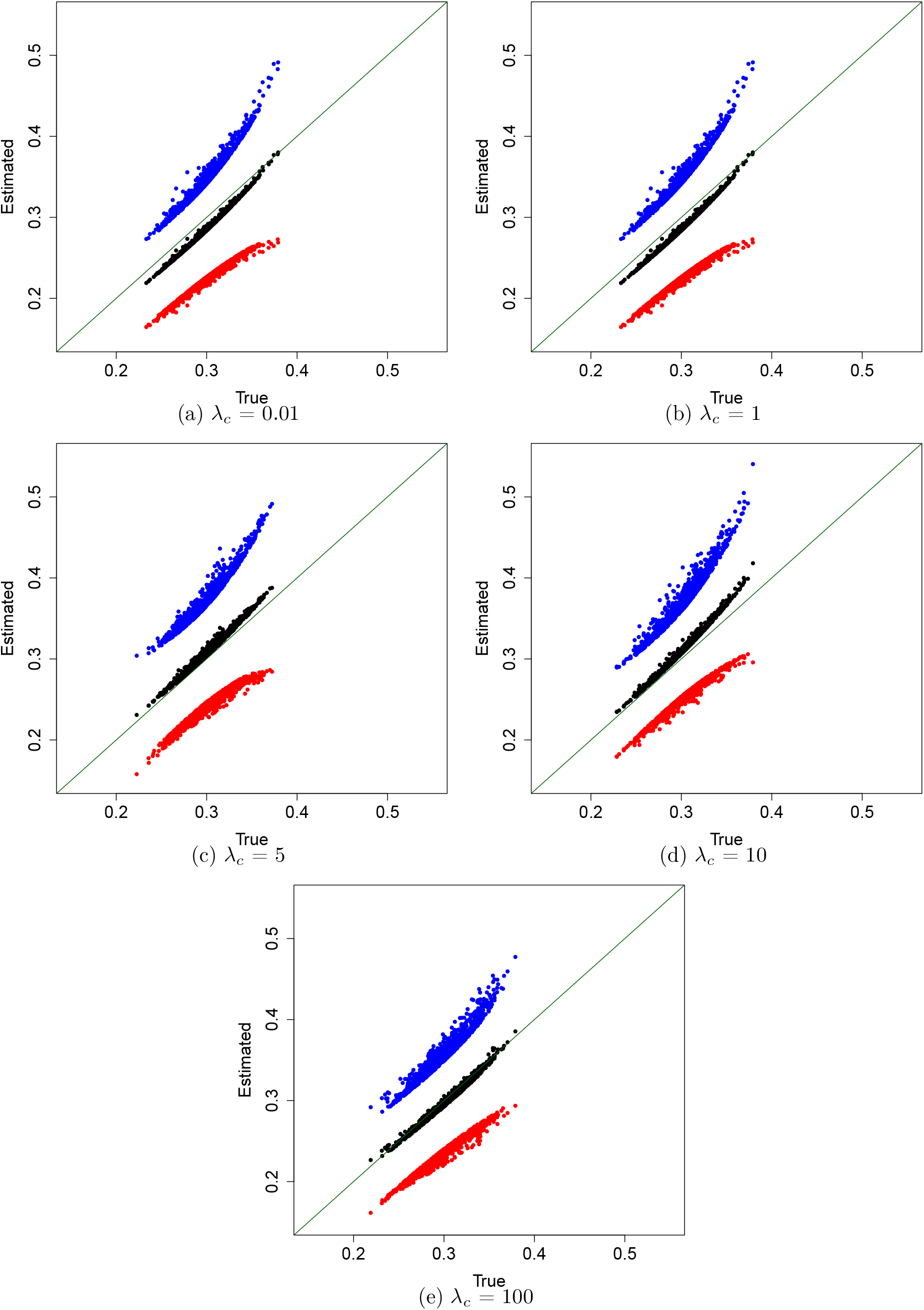
Comparisons of true conditional heritabilities vs their average estimates (black) +/ −1 empirical S.D. (blue/red) across 100 simulations for each of the *n* = 1000 subjects with varying values of *λ*_*c*_. The other parameter values are specified in Section F.2. The true conditional heritabilities are all within 1 empirical S.D. of the average estimates.

**Figure F.10:**
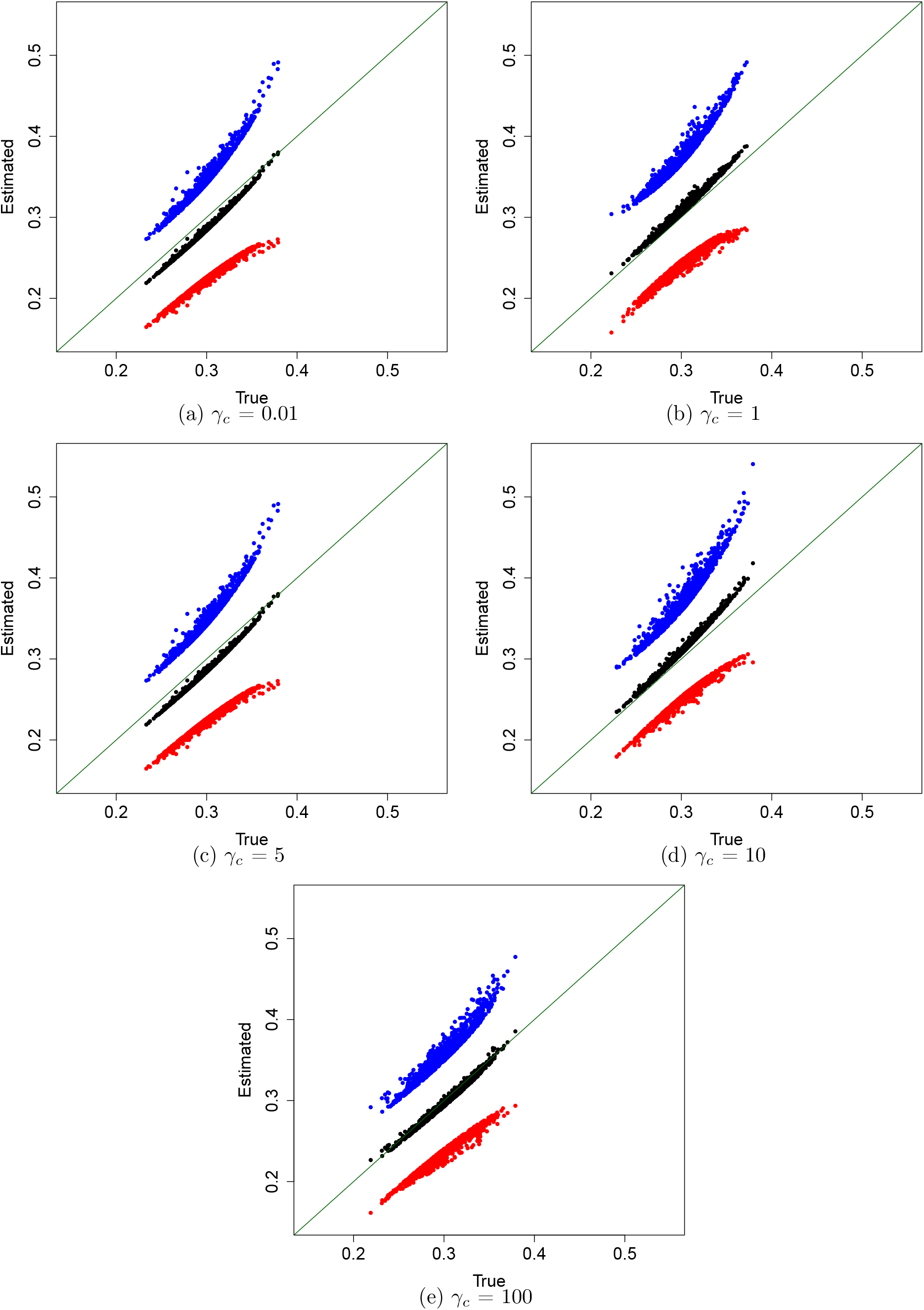
Comparisons of true conditional heritabilities vs their average estimates (black) +/ −1 empirical S.D. (blue/red) across 100 simulations for each of the *n* = 1000 subjects with varying values of γ_*c*_. The other parameter values are specified in Section F.2. The true conditional heritabilities are all within 1 empirical S.D. of the average estimates.

**Figure F.11:**
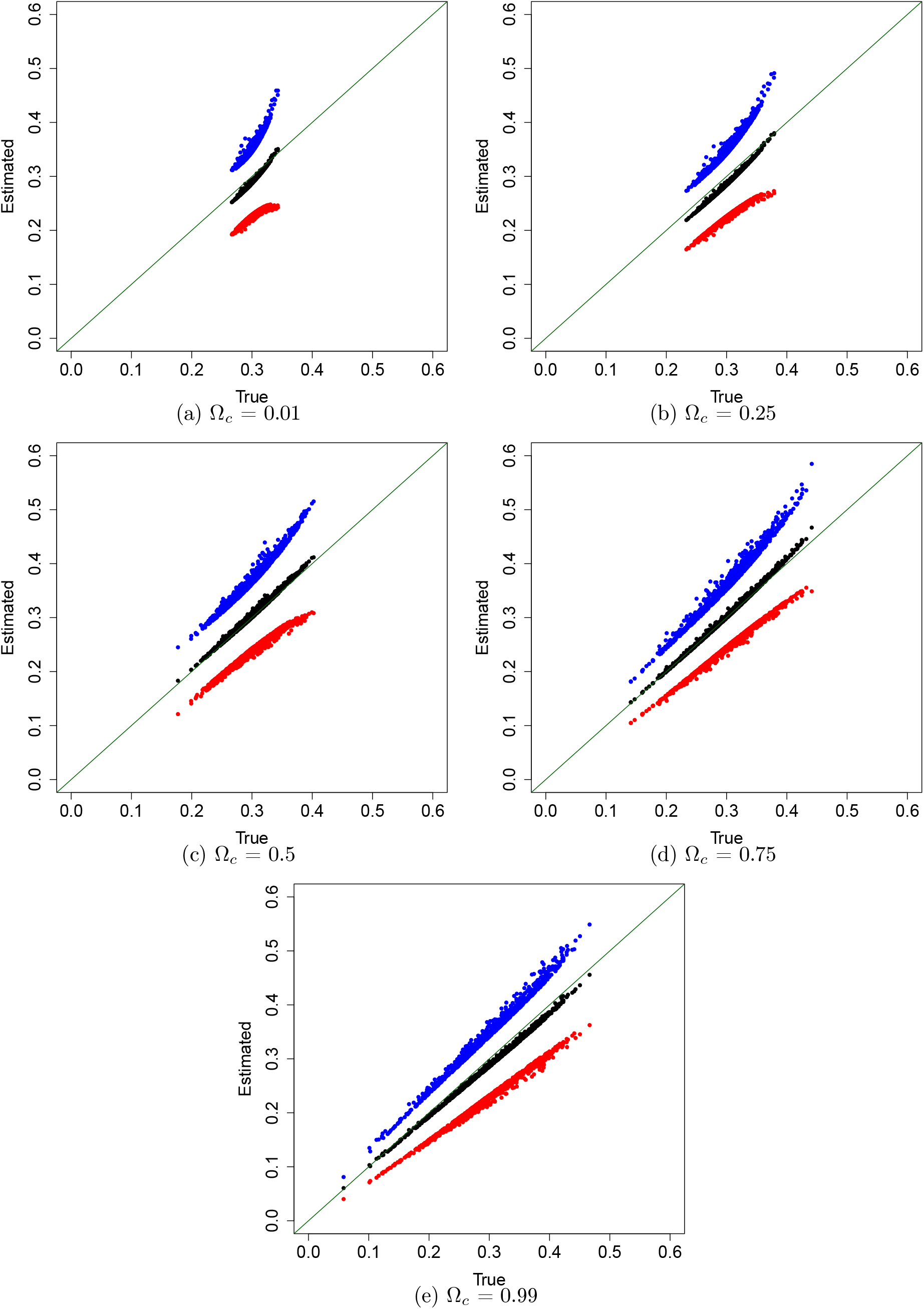
Comparisons of true conditional heritabilities vs their average estimates (black) +/ −1 empirical S.D. (blue/red) across 100 simulations for each of the *n* = 1000 subjects with varying values of Ω_*c*_. The other parameter values are specified in Section F.2. The true conditional heritabilities are all within 1 empirical S.D. of the average estimates.

**Figure F.12:**
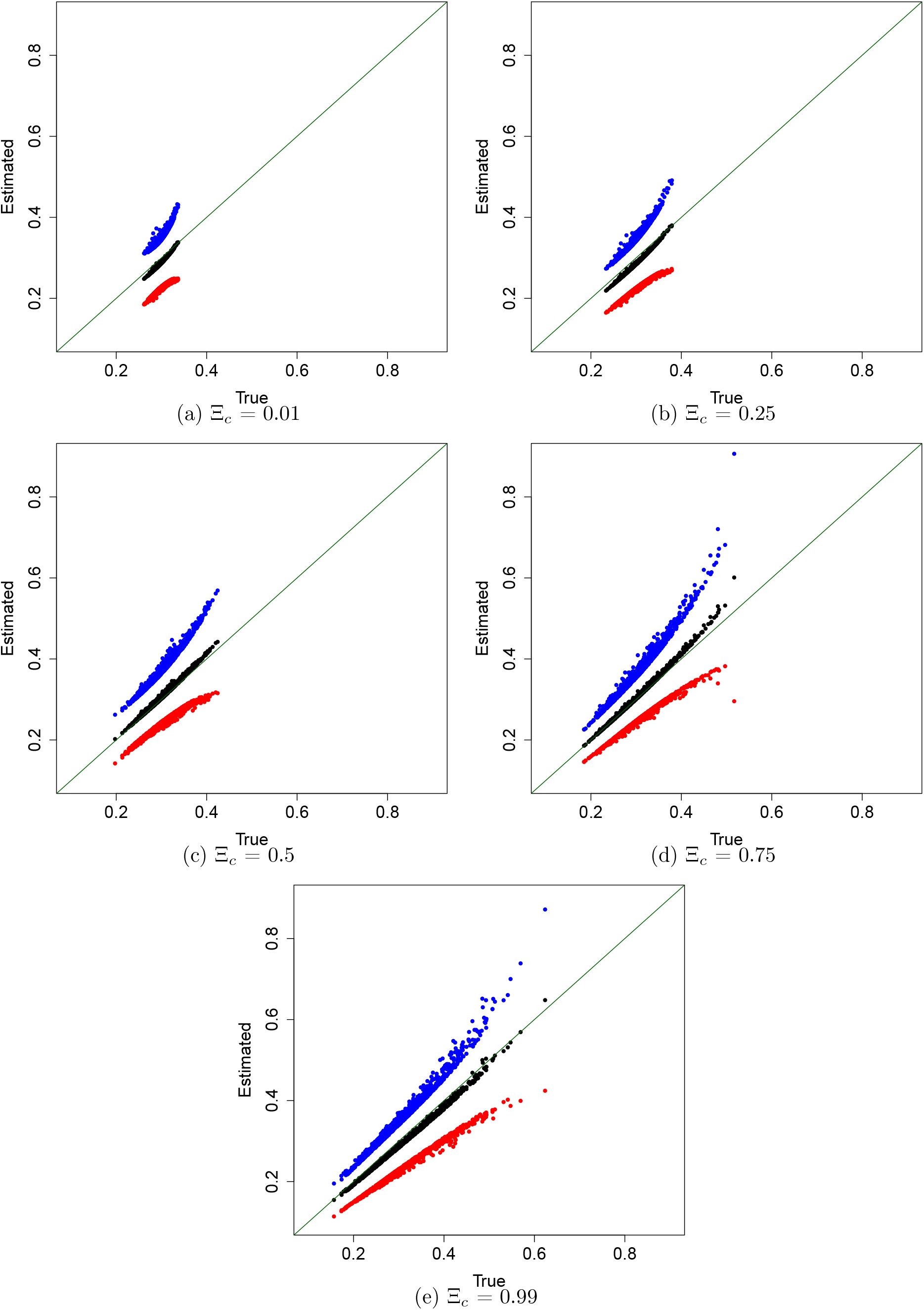
Comparisons of true conditional heritabilities vs their average estimates (black) +/ −1 empirical S.D. (blue/red) across 100 simulations for each of the *n* = 1000 subjects with varying values of Ξ_*c*_. The other parameter values are specified in Section F.2. The true conditional heritabilities are all within 1 empirical S.D. of the average estimates.

**Figure F.13:**
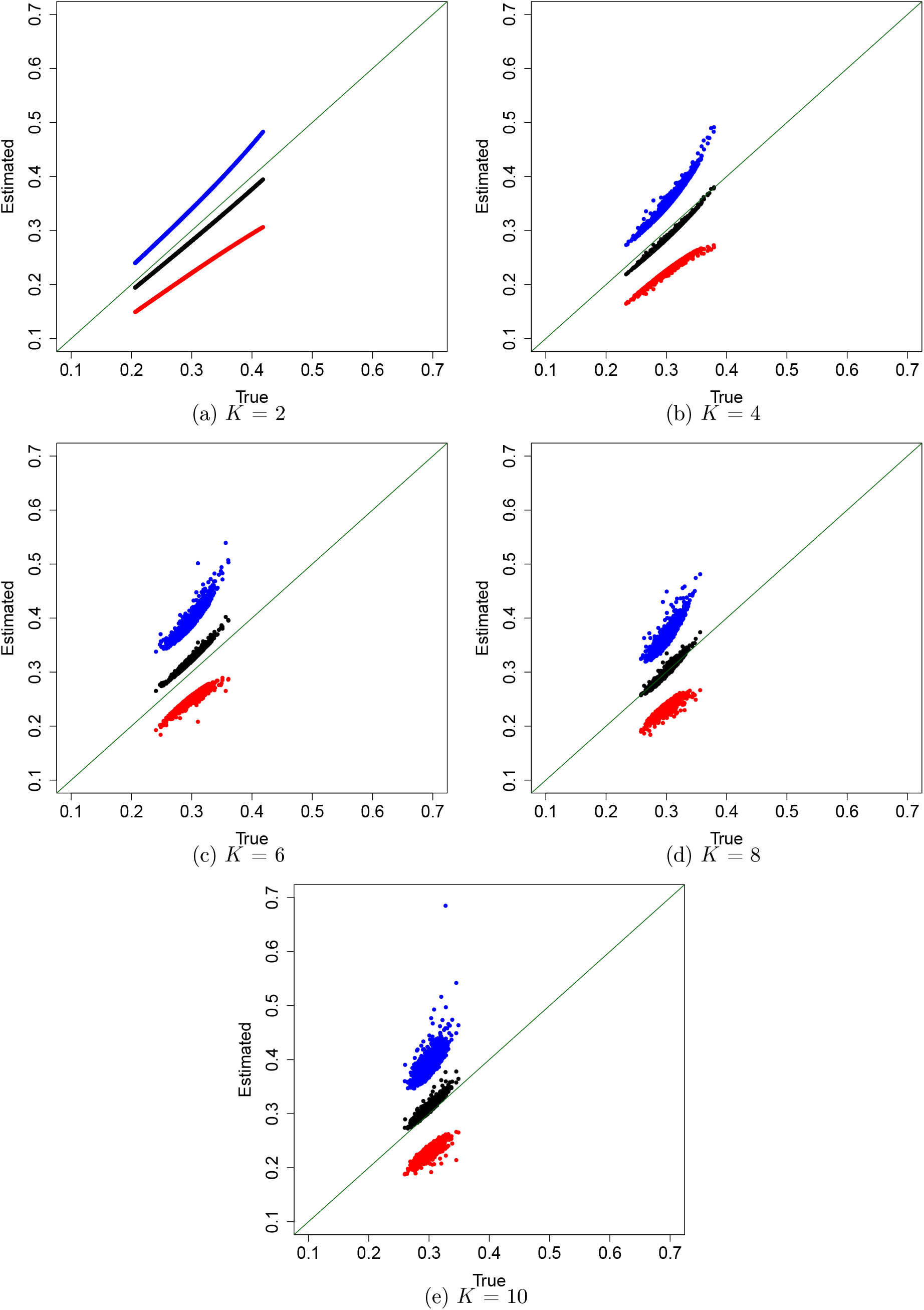
Comparisons of true conditional heritabilities vs their average estimates (black) +/ −1 empirical S.D. (blue/red) across 100 simulations for each of the *n* = 1000 subjects with varying values of p. The other parameter values are specified in Section F.2. The true conditional heritabili are all within 1 empirical S.D. of the average estimates.

### F.5 Sensitivity to use of principal components

We are also interested in seeing the effect of using principal components instead of ancestry proportions for our covariate-adjusted estimation procedures. This mimics the scenario in which we do not observe ancestry proportions and instead need to use principal components as proxies for these. We use the same simulation settings and parameters described in Section F.2, but change the covariate matrix **S** to be composed of 1-10 principal components along with a column on 1s for an intercept term. The principal components are obtained as described in Section A.5 and since we expect them to remain stable for large *m* and *n* they are only calculated once and then kept fixed across the rest of the replications. Note that even though we now use principal components and an intercept term for **S**, we still use simulated ancestry proportions ***s*** for generating SNP matrices and phenotypes as described in Section F.1.

We present the results for covariate-adjusted marginal heritability estimation with varying number of principal components in Table F.9. We can see that that the heritability estimates come close to the target heritability when using 3 or more principal components, and are underestimated otherwise. A similar behaviour is observed with conditional heritability estimates as shown in Figure F.14 where GWASH adjusted with principal components is used for the associated estimate of marginal heritability needed for conditional heritabilities estimates.

The number of principal components needed to adequately estimate covariate-adjusted marginal and conditional heritabilities matches the theory discussed previously. Since we have *K* = 4 ancestries, we need to include *K* − 1 = 3 principal components as discussed in Section 2.6. Moreover, including more than *K* − 1 principal components does not seem to negatively impact the estimates in this simulation study.

**Table F.9:** Summary of the marginal heritability estimation simulation study for estimating marginal heritability using varying number of principal components included as covariates. We report the mean and standard deviation in parenthesis across the 100 repetitions. The target marginal heritability is 0.3. Any mean that is within one standard deviation of the target heritability is colored in blue, otherwise it is colored in red. As a reference, the mean estimates of covariate-adjusted GWASH and LDSC-1 with ancestry proportions are 0.283 (S.D. = 0.057) and 0.284 (S.D. = 0.058) respectively.

| num_PCs | GWASH Adjusted | LDSC-1 Adjusted | $\hat{\mu}_2$ adjusted | $\hat{\mu}_2$ unadjusted |
| --- | --- | --- | --- | --- |
| 1 | 0.153<br>(0.123) | 0.166<br>(0.099) | 3.531 | 4.546 |
| 2 | 0.200<br>(0.041) | 0.212<br>(0.047) | 2.634 | 4.564 |
| 3 | 0.310<br>(0.063) | 0.312<br>(0.063) | 1.675 | 4.735 |
| 4 | 0.308<br>(0.067) | 0.309<br>(0.068) | 1.680 | 5.011 |
| 5 | 0.293<br>(0.060) | 0.294<br>(0.061) | 1.681 | 4.591 |
| 6 | 0.307<br>(0.066) | 0.309<br>(0.067) | 1.673 | 5.240 |
| 7 | 0.304<br>(0.063) | 0.305<br>(0.063) | 1.671 | 4.914 |
| 8 | 0.317<br>(0.060) | 0.317<br>(0.060) | 1.664 | 4.850 |
| 9 | 0.296<br>(0.051) | 0.297<br>(0.052) | 1.668 | 4.731 |
| 10 | 0.313<br>(0.058) | 0.314<br>(0.058) | 1.664 | 4.612 |

**Figure F.14:**
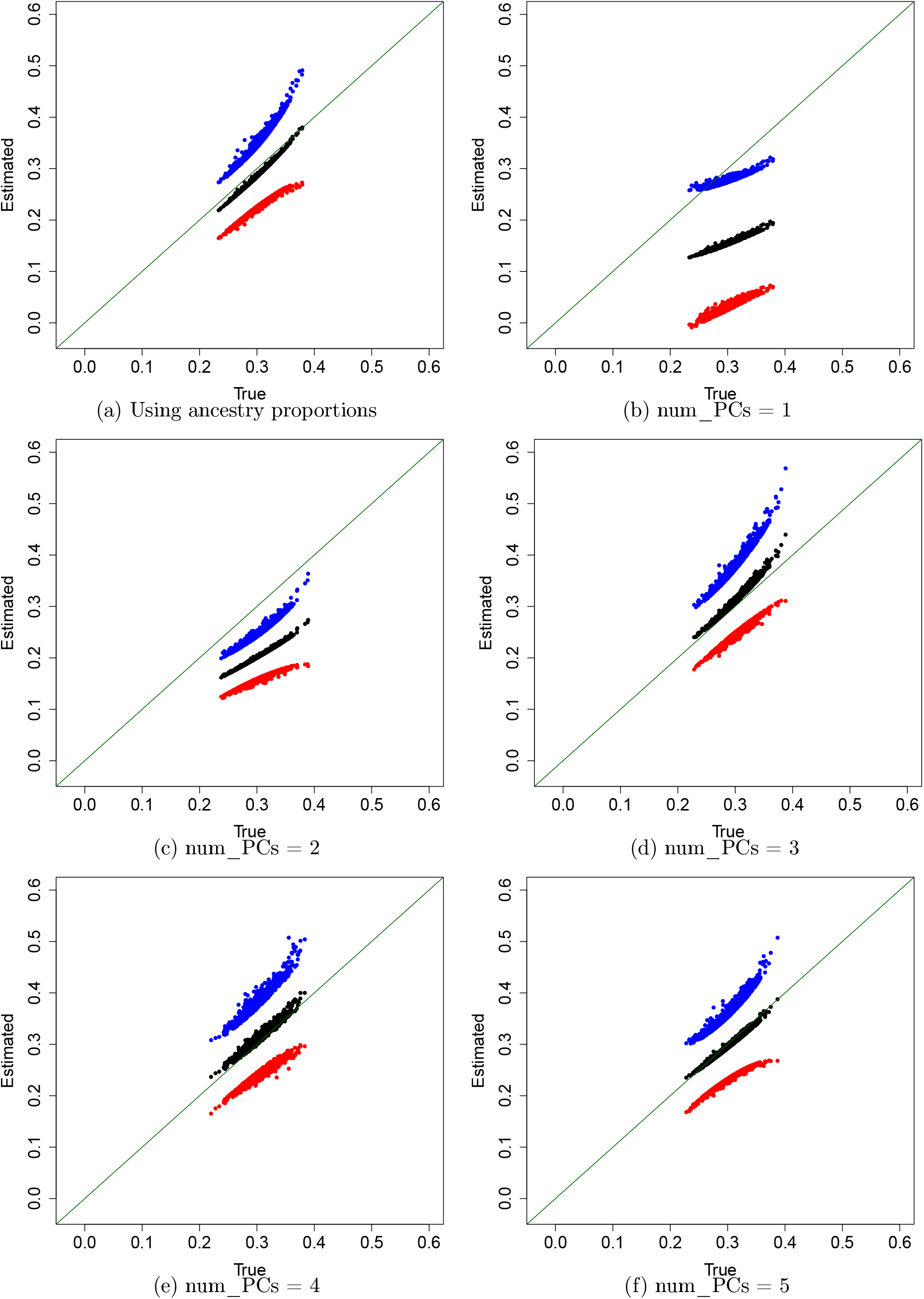
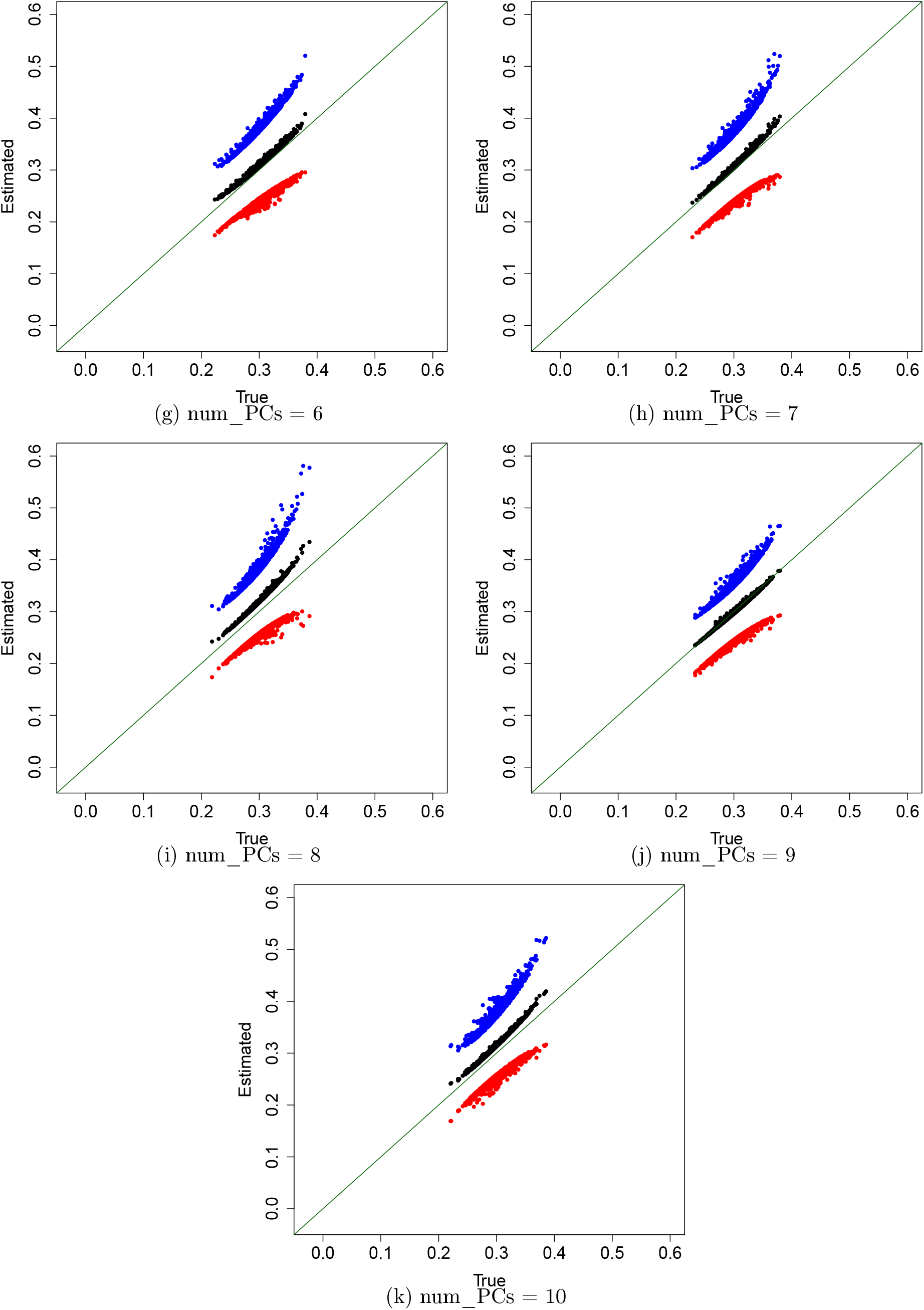
Continues on next page. Comparisons of true conditional heritabilities vs their average estimates (black) +/− 1 empirical S.D. (blue/red) across 100 simulations for each of the *n* = 1000 subjects with varying number of principal components. The true conditional heritabilities are all within 1 empirical S.D. of the estimates when three or more principal components are included. Comparisons of true conditional heritabilities vs their average estimates (black) +/ −1 empirical S.D. (blue/red) across 100 simulations for each of the *n* = 1000 subjects with varying number of principal components. The true conditional heritabilities are all within 1 empirical S.D. of the estimates when three or more principal components are included.

### F.6 Supplementary small-scale simulations

In Appendix G we present supplementary simulations that were conducted at earlier stages of this project. Although these focus on a two population case and use a different simulation framework, our framework for estimating covariate-adjusted marginal and conditional heritabilities is still shown to be valid. Moreover, even though these simulations are less comprehensive in comparison to what those we have presented above, they also illustrate a specific scenario with admixed individuals in which *t*(***s***) and Var(*y*|***s***) are non-linear in ***s*** and use GAMs used to adequately estimate conditional heritabilities.

The following R packages were used in this section: data.table [11] and xlsx [12].

## G Supplementary small-scale simulation study

### G.1 Simulation with East Asian and European samples from the 1000 Genomes Project

We were first interested in testing out our methods under the scenario in which there are two ancestries in a sample without admixed individuals. For this, we used the 1000 Genomes dataset for the SNP matrix but simulated the phenotype.

We used plink 1.9 and 2.0 to do some initial data-handling and obtained SNPs from a contiguous chunk on chromosome 22 from the 1000 Genomes data [7]. To be able to test our method on a dataset consisting of two non-admixed sub-populations, we restricted ourselves to European and East Asian subjects leading to a sample size of *n* = 1007 subjects (503 European subjects and 504 East Asian subjects). We removed SNPs with no variation and mean-centered the remaining ones. This led to a total subset of 14653 SNPs.

Before running the realizations of the simulation study, we calculated the first eigenvector of the in-sample kinship matrix obtained from the 14653 contiguous SNPs, i.e. the first principal component.

Then, for each realization of our simulation we randomly selected *m* = 100 of the 14653 SNPs. The effects of 50 of these were drawn from a standard normal distribution, and the effects of the rest were set to 0.

We simulated the phenotype by using model 3 which is a sub-case of model 1. The values of *γ*_1_ and *γ*_2_ changed in each realization and were obtained by setting them to 10-th and 90-th percentile of (***β***^*T*^ ***x***_1_, …, ***β***^*T*^ ***x***_*n*_) where ***x***_*i*_ represents the observed mean-centered SNPs for the *i*-th subject. This was done to ensure that the population ancestry variables had a strong effect on the phenotype in the simulation regardless of the set of SNPs that were used in the realization.

Moreover *ϵ* was simulated as a random variable distributed as either Normal(0, *σ*^2^(1, 0)) if (*s*_1_, *s*_2_) = (1, 0), or Normal(0, *σ*^2^(0, 1)) if (*s*_1_, *s*_2_) = (0, 1). The parameters *σ*^2^(1, 0) and *σ*^2^(0, 1) were taken such that the underlying conditional SNP-heritabilities described in Equation (4) were *h*^2^(1, 0) = 0.3 and *h*^2^(0, 1) = 0.7 respectively. To do so, the covariance matrix **∑**(1, 0) was set to the sample covariance matrix obtained by restricting to the European samples and the *m* = 100 SNPs. Similarly, the covariance matrix **∑**(0, 1) was set to the sample covariance matrix obtained by restricting to the East Asian samples and the *m* = 100 SNPs. Then, we rearranged Equation (4) to obtain 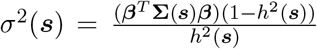, and plugged in the covariance matrices, SNP-effect sizes and desired SNP-heritability to get the necessary values for *σ*^2^(0, 1) and *σ*^2^(1, 0).

We applied three estimation approaches to obtain estimators for the marginal heritability:

1. The first estimation approach (M1), which we called the “naive” approach, consisted of obtaining the Adjusted *R*^2^ and GWASH heritability estimates with our simulated phenotypes and the *m* = 100 mean-centered SNPs without any corrections.
2. The second approach (M2) consisted of obtaining 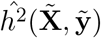 described in Section 2.2, by regressing out ancestry proportions from the *m* = 100 SNPs and the simulated phenotype vector. We used both GWASH and Adjusted *R*^2^ for ĥ^2^.
3. The third estimation approach (M3) was analogous to the second approach except that the first principal component that had been calculated before the simulation was regressed out to obtain 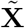 and 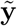. In all cases GWASH was calculated from Equation (A.3) and Adjusted *R*^2^ was obtained using the “summary” function for “lm” objects in R.

We also applied three estimation approaches to obtain estimates for conditional heritability:

- The first approach (C1) consisted of separating out the sample into European and East Asian subjects and obtaining GWASH and Adjusted *R*^2^ estimates separately for both.
- The second approach (C2) involved applying the conditional heritability estimation approach we propose in Section 2.5. For this we took **S** to be a matrix with two columns **S**_1_ and **S**_2_ which represent the proportions of European and East Asian ancestry respectively for each subject. Also, to estimate 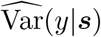 and 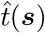 we used the linear regression approach instead of the sub-setting approach. We used both GWASH and Adjusted *R*^2^ to estimate 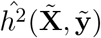.
- The third approach (C3) was analogous to the second approach but involved using principal components instead of ancestry proportions as described in Section A.5.

In all cases GWASH was calculated from Equation (A.3) and Adjusted *R*^2^ was obtained using the “summary” function for “lm” objects in R version 4.0.2. We also calculated the target marginal heritability using Equation (2) taking ***s*** to be distributed as categorical distribution with two categories with probabilities 0.5 each in order to calculate the expectation.

We repeated this procedure 200 times. During each realization we stored the three marginal heritability estimates obtained from approaches M1 - M3, the target marginal heritability, the two conditional heritability estimates obtained from approach C1 (one estimate for Europeans and another one for East Asians) and the two heritability estimates obtained from approach C2 (one estimate for Europeans and another on for East Asians). For approach C3, each individual got their own heritability estimate since we conditioned on principal components and these are continuous variables take different values for each individual. To obtain values that could be compared to the other two conditional heritability estimation approaches, we averaged the heritability estimates for the individuals with European ancestry and also averaged the heritability estimates for the individuals with East Asian ancestry. We then stored these averaged heritability estimates.

It is worth mentioning that C1 serves as a reference to compare the other approaches to, and is not an approach we advocate since in practice a sample might not be as easily separable (specially in the presence of individuals with varying degrees of admixture) - this is an advantage of our proposed framework.

### G.2 Simulation with admixed individuals

We were also interested in testing out our method under the scenario in which there are two ancestries (A and B) in a sample with admixed individuals who we assumed had fifty percent ancestry A and fifty percent ancestry B.

For each repetition of our simulation we generated two vectors (**p**_**A**_ and **p**_**B**_ respectively) of length *m* = 100 populated with realizations of Uniform(0, 1) random variables to represent the allele frequencies of people with only ancestry A and only ancestry B. We then generated a third vector (**p**_**ADM**_) of length *m* = 100 where each entry was the average of the corresponding entries of the other two vectors. This represented the allele frequencies of the admixed individuals.

We simulated the SNP-matrix for 500 individuals with only ancestry A. The matrix had 100 columns total where the j-th column corresponded to the j-th SNP and was populated with realizations of Binomial(2, 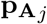) variables where 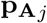 represented the j-th entry of vector **p**_**A**_. We repeated an analogous procedure to obtain SNP-matrices for 500 individuals with only ancestry B and 500 admixed individuals. Then we obtained the sample SNP matrix **X** of dimensions 1500 × 100 by combining the three SNP matrices by rows. Due to this procedure, the first 500 rows corresponded to the SNP values of individuals from population A, the next 500 rows corresponded to SNP values of individuals from population B, and the final 500 rows corresponed to SNP values of admixed individuals. This led to a total of 1500 subjects and 100 SNPs. The columns of this matrix were then mean centered to obtain the matrix **X**.

Then we simulated a vector of SNP effects ***β*** of length 100. We set 50 SNP effects to be drawn from a standard normal distribution, and the effects of the rest were set to 0.

Similar to the previous section, the phenotype was simulated using model 3 but with two differences to the previous simulation. First, the variables *s*_1_ and *s*_2_ from model 3 were used to represent the proportion of ancestry A and B respectively, and so in this setting we had the possibility of ***s*** = (*s*_1_, *s*_2_) = (0.5, 0.5) which was not possible in the previous simulation setting. Second, to account for the third admixed population, we set three conditional heritabilities *h*^2^(1, 0) = 0.3, *h*^2^(0, 1) = 0.7 and *h*^2^(0.5, 0.5) = 0.5 corresponding to individuals from the three different populations respectively using an analogous process as before to generate the pertinent *ϵ* such that *ϵ*|***s*** ∼ Normal(0, *σ*^2^(***s***)). The process to obtain *γ*_1_ and *γ*_2_ was kept the same as the previous simulation.

We then applied the three estimation procedures M1-M3 for marginal heritability described in the previous section, but calculating the first principal component from the in-sample kinship matrix from the simulated SNPs in each realization (instead of using pre-calculating them before the simulation) for procedure M3. We also calculated the target marginal heritability using Equation (2) taking ***s*** to be distributed as categorical distribution with three categories with probabilities 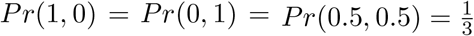 in order to calculate the expectation.

We also applied the three estimation approaches C1-C3 to obtain estimates for conditional heritability described in the previous section with the same modification for calculating principle components mentioned in the above paragraph.

Similar to before, this procedure was repeated 200 times storing the three marginal heritability estimates obtained from approaches M1 - M3, the target marginal heritability, and estimates for the three conditional heritabilities using procedures C1 and C2. As before, for approach C3, each individual got their own heritability estimates and for comparative purposes these were averaged per ancestry group (only ancestry A, only ancestry B and admixed individuals).

After carrying out the simulations we noticed that approaches C2 and C3 were led to biased estimates for the admixed individuals. By taking a look at a single realization of the simulation we noticed that *t*(***s***) and Var(*y*|***s***) did not have a linear relationship ancestry proportions. We therefore repeated this simulation study by modifying the estimation of *t*(***s***) and Var(*y*|***s***) as follows:

- Coding the population ancestry proportions as factors when fitting the linear regression model to estimate *t*(***s***) and Var(*y*|***s***). We called this approach D2.
- Using generalized additive models (GAMs) fitted on the first principal component of the in-sample kinship matrix to obtain estimates for *t*(***s***) and Var(*y*|***s***). This was done using the “gam” function with default parameters from package “mcgv” in R version 4.0.2 (details can be found in [43] [44]). We called this approach D3.

### G.3 Results from simulation study

#### G.3.1 Simulation with East Asian and European samples from the 1000 Genomes Project

**Figure G.15:**
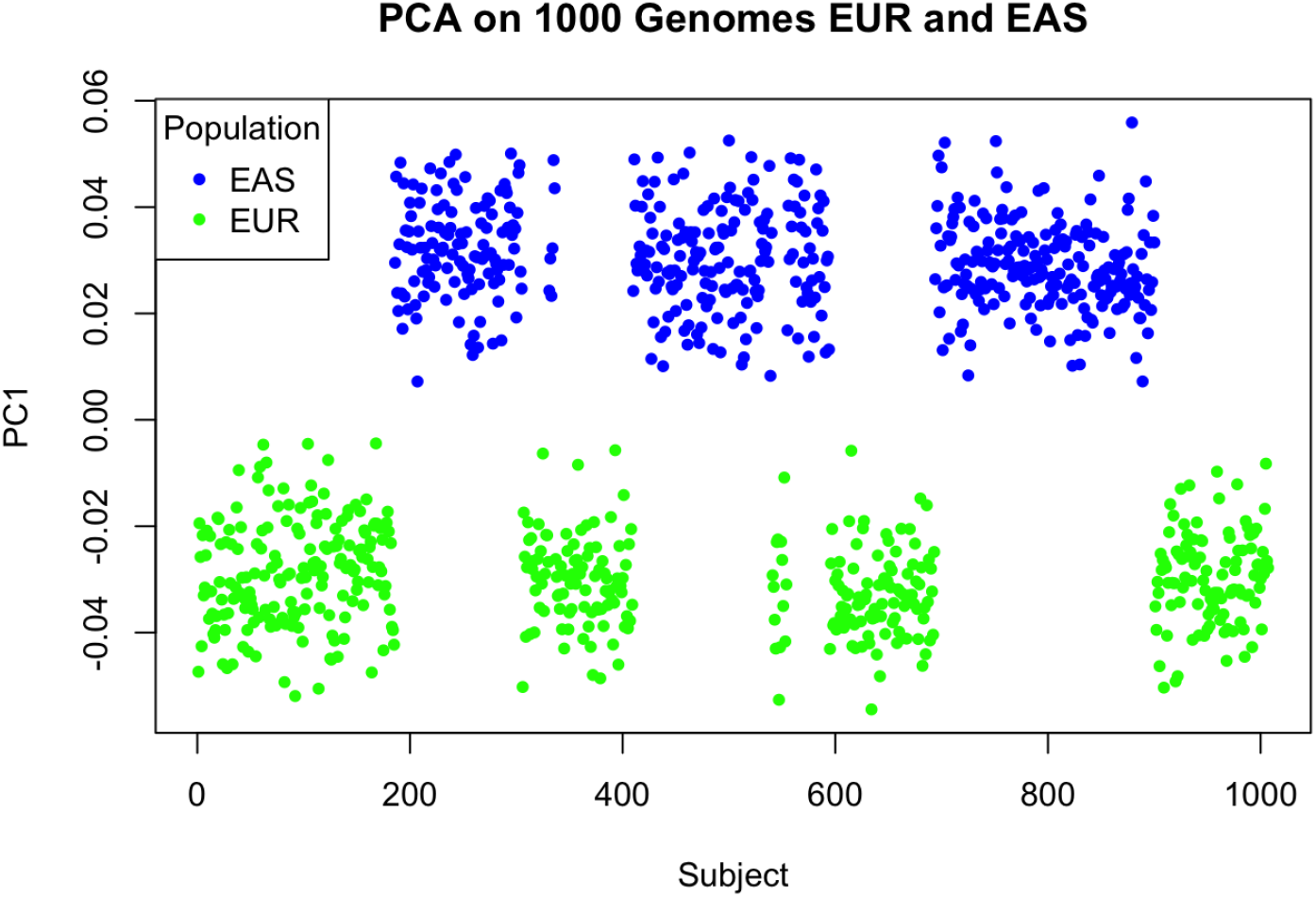
Plot of the principal components calculated for the simulation study of European and East Asian ancestry.

A plot of the principal components calculated for this simulation is shown in Figure G.15. The principal components in these plots are colored by ancestry. As it can be seen, these principal components seem to cluster around 0.03 for subjects with East Asian ancestry and around −0.03 for subjects with European ancestry which matches the theoretical values obtained the two-population equation from [20] and [23]. If we had included more SNPs to calculate the principal components we would expect this clustering to be even tighter.

**Table G.1:** Summary of marginal heritability results for the simulation study with subjects of European and East Asian ancestry. We report the mean and standard deviation in parenthesis across the 200 repetitions. The mean target marginal heritability across repetitions is 0.419 (0.0171). Any mean that is not within one standard deviation of the mean target heritability is colored in red.

| Procedure | Estimator | Mean estimate |
| --- | --- | --- |
| M1) Naïve | Adjusted $R^2$ | 0.602 (0.0725) |
|  | GWASH | 1.57 (0.478) |
| M2) Regressing ancestry proportions | Adjusted $R^2$ | 0.425 (0.0272) |
|  | GWASH | 0.438 (0.0811) |
| M3) Regressing PCs | Adjusted $R^2$ | 0.415 (0.0306) |
|  | GWASH | 0.426 (0.0813) |

Table G.1 summarizes the marginal heritability results from our simulation study. Naiive GWASH in particular has an estimate which exceeds 1, which is clearly inflated since heritability should only take values between 0 and 1. Moreover, we can see that regardless of the framework, the mean of the GWASH estimates tends to be higher than the mean of the adjusted *R*^2^ estimates. Although GWASH has a higher standard error in these simulations than adjusted *R*^2^, we would not be able to use adjusted *R*^2^ in practice due to its limitation of *m* << *n*, and the inclusion of this estimator just helps show that the theory developed holds true for different heritability estimators. Indeed, as can be seen from the table, the average GWASH and adjusted *R*^2^ from procedures M2 and M3 are within one standard deviation of the average target marginal heritability.

**Table G.2:**
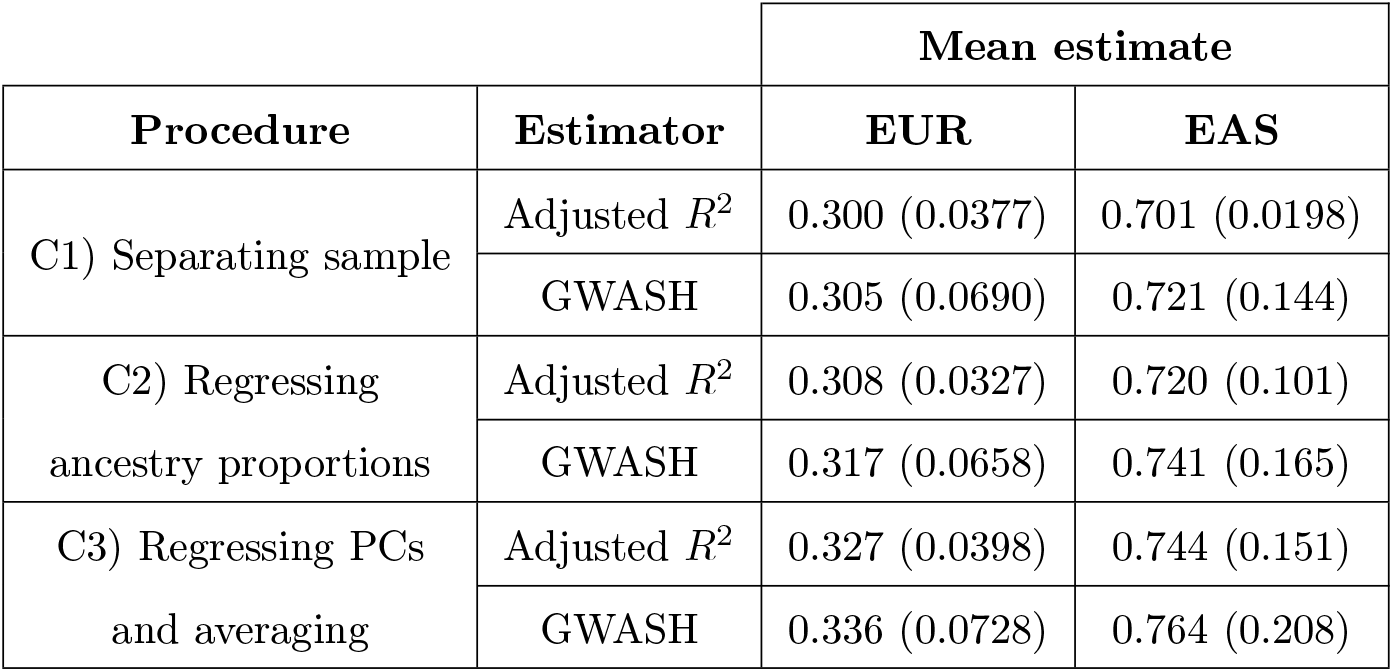
Summary of conditional heritability results for the simulation study with subjects of European and East Asian ancestry. We report the mean and standard deviation in parenthesis across the 200 repetitions. The true simulated conditional heritability is *h*^2^(1, 0) = 0.3 for subjects of European ancestry and *h*^2^(0, 1) = 0.7 for subjects of East Asian ancestry. All estimation procedures and estimators have mean estimates that are within one standard deviation of these quantities.

Table G.2 summarizes the conditional heritability results for our simulation. Moreover, the standard deviation of the GWASH estimates tends to be higher than the standard deviation of the adjusted *R*^2^ estimates which is carried over from the marginal estimates. Furthermore, the standard deviations of the conditional heritability estimates for East Asian subjects seems to generally be higher than for European subjects, with the exception of C1 with Adjusted *R*^2^. This could be attributed to differences in SNP-distributions accross the two populations that affect the covariance matrix for GWASH estimates under procedure C1, and *t*(***s***) for all conditional heritability estimates for procedures C2 and C3.

After carrying out these simulations, the key takeaway is that our general estimation procedure described in Section 2.5 leads to adequate estimates of conditional heritability, comparable separating out the sample. Moreover, replacing ancestry proportions with principal components yields similar estimates under this framework.

#### G.3.2 Simulation with admixed individuals

**Figure G.16:**
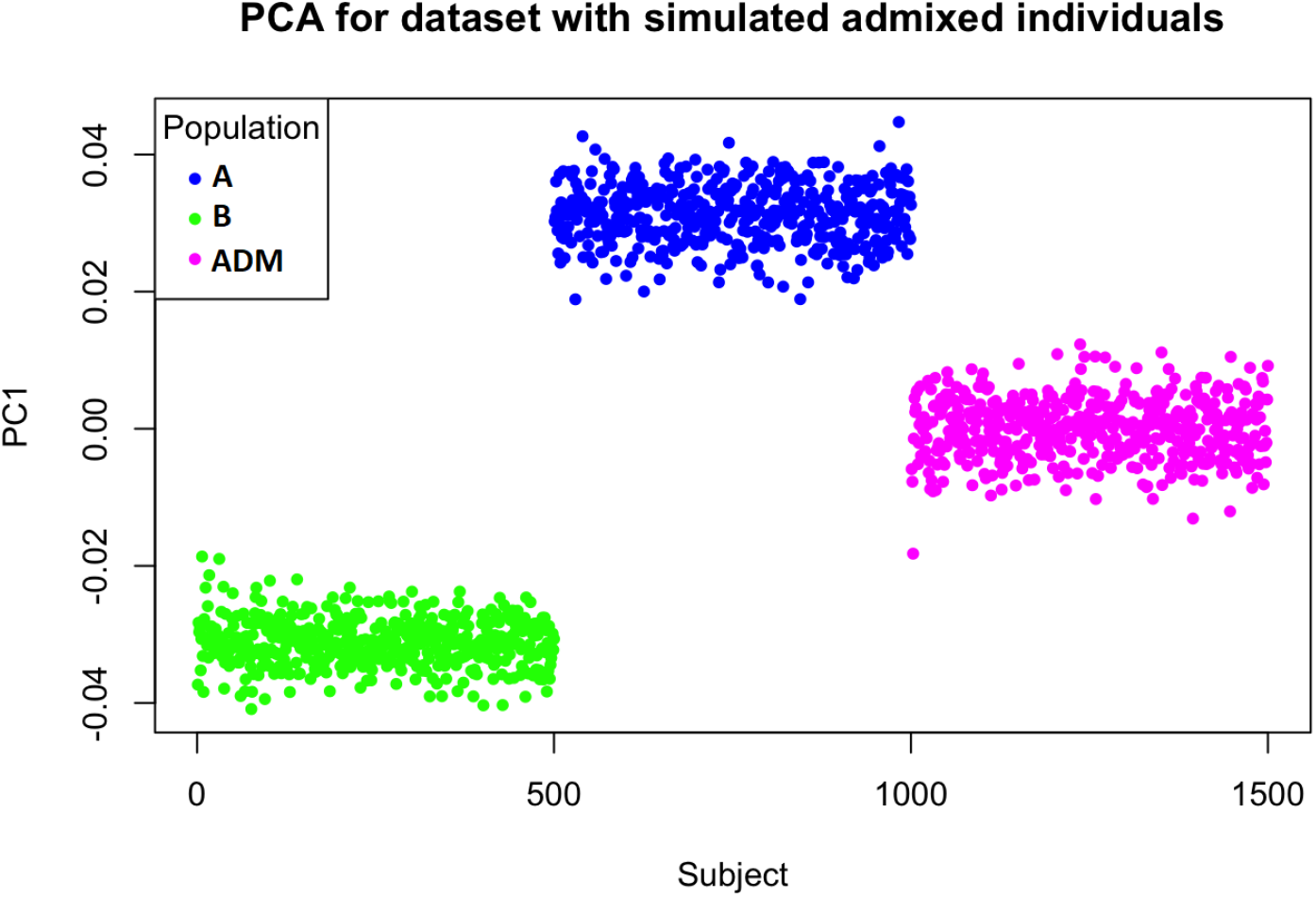
Plot of the principal components calculated for a realization of the simulation study of two ancestries and admixture

A plot of the principal components calculated for a realization of the simulation study is shown in Figure G.16. The principal components in these plots are colored by ancestry. As it can be seen, these principal components seem to cluster around 0.03 for subjects with only A ancestry and around −0.03 for subjects with only B ancestry. The principal components for admixed individuals cluster around 0 which is half way between 0.03 and −0.03. This matches the results from Ma and Amos [19] who illustrate that theoretical principal components for admixed individuals fall between the theoretical principal components of non-admixed individuals in proportion to the admixed individuals’ ancestry compositions (in this case 0 falls half way between −0.03 and 0.03). As mentioned before, if we had included more SNPs to calculate the principal components we would expect this clustering to be even tighter.

**Table G.3:**
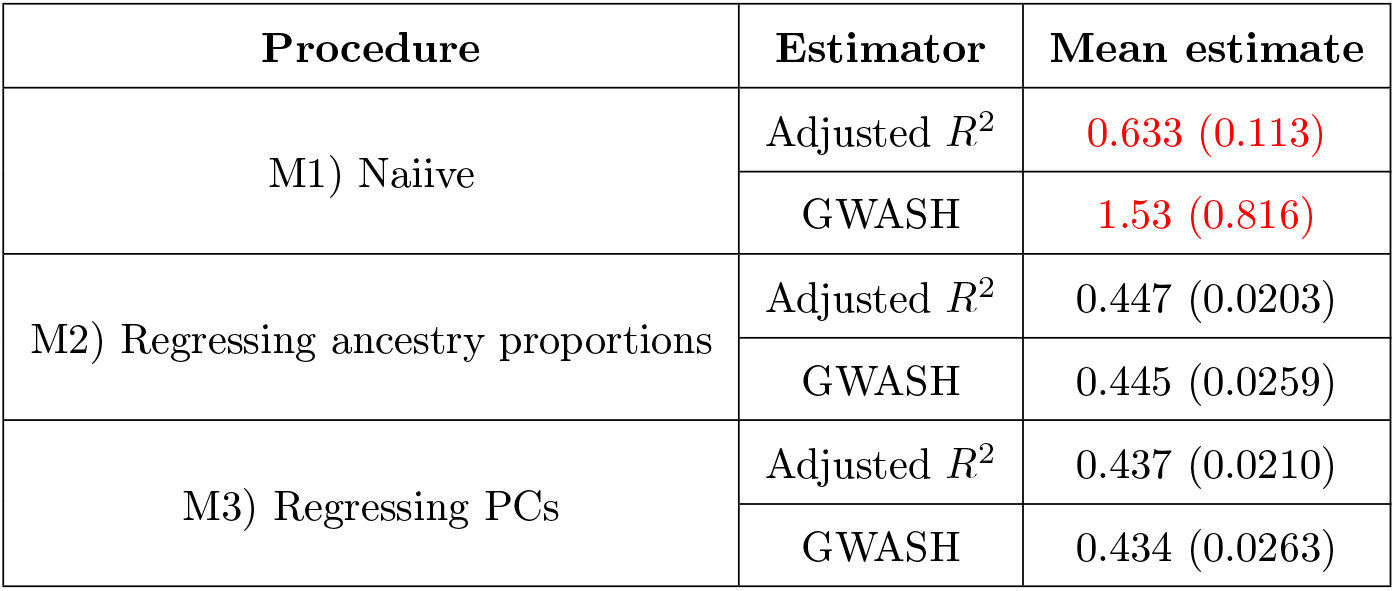
Summary of marginal heritability results for the simulation study of samples with two ancestries and admixture. We report the mean and standard deviation in parenthesis across the 200 repetitions. The mean target marginal heritability across repetitions is 0.448 (0.0101). Any mean that is not within one standard deviation of the mean target heritability is colored in red.

| Procedure | Estimator | Mean estimate |
| --- | --- | --- |
| M1) Naiive | Adjusted $R^2$ | 0.633 (0.113) |
|  | GWASH | 1.53 (0.816) |
| M2) Regressing ancestry proportions | Adjusted $R^2$ | 0.447 (0.0203) |
|  | GWASH | 0.445 (0.0259) |
| M3) Regressing PCs | Adjusted $R^2$ | 0.437 (0.0210) |
|  | GWASH | 0.434 (0.0263) |

Table G.3 summarizes the marginal heritability results from this simulation study. Our findings are analogous to those in the previous section.

**Table G.4:** Summary of conditional heritability results for the simulation study of samples with two ancestries and admixture. We report the mean and standard deviation in parenthesis across the 200 repetitions. The true conditional heritability is *h*^2^(1, 0) = 0.3 for subjects of ancestry only A, *h*^2^(0, 1) = 0.7 for subjects of ancestry only B, and *h*^2^(0.5, 0.5) = 0.5 for admixed individuals. Any mean that is not within one standard deviation of the true conditional heritability is colored in red.

| Procedure | Estimator | Mean estimate |  |  |
| --- | --- | --- | --- | --- |
|  |  | Only A | Only B | Admixed |
| C1) Separating sample | Adjusted $R^2$ | 0.299 (0.0383) | 0.699 (0.0177) | 0.499 (0.0326) |
|  | GWASH | 0.297 (0.0432) | 0.696 (0.0548) | 0.490 (0.0476) |
| C2) Regressing ancestry proportions | Adjusted $R^2$ | 0.322 (0.0275) | 0.749 (0.0928) | 0.448 (0.0204) |
|  | GWASH | 0.321 (0.0310) | 0.745 (0.0949) | 0.446 (0.0259) |
| C3) Regressing PCs and averaging | Adjusted $R^2$ | 0.323 (0.0280) | 0.733 (0.0921) | 0.446 (0.0220) |
|  | GWASH | 0.321 (0.0315) | 0.729 (0.0944) | 0.443 (0.0274) |

Table G.4 summarizes the conditional heritability results for our simulation. For individuals with only ancestry A and only ancestry B, all estimation procedures and estimators have mean estimates that are within one standard deviation of these quantities. However, in the case of admixed individuals, this is only true for procedure C1, and procedures C2 and C3 lead to estimates that are below one standard deviation of the true conditional heritability. To address this, we implemented another estimation procedure and the results are summarized in Table G.5.

**Table G.5:** Summary of conditional heritability results for the simulation study of samples with two ancestries and admixed individuals after modifying the estimation procedure for the conditional variance and trace. As before, the true conditional heritability is *h*^2^(1, 0) = 0.3 for subjects of ancestry only A, *h*^2^(0, 1) = 0.7 for subjects of ancestry only B, and *h*^2^(0.5, 0.5) = 0.5 for admixed individuals. We report the mean and standard deviation in parenthesis across the 200 repetitions. All estimation procedures and estimators in this table have mean estimates that are within one standard deviation of these quantities.

| Procedure | Estimator | Mean estimate |  |  |
| --- | --- | --- | --- | --- |
|  |  | Only A | Only B | Admixed |
| C1) Separating sample | Adjusted $R^2$ | 0.299 (0.0383) | 0.699 (0.0177) | 0.499 (0.0326) |
|  | GWASH | 0.297 (0.0432) | 0.696 (0.0548) | 0.490 (0.0476) |
| D2) Modification to C2<br>for variance and trace | Adjusted $R^2$ | 0.301 (0.0315) | 0.706 (0.0793) | 0.505 (0.0415) |
|  | GWASH | 0.300 (0.0346) | 0.702 (0.0828) | 0.503 (0.0433) |
| D3) Modification to C3<br>for variance and trace | Adjusted $R^2$ | 0.301 (0.0306) | 0.692 (0.0793) | 0.503 (0.0364) |
|  | GWASH | 0.299 (0.0339) | 0.687 (0.0825) | 0.500 (0.0390) |

Table G.5 summarizes the conditional heritability results for our simulation after modifying the estimation procedure for *t*(***s***) and Var(*y*|***s***). Apart from all estimation procedures and estimators in this table having mean estimates that are within one standard deviation of these quantities, procedures D2 and D3 have mean estimates that are closer to the true conditional heritabilities, when compared to C2 and C3 from Table G.4.

After carrying out this simulation, the key takeaway is that our general estimation procedure described in Section 2.5 leads to adequate estimates of conditional heritability, but one should be careful regarding the models used to estimate *t*(***s***) and Var(*y*|***s***). Grouping ancestry proportions into factors before a regression model for *t*(***s***) and Var(*y*|***s***) led to conditional heritability estimates closer to the true values that were comparable to separating out the sample. Moreover using GAMs with principal components to model *t*(***s***) and Var(*y*|***s***) also led to conditional heritability estimates closer to the true values that were also comparable to separating out the sample. If we have ancestry proportion variables with varying degrees of admixture, using GAMs instead of grouping might be a more reasonable approach (for this simulation we could group the admixed individuals because all of them had the same ancestry composition).

## H Application to the ABCD dataset (full analysis)

In this section we will be applying our estimation frameworks on IQ scores obtained from the Adolescent Brain Cognitive Development^SM^ Study (ABCD) (https://abcdstudy.org/about/). Due to its genetic diversity, our methods can be used to obtain novel insights from it. We will first describe the data processing and quality control procedures used for our analysis. We shall then conduct a covariate assessment that illustrates confounding between ancestry and income, as well as relationships between principal components and ancestry. We shall then verify the normality of our phenotype of interest and proceed to estimate marginal and conditional heritabilities. We shall show that there is a decreasing trend on the conditional extrinsicalities 1 − *h*^2^(***s***) with income which has been reported in previous studies.

### H.0.1 Data Description

We were interested in applying the marginal and conditional heritability estimation procedure on a real dataset. For this, we used data from the Adolescent Brain Cognitive Development^SM^ Study (ABCD) (https://abcdstudy.org/about/). Details about the genetic data that we accessed can be found in [13]. We were provided data on age; self reported race/ethnicity (categorized as White, Black, Asian, Hispanic or Other); household income (categorized as “less than $50K”, “greater than or equal to $50K and less than $100K” and “greater than or equal to $100K”); sex (categorized as male or female); proportion of African, European, East Asian and American genetic ancestry; first 10 principal components; and IQ scores. The genetic ancestry variables that were made available to us were previously calculated using “fastStructure” [32], whereas the principal components had been calculated using PC-Air [6] with the GENESIS package [14] (more details about the calculations of these for the ABCD dataset can be consulted in [13]). Ten principal components were used as per the recommendations in [18]. For IQ scores we used the crystallized cognition composite scores described in [16].

### H.0.2 Software used

For this section we made use of both PLINK 1.9 [4] [30] [28] and PLINK 2.0 [4] [29]. During the time of this analysis there were continuous and asynchronous releases for the two. The PLINK 1.9 version for this analysis corresponded to the release from January 16th 2023 for 64-bit Linux, whereas the PLINK 2.0 version corresponded to the release from January 9th 2023 for 64-bit Linux. We used PLINK 1.9 for our quality control steps since the code provided to do so in [21] could only be run in that build at the time of the analysis. We used PLINK 2.0 for running regression models for the GWAS required in this section due to the improvements in speed from its predecessor.

The following R packages from version 4.0.2 were also used in this section: data.table [11], parallel [31], qs [5], mgcv [43] [44], latex2exp [24], tab [40] and ggplot2 [42].

### H.0.3 Selection of individuals and SNPs

We ran some steps to select individuals and SNPs to be able to adequately run our methods. We first removed subjects missing any of the variables mentioned above that were provided for our analysis. Moreover, since the ABCD is a longitudinal study, there were individuals who had multiple observations corresponding to different time points in which they took the test. For such individuals, we only kept the observations pertaining to their first test.

Next we followed some of the quality control steps described by [21] to further restrict individuals included in our analysis which we describe here. We used the available unimputed SNPs and restricted ourselves to the individuals described above. We then removed unimputed SNPs with missingness greater than 20% and then individuals that had more than 5% of the remaining unimputed SNPs missing. We then removed individuals whose genetic sex differed from their reported sex. We followed by removing related individuals determined by a pi-hat score greater than 0.2. This led to *n* = 6623 individuals for further analysis. All these steps were carried out using PLINK 1.9.

It is worth mentioning that for our purposes unimputed SNPs were only used for the selection of individuals as detailed above. For SNPs to include for further analysis we had access to a dataset from the ABCD study with SNPs that had been imputed using the TOPMED imputation server [37]. Specific details regarding this imputation process for the ABCD dataset can be found in [13]. We restricted this dataset to include just the *n* = 6623 individuals from the previous steps using PLINK 2.0. We also used PLINK 2.0 to remove any SNPs with minimum allele frequency less than 5%, any multiallelic SNPs, any SNPs out of Hardy Weinberg Equilibrium (HW p-value less than 10^−10^), any non-autosomal SNPs and any SNPs in the MHC region [8].

We then restricted ourselves to SNPs that were also present in a GWAS analysis carried out by Sniekers et al. for IQ-scores [36] because we were interested in comparing our results. We then randomly selected 750,000 of these SNPs to continue to use for further analysis leading to a final dataset with *n* = 6623 individuals and *m* = 750, 000 SNPs. This random selection of *m* = 750, 000 SNPs was done keeping in mind a comprehensive analysis with different traits carried out in [33] that showed that GWASH heritability estimates tend to converge after *m* = 750, 000 SNPs which can save computational time and resources.

A graphical summary of the steps to select individuals and SNPs is presented in Figure H.17.

**Figure H.17:**
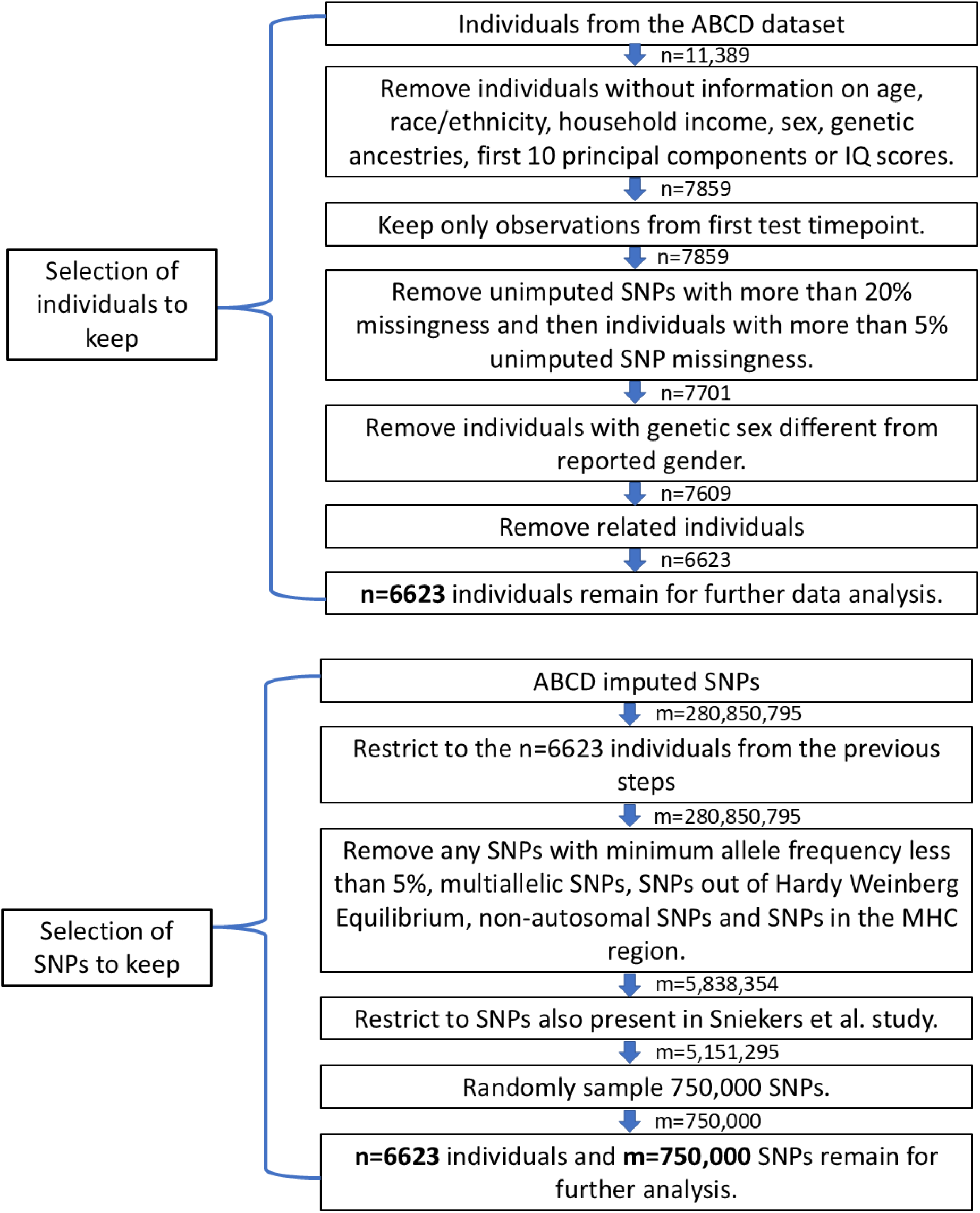
Summary of the individual and SNP selection steps followed for the ABCD data analysis.

### H.0.4 Covariate assessment

Recall that we are using the underlying model described in Equation (1). For our data application our phenotype *y* corresponds to an individual’s IQ-score and the vector ***x*** corresponds to an individual’s SNP values for the 750,000 SNPs kept in the data analysis. Recall that ***s*** represents a vector with covariates of interest that can affect either the phenotype or genotype distribution. For this data analysis our covariates of interest are an intercept term, age, sex (taking a value of 0 for females and 1 for males before centering and scaling), two indicators for household income (one variable representing household income greater than or equal to $50K but less than $100K, and another variable representing household income greater than or equal to $100K) and 10 principal components that had been previously calculated on a larger version of our dataset using PC-Air. Principal components act as proxy variables that capture underlying ancestry. We note that to avoid issues with multicollinearity we do not include the variable representing household income less than $50K since it can be inferred from the values of the other income variables.

It is worth mentioning that the ABCD dataset also has other variables that are associated with ancestry - namely self-reported race/ethnicity (with categories White, Black, Hispanic, Asian and Other) and proportion of European, American, East Asian and African ancestry previously calculated using “fastStructure” [32]. Using principal components as the proxies for ancestry has advantages over the self-reported race/ethnicity variables and the ancestry proportions. For example the article that introduces cov-LDSC [18] has already shown that regressing out principal components helps obtain estimates of adjusted marginal heritability for admixed populations and it recommends using 10 of these in practical settings. Regressing out self-reported race/ethnicity might not be granular enough to capture the variability within the “Hispanic” and “Other” categories. Moreover, we only have estimates for four ancestry proportions and there might be more ancestral variability that might need to be captured (e.g. differences in individuals from North European and South European ancestry). Using principal components might capture these more granular differences. Furthermore self-reported ancestry/ethnicities may be influenced by culture and not accurately reflect genetic ancestry.

We made some additional plots to further justify our use of principal components. From plots of principal components against each other colored by self-reported race/ethnicity (Figure H.18) we can see that principal components can help distinguish between groups quite well. For example, in the plot of the first principal component vs the second, we can see that individuals identifying as “Black” tend to have higher values for the first principal component and individuals identifying as “White” tend to have lower values for the second principal component. Moreover, with the plot of the second principal component vs the third we can see that individuals identifying as Hispanic tend to have higher values for the third principal component than other individuals.

Figures H.19, H.20, H.21 and H.22 help illustrate the relationship between principal components and the four ancestry proportions. For example, we can see from Figure H.19 that the 1st principal component is highly correlated with African Ancestry. Moreover we can also see from these plots that the PCs might capture more granularity than European, Hispanic, African and East Asian. This can be seen in Figure H.20 where the fourth principal component seems to take a markedly wider range of values for people with high European genetic ancestry. This suggests that the fourth principal component might be capturing genetic diversity within people of European ancestry.

As an aside, one interesting thing to notice from Figures H.20 and H.21 is that there is a fair amount of admixture within individuals self-reported as Asian since we see several individuals with more than 40 percent European ancestry in this category. Moreover, historically described admixture within subjects self-reported as Hispanic can be observed with individuals in this category having a wide range of European and American ancestry proportions (Figures H.20 and H.22), as well as in individuals self-reported as Black who have a large range of African and European ancestry proportions (Figures H.19 and H.20). Finally, individuals self-reported as “Other” have a scattered range and combination of ancestry proportions, but most seem to have at least 40 percent European ancestry (Figure H.20).

All this said, self-reported race/ethnicity is a good variable to use to summarize findings and relationships between variables. For example, as it can be shown from Figure H.23, there is confounding between household income and self-reported race/ethnicity with certain groups reporting a higher household income than others. We shall also use this variable later to group results from the application of our method.

**Figure H.18:**
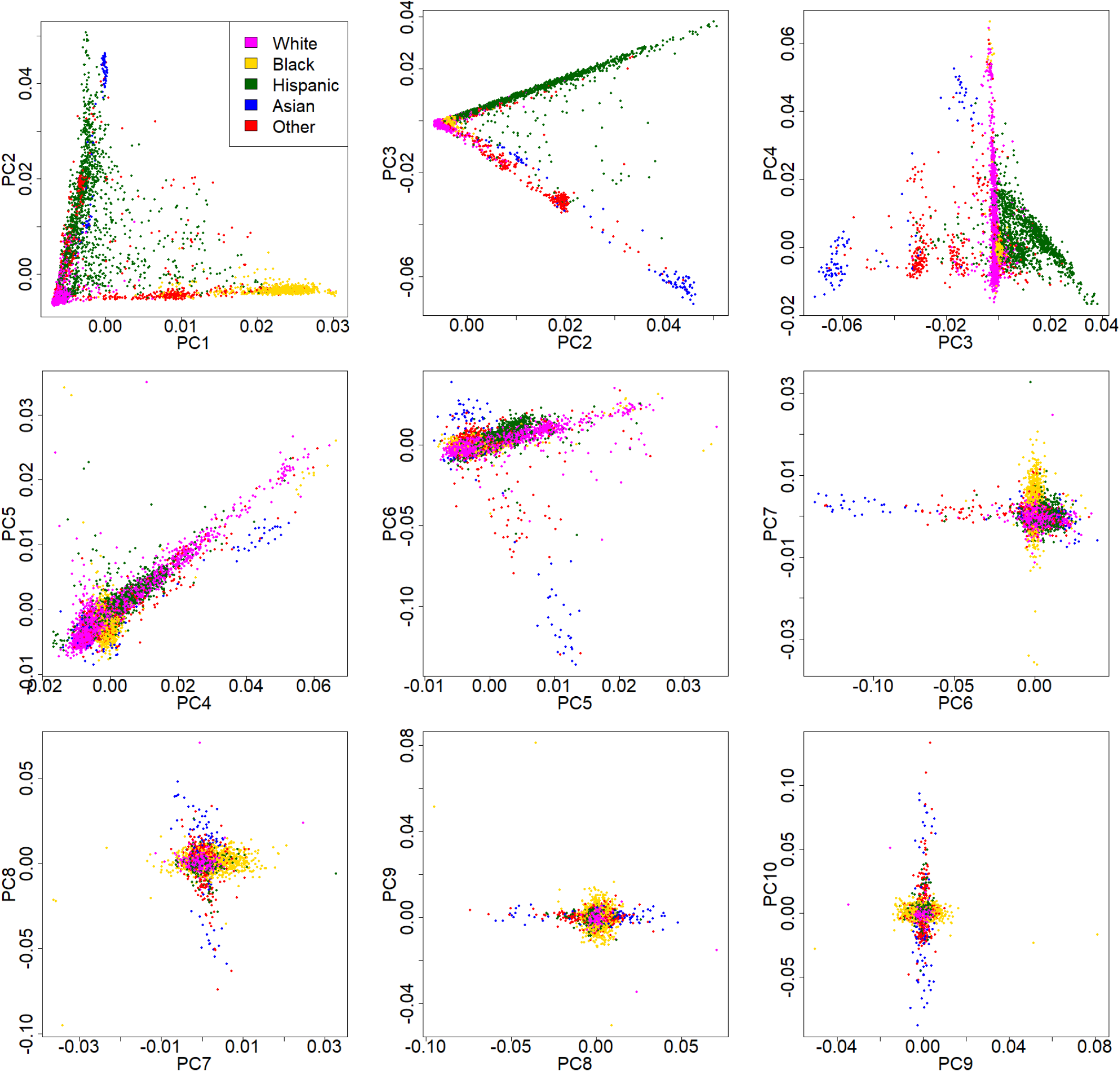
Plots of principal components against each other for individuals included from the ABCD dataset colored by self reported race/ethnicity.

**Figure H.19:**
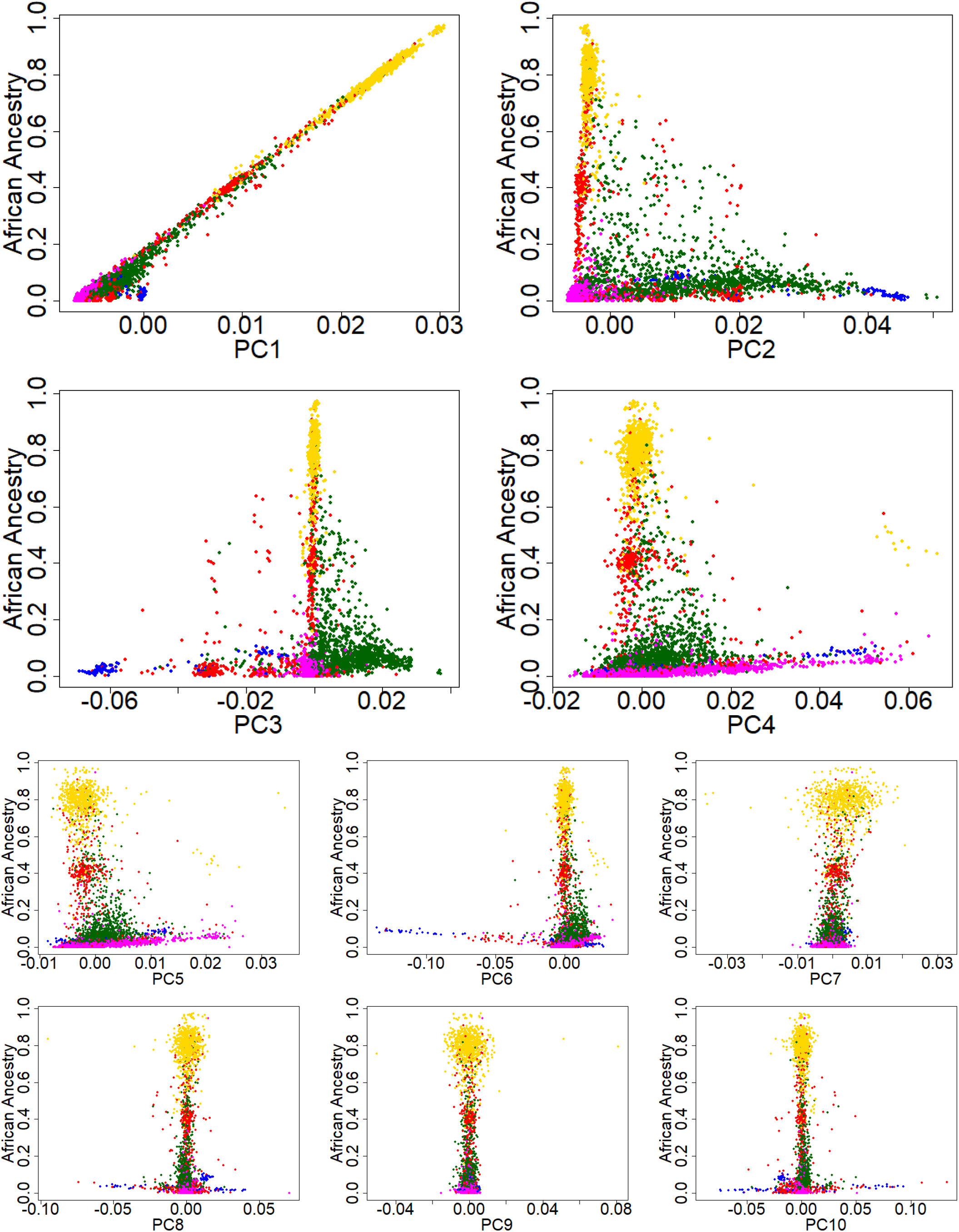
Plots of African genetic ancestry proportion vs PCs for individuals included from the ABCD dataset colored by self reported race/ethnicity (Black - yellow, White - magenta, Hispanic - green, Asian - blue, Other - red).

**Figure H.20:**
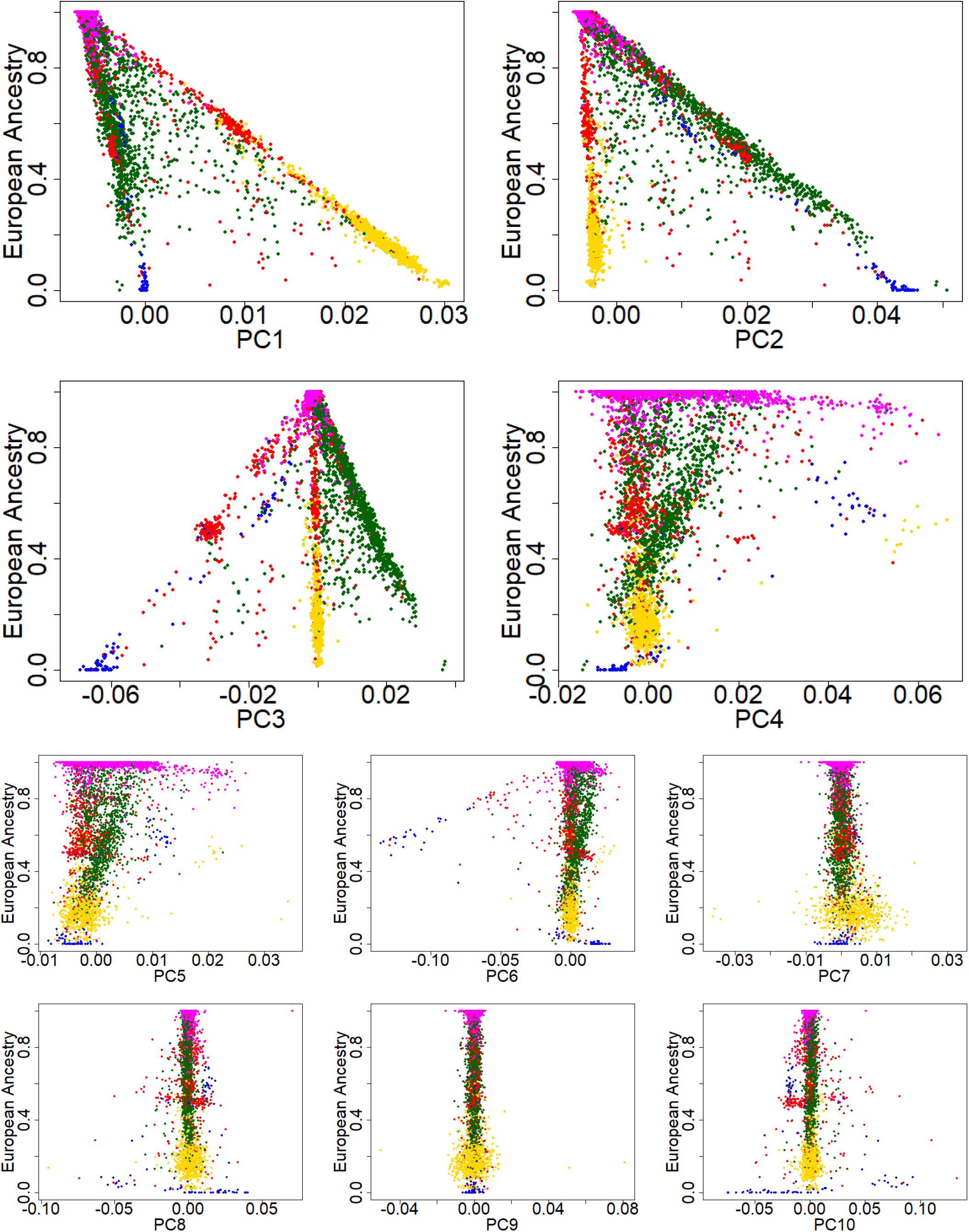
Plots of European genetic ancestry proportion vs PCs for individuals included from the ABCD dataset colored by self reported race/ethnicity (Black - yellow, White - magenta, Hispanic - green, Asian - blue, Other - red).

**Figure H.21:**
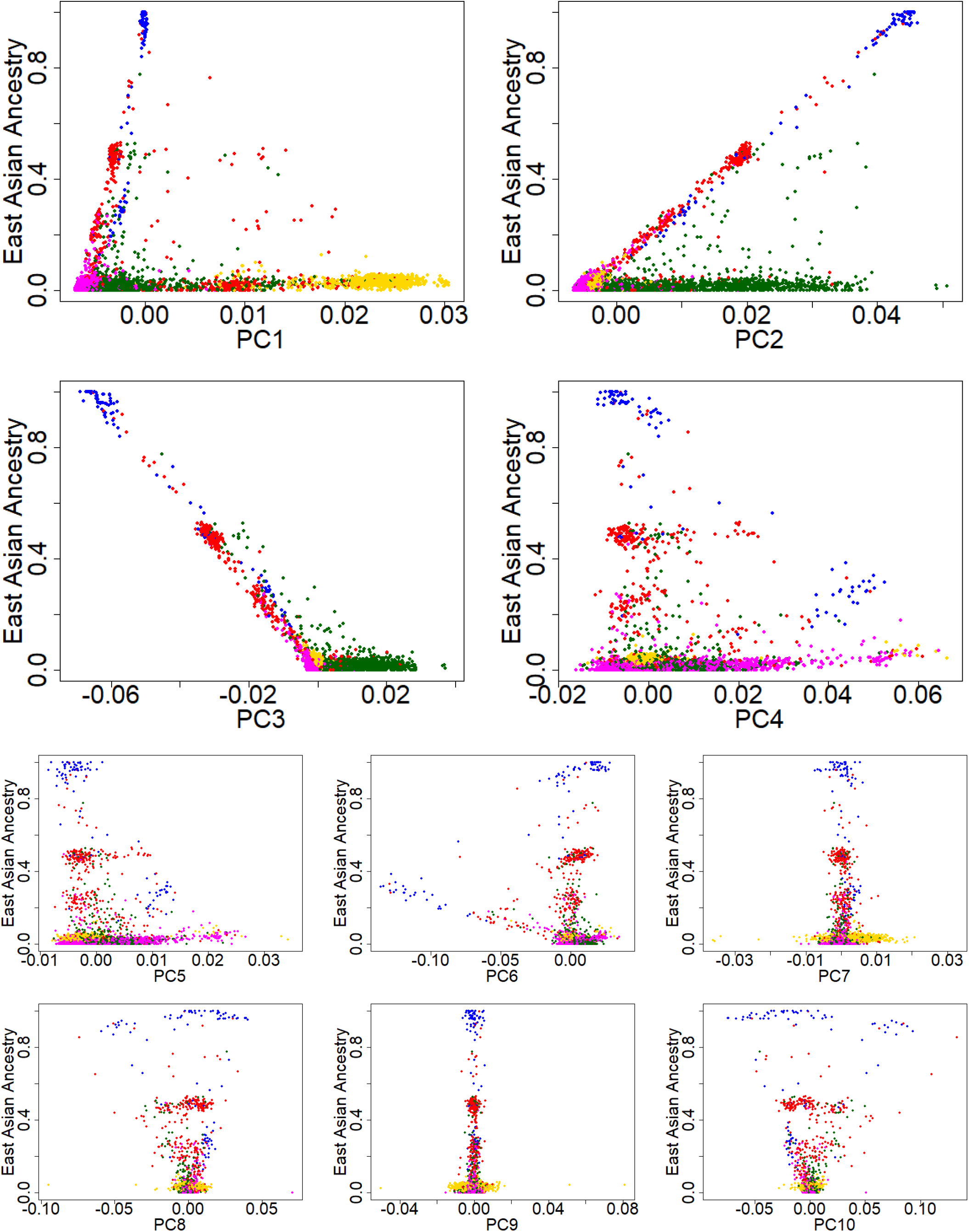
Plots of East Asian genetic ancestry proportion vs PCs for individuals included from the ABCD dataset colored by self reported race/ethnicity (Black - yellow, White - magenta, Hispanic - green, Asian - blue, Other - red).

**Figure H.22:**
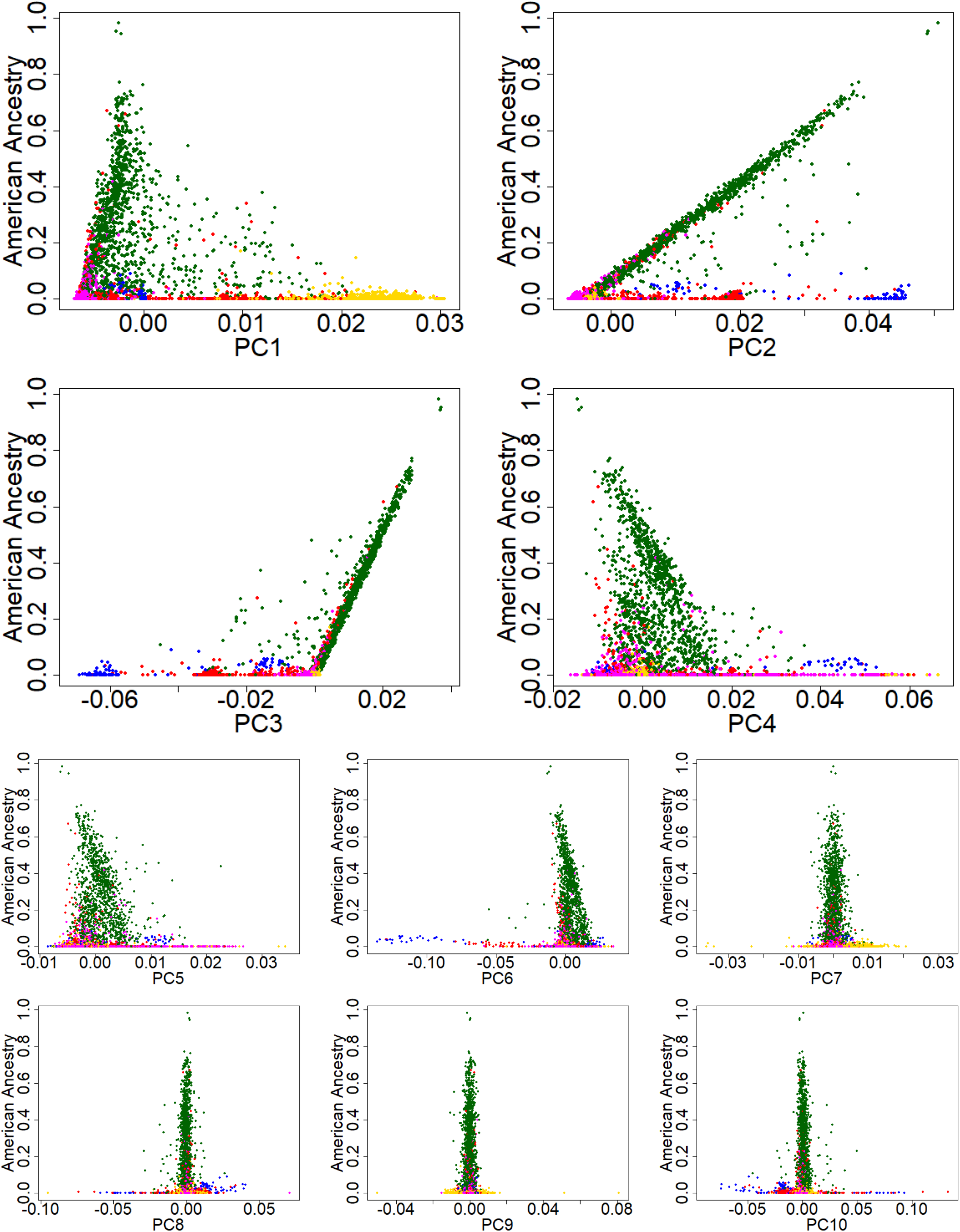
Plots of American genetic ancestry proportion vs PCs for individuals included from the ABCD dataset colored by self reported race/ethnicity (Black - yellow, White - magenta, Hispanic - green, Asian - blue, Other - red).

**Figure H.23:**
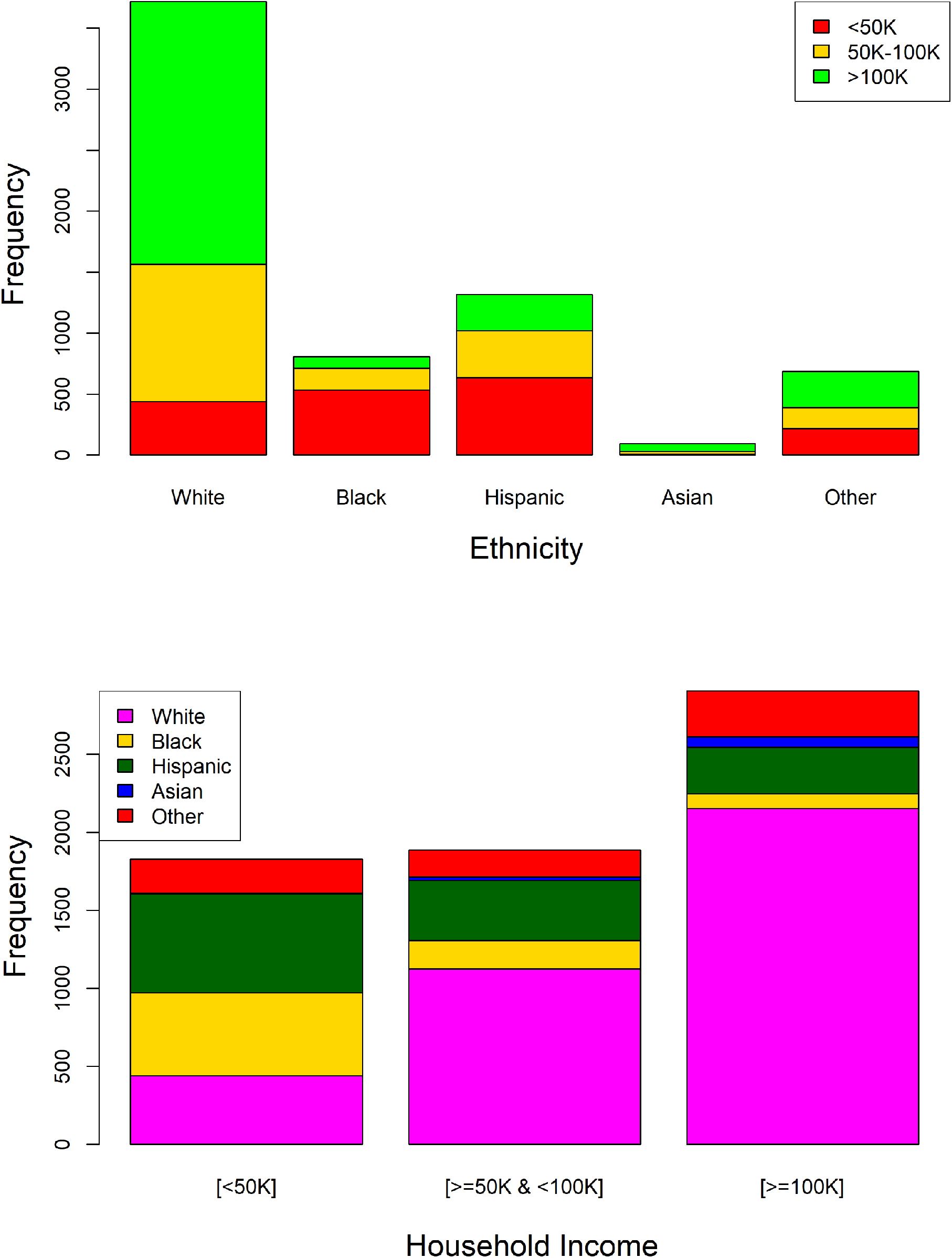
Stacked barplots showing confounding between income and self reported race/ethnicity.

### H.0.5 Checking normality assumptions

We made histograms and QQ-plots of the IQ scores and their residualized version (regressing out the covariates and extracting the residuals). These are shown in Figure H.24. Although these variables deviated slightly from a normal distribution, they seemed to be close enough to carry on our analysis without transformations.

**Figure H.24:**
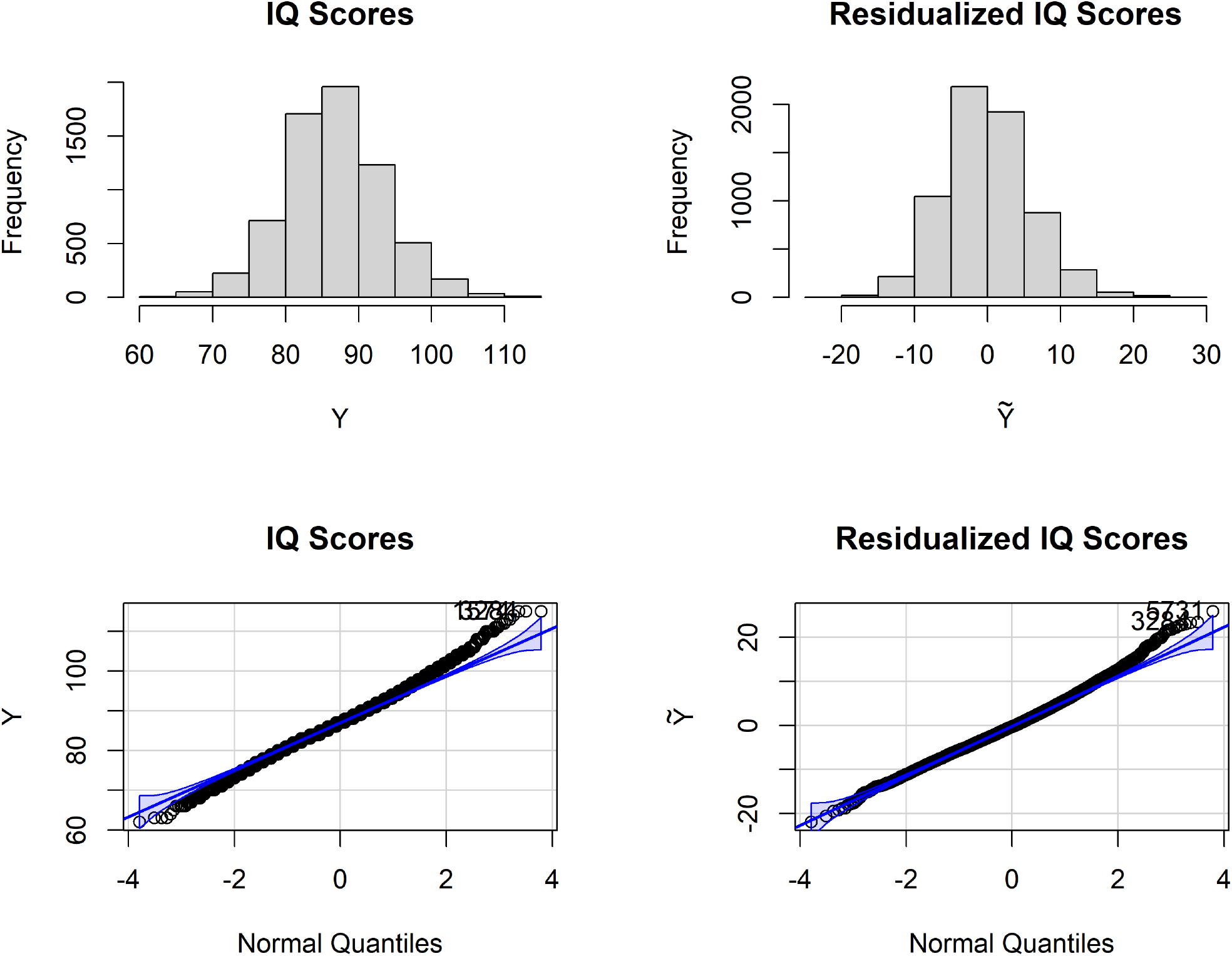
Histograms and QQ-Plots to evaluate the normality of IQ Scores and their residualized version after regressing out covariates.

### H.0.6 Marginal heritabilities

To calculate adjusted and unadjusted LD-scores, 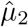 and 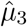 we first took **X** to be the matrix of the 750,000 SNP values for the subjects in the ABCD dataset and obtained 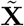 as discussed in Section 2.5 by residualizing on the 15 covariates of interest defined in Section H.0.4. We calculated the unadjusted LD-scores using the approximation from Equation (A.2) taking 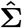 to be the sample correlation matrix obtained from **X** and *Q*_*j*_ to be the set of all SNPs that are in the same chromosome as SNP *j*. We also used this equation to calculate the adjusted LD-scores, but setting 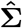 to be the sample correlation matrix obtained from 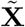 instead of **X**. Next,we obtained the unadjusted 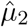by plugging in the unadjusted LD-scores into the approximation from Equation (A.7). The adjusted 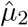 was obtained analogously by plugging in the adjusted LD-scores into this equation. The resulting unadjusted and adjusted 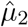 were 68.27 and 16.29 respectively. We also approximated 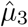using the weighted average of per-chromosome estimates obtained described in Section S4 of the Supplementary Material of [33]. The resulting unadjusted and adjusted 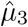were 73976.32 and 942.01 respectively. The reason that the adjusted values for 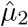 and 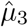are much lower than the unadjusted ones can be explained by the residualization on covariates removing correlation between SNPs that is caused by underlying population structure and admixture in the sample.

To calculate squared correlation scores, we first ran PLINK 2.0 to conduct a GWAS for the IQ scores with our subset of 750,000 SNPs covarying for the 15 covariates of interest defined in Section H.0.4. We also downloaded the GWAS summary statistics provided by Sniekers et al. [36]. To obtain the squared correlation scores 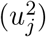 for the GWASH estimator, we transformed the Sniekers et al. [36] GWAS summary statistics with Equation (A.5) and ABCD GWAS summary statistics with Equation (C.1).

To obtain the GWASH estimate for marginal heritability we plugged in the adjusted 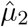 and squared correlation scores 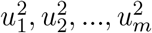 into Equation (A.3). We also ran LDSC-1 using the software provided by [3] with the summary statistics from the ABCD GWAS and Sniekers et al. GWAS, and our adjusted LD-scores obtained from the ABCD dataset. We also obtained the Naive 2 estimates described in Section F.2. As described in Section F.2, Naive 2 estimates are the most likely to be computed in practice if no further adjustment is made. Standard errors for the GWASH estimates were obtained as described by Equation (A.4) and for the LDSC-1 estimates were obtained from the output generated by the software pertaining to [3]. We summarize our results in Table H.1.

**Table H.1:** Covariate adjusted and Naive 2 marginal heritability estimates for the ABCD and Sniekers et al. IQ data. GWASH and LDSC-1 give similar estimates, but the estimated heritability is higher in the ABCD dataset than the Sniekers et al. dataset.

|  | ABCD |  | Sniekers et al. |  |
| --- | --- | --- | --- | --- |
|  | Adjusted | Naive 2 | Adjusted | Naive 2 |
| GWASH | 0.295<br>(0.052) | 0.070<br>(0.034) | 0.165<br>(0.007) | 0.039<br>(0.006) |
| LDSC-1 | 0.302<br>(0.049) | 0.071<br>(0.012) | 0.187<br>(0.008) | 0.049<br>(0.002) |

As can be seen from Table H.1, the GWASH and LDSC-1 estimates are similar to each other. In the literature, the heritability of intelligence is estimated to be between 25% to 50%, with twin studies estimating it closer to 50% and SNP-heritability studies closer to 25% [26]. Our covariate-adjusted marginal heritability with the ABCD dataset is within this ballpark, whereas the corresponding Sniekers et al. estimate is below this. This is expected because Sniekers et al. acknowledge that heritability estimated from their study is understimated [36], and the crystalized metric used to measure intelligence for the ABCD dataset has also been previously shown to be strongly associated with polygenic predictors [16].

Moreover, the Naive 2 estimates are below this too, though we would caution regarding the interpretation of this for the Sniekers et al. data. We use the same adjusted and unadjusted 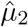and LD-scores for the ABCD and Sniekers et al. dataset to generate Table H.1 for comparative purposes. However, it is worth keeping in mind that these datasets come from distinct population types with the ABCD dataset being more heterogenous and admixed, and so the underlying 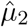 and LD-scores could differ between the two. In practice, if we had access to covariates and genotypes from the Sniekers et al. study we would expect the covariate-adjusted and Naive 2 frameworks to give similar estimates for the Sniekers dataset.

That said, Sniekers et al. [36] estimate the SNP-heritability for their dataset using LDSC and obtain a heritability of 0.20 (S.E.= 0.01) with 12,104,294 SNPs which is within two standard errors of the estimated heritability of 0.187 that we obtain from using the adjusted LD-scores from the ABCD dataset restricting to 750,000 SNPs. Furthermore, when we use an alternative 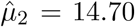 corresponding to 750,000 SNPs from the 1000 Genomes European sample [7] we obtain an estimated GWASH heritability of 0.183 for the Sniekers et al. dataset which is not too different from the corresponding heritabilities in Table H.1. Details to obtain this recalibrated 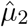 can be found in Appendix I.

#### H.0.7 Conditional heritabilities and extrinsicalities

Next, we estimated conditional heritabilities as described in Section 2.5. Since covariate adjusted GWASH and LDSC-1 give similar estimates for marginal heritability (Table H.1), we used covariate adjusted GWASH as the estimate for 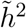. We tested the sensitivity of our approach to using using generalized additive models (GAMs) instead of linear models (LMs) for obtaining the estimated trace parameter 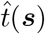 and variance parameter 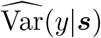. Both types of models were fit on the covariates of interest defined in Section H.0.4. This was done using the “gam” function with default parameters from package “mcgv” in R version 4.0.2 (details can be found in [43] [44]) and the “lm” function in R version 4.0.2. Since our covariates included principal components, which are a continuous variables that differ for each individual, we obtained estimates for each subject in our dataset.

There was high correlation between the estimates from the two approaches in the dataset. The estimates for the trace parameter *t*(***s***) had a Pearson correlation coefficient of 0.947 and the estimates for the variance parameter Var(*y*|***s***) had a Pearson correlation coefficient of 0.848. Moreover the conditional heritability estimates with GAM and LM models coincided for the most part too as can be seen in Figure H.25. They had a Pearson correlation coefficient of 0.926 and there were only 4 out of 6623 observations with estimates differing by more than 0.1. These correspond to those with outlier values in the seventh principal component as shown in Figure H.26. Although GAMs are more flexible models, one would expect them to be more sensitive to outliers due to their additional complexities. Thus we proceed with the linear model estimates since these make use of a more parsimonious modeling framework and generally coincide with GAM estimates too.

**Figure H.25:**
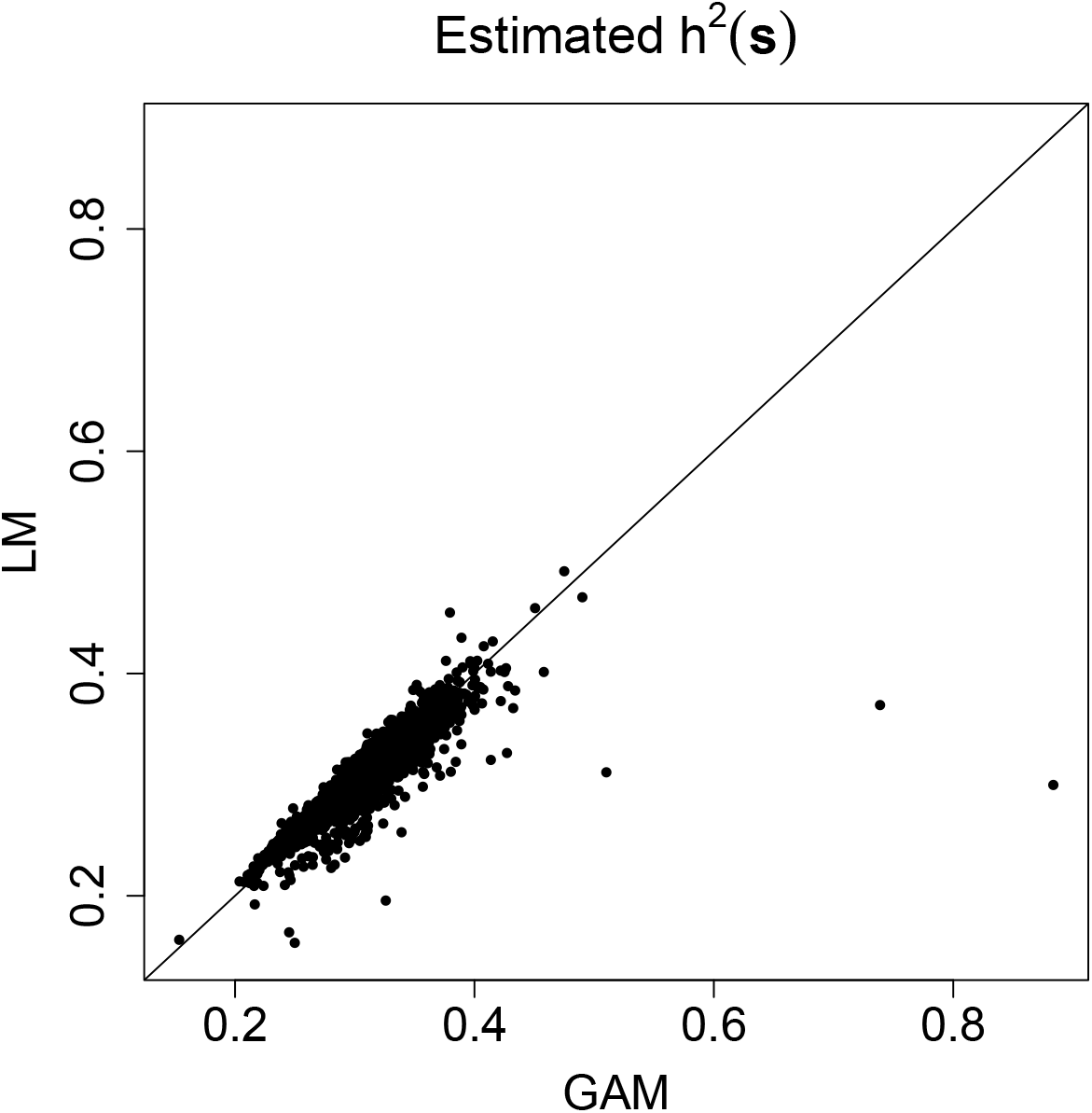
LM vs GAM conditional heritability estimates. The estimates coincide for the most part with only four pairs differing by more than 0.1.

**Figure H.26:**
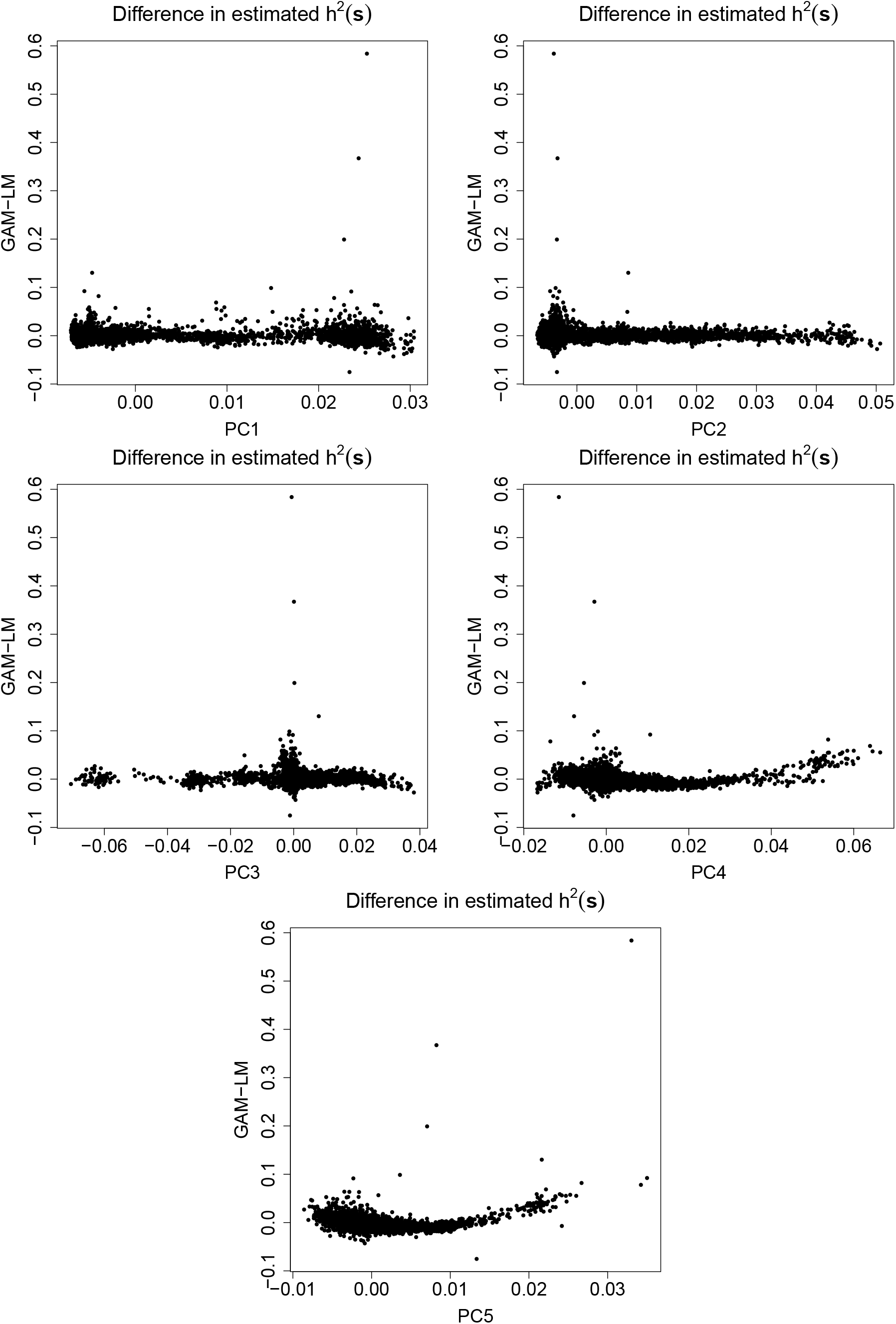
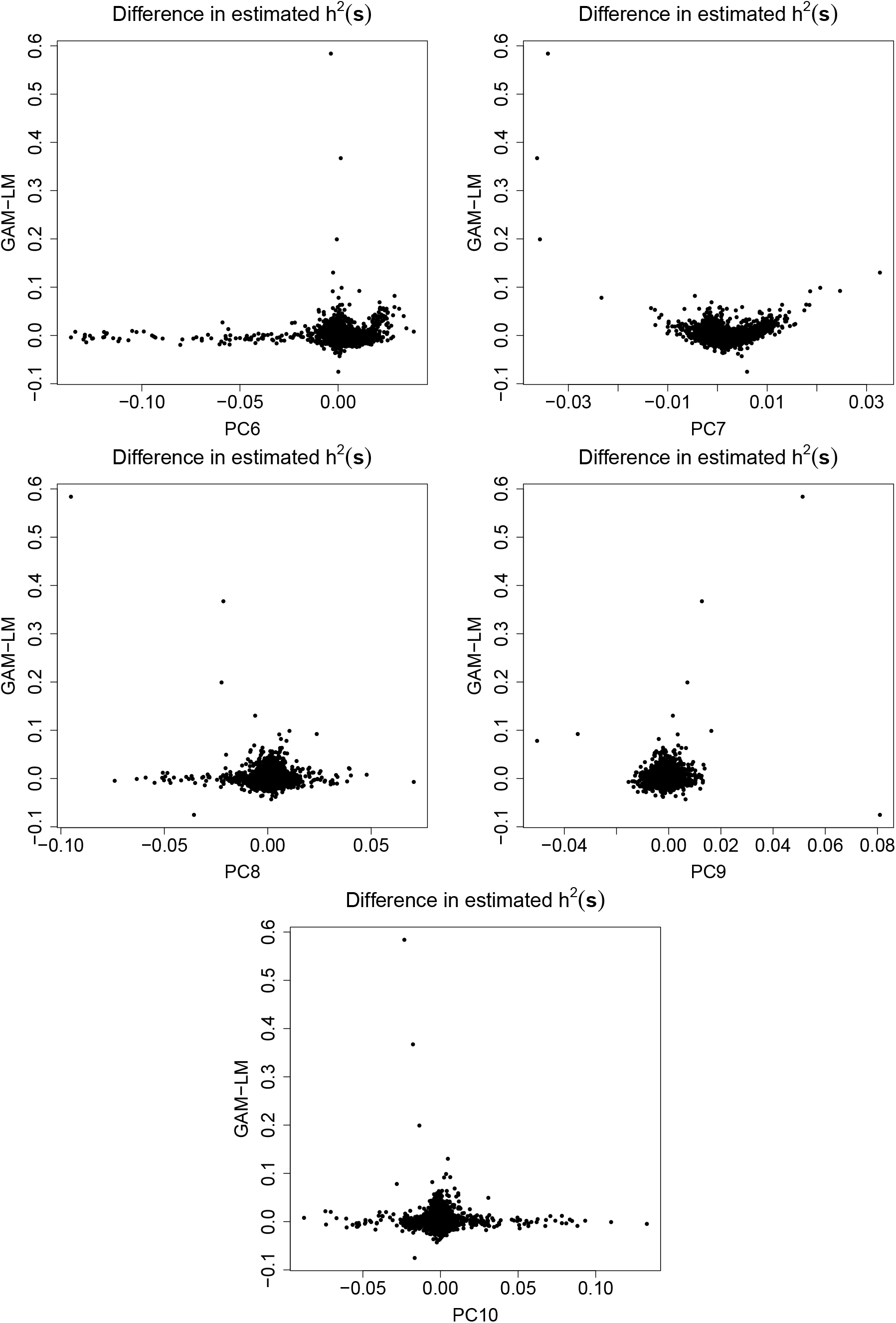
Continues on next page. Scatter plots of the difference between GAM and LM conditional heritability estimates vs principal components. The estimates that differ the most between the GAM and LM frameworks correspond to outliers in the seventh principal component. Scatter plots of the difference between GAM and LM conditional heritability estimates vs principal components. The estimates that differ the most between the GAM and LM frameworks correspond to outliers in the seventh principal component.

There was no discernible difference in the distribution of the estimated conditional heritabilities obtained when comparing male and female subjects as shown in Figure H.27. The distribution of estimates for 1 − *h*^2^(***s***), Var(*y*|***s***), *τ* ^2^(***s***) and *σ*^2^(***s***) separated out by income and ethnicity are illustrated in Figure H.28. The estimates for parameters *τ* ^2^(***s***) and *σ*^2^(***s***) are obtained as described in Section 2.6. We decide to plot the estimated extrinsicalities 1 − ĥ^2^(***s***) instead of the estimated heritabilities 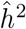(***s***) since non-genetic factors are more pertinent to the differences in subjects in our dataset. This is because, as shown in Figure H.28, we observe an inverse trend between phenotypic variance Var(*y*|***s***) and income, which is most influenced by the non-genetic variance *σ*^2^(***s***) instead of the genetic variance *τ* ^2^(***s***).

To further assess the precision of our conditional heritability estimates, we extracted the median extrinsicalities from each income-ethnicity group and plotted them along with their corresponding estimates for standard errors using the square-root of Equation (E.1). We also plotted extrinsicalities plus/minus one estimated standard error for each subject in the ABCD data analysis in a scatterplot. This corresponds to Figures H.29 and 3.4b. From these figures, we notice that that the standard errors are highest when the estimated extrinsicality is close to 0.5 and then decrease with increasing extrinsicality. This is similar to the behaviour of the GWASH estimator for homogenous samples [33]. Moreover, the estimated standard errors seem to be generally higher in the Asian subgroup and lowest in the White subgroup, which can be explained by the difference in number of subjects for each of these (Figure H.23). Although there is an overlap in the standard error bars for the median conditional extrinsicalities for each income-ethnicity group in Figure H.29, when looking at all subjects in Figure 3.4b we notice conditional extrinsicalities outside one standard error of each other, specially when comparing the highest ones to the lowest ones. Future work is required for implementing a hypothesis testing framework and making any statements on the statistical significance of these differences, but from a descriptive perspective these methods provide useful insights about the potential contribution of the genetic and environmental differences on a trait of interest at a granular level.

**Figure H.27:**
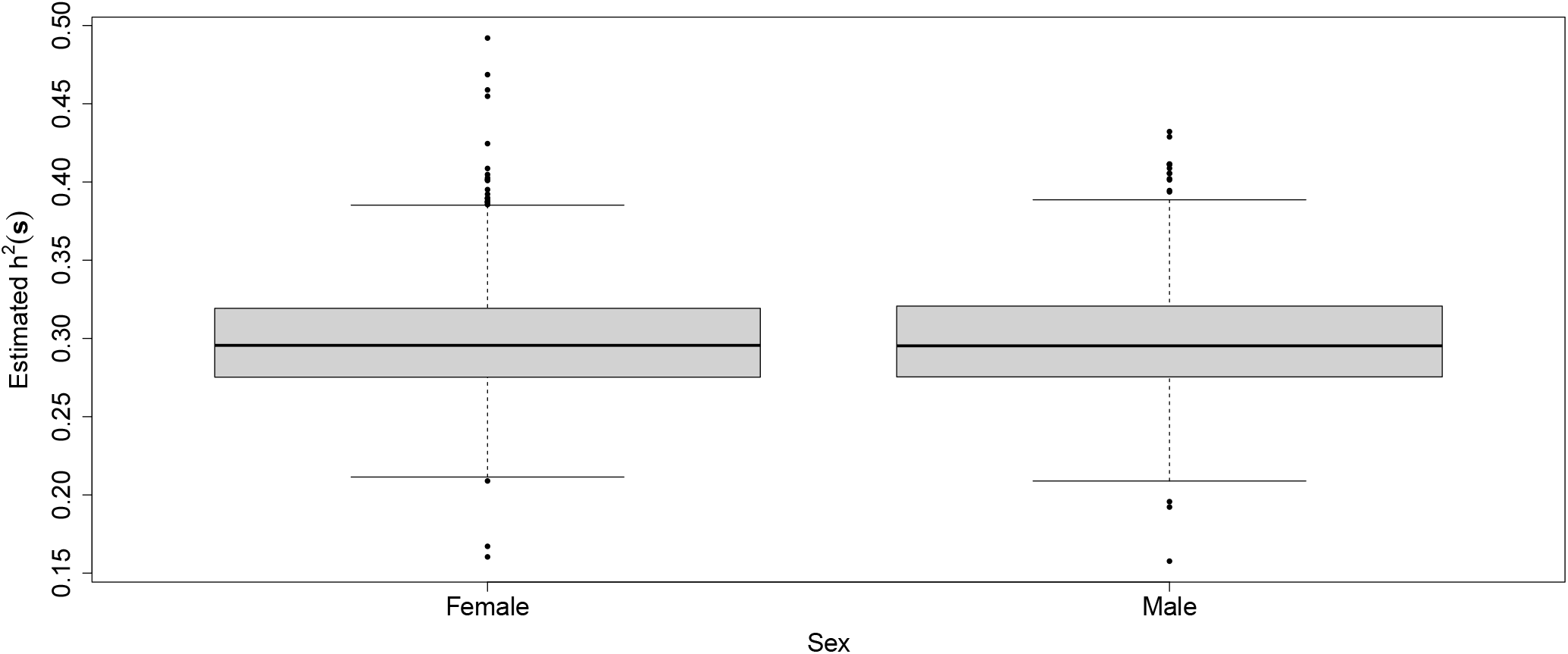
Boxplots of estimated *h*^2^(***s***) grouped by sex. There are no discernible difference in the distribution of the estimated conditional heritabilities.

**Figure H.28:**
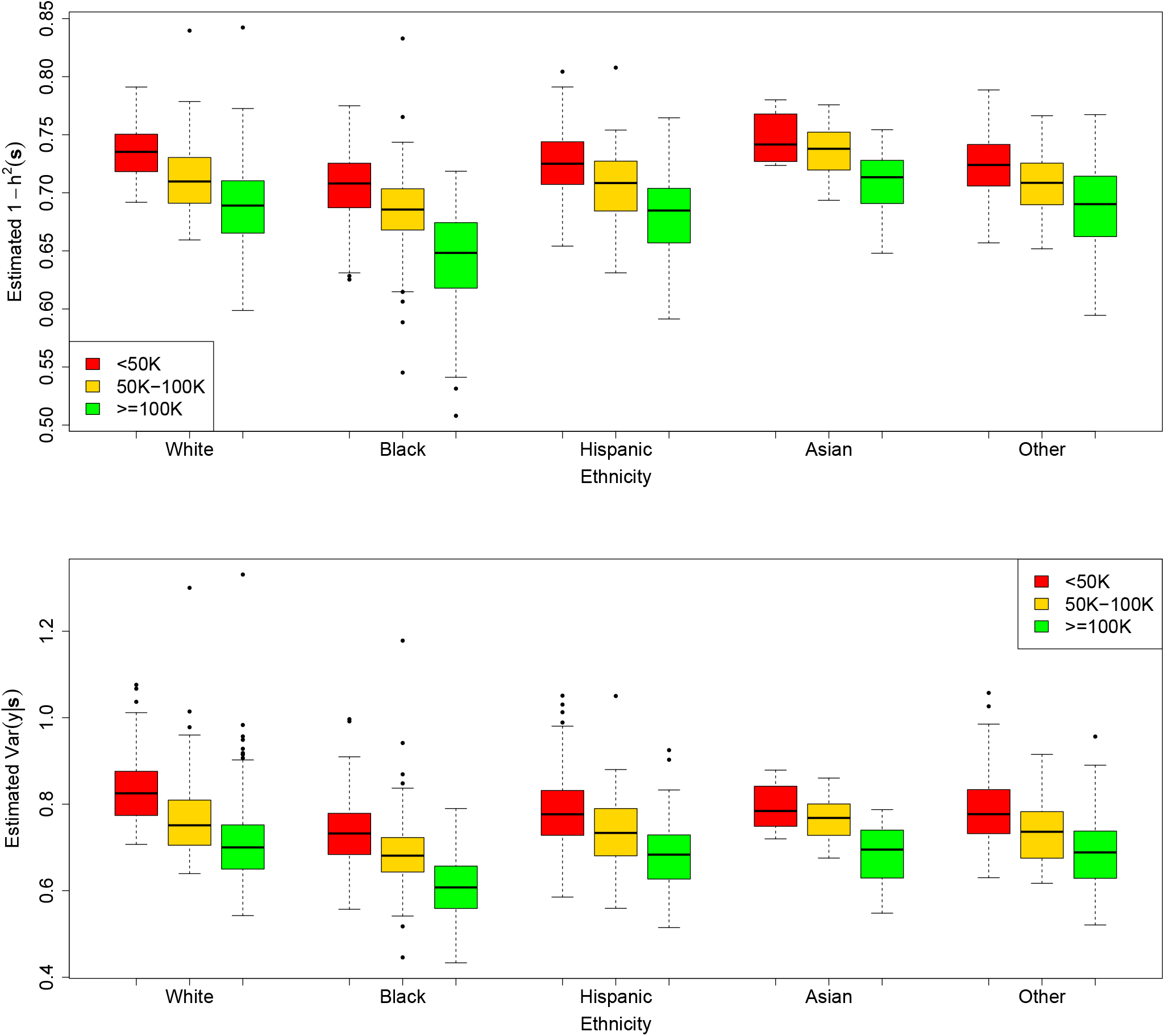
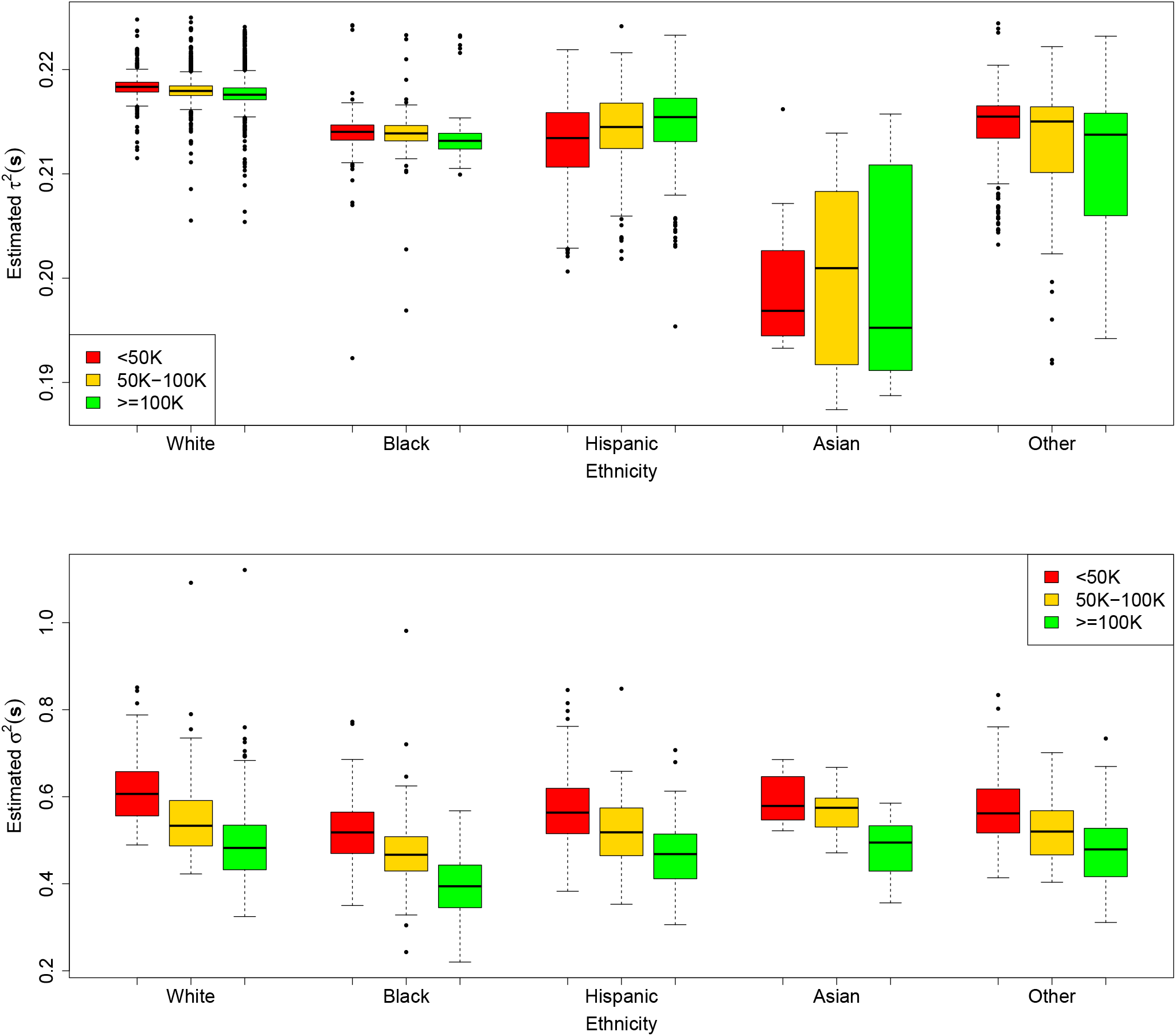
Continues on next page. Boxplots of estimated 1 −*h*^2^(***s***), Var(*y* |***s***), *τ* ^2^(***s***) and *σ*^2^(***s***) for IQ scores in the ABCD dataset. We observe an inverse trend between phenotypic variance Var(*y* |***s***) and income, driven by the non-genetic variance *σ*^2^(***s***). This explains the inverse trend between the estimated extrinsicality 1 − *h*^2^(***s***) and income. Boxplots of estimated 1 − *h*^2^(***s***), Var(*y* |***s***), *τ* ^2^(***s***) and *σ*^2^(***s***) for IQ scores in the ABCD dataset. We observe an inverse trend between phenotypic variance Var(*y*| ***s***) and income, driven by the non-genetic variance *σ*^2^(***s***). This explains the inverse trend between the estimated extrinsicality 1 − *h*^2^(***s***) and income.

**Figure H.29:**
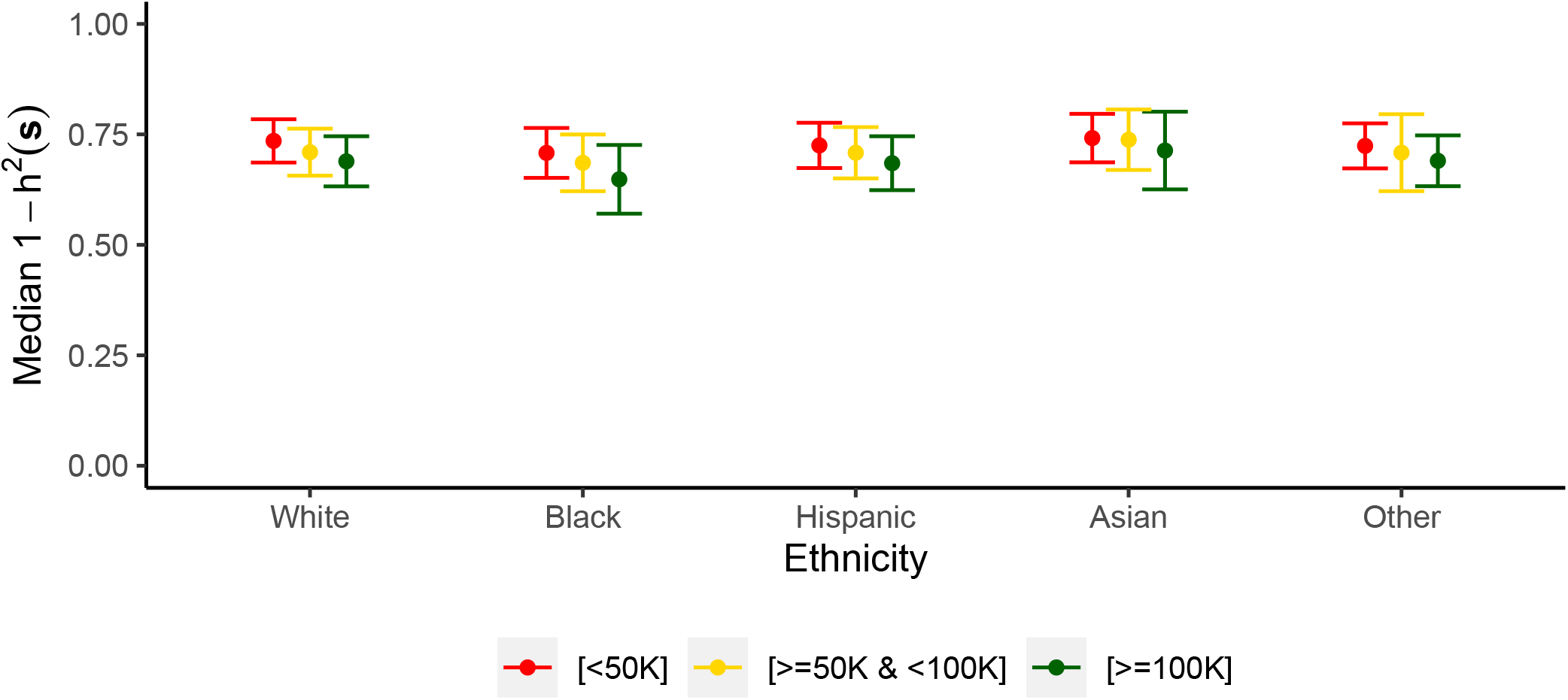
Median estimated extrinsicalities for each income-ethnicity group along with their corresponding estimated standard errors. We observe a similar trend as that illustrated and explained in Figure H.28.

Additional plots for this section can be consulted in Appendix J.

## I Calculation of the recalibrated 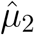 from Section H.0.6

From Section 8.4 of [33] we have the following approximation:

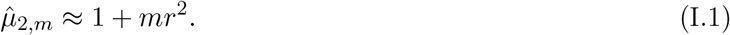

With 872, 188 SNPs and using the 1000 Genomes European sample [7], the authors of [33] obtain 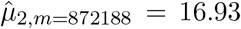. Plugging this into Equation (I.1) and making *r*^2^ the subject of the formula, we obtain:

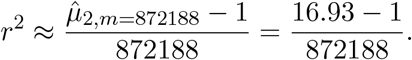

To obtain a *µ*_2_ that is recalibrated for 750, 000 SNPs, we plug this approximated *r*^2^ into Equation (I.1) and set *m* = 750, 000:

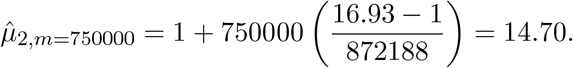

## J Supplementary Figures for Section 3.2

Here we present other figures that could be of interest in addition to those presented in Section 3.2 and Appendix H.

Regarding our estimated standard errors, we can see from Figure J.32 that the standard errors are highest when the estimated extrinsicality is close to 0.5 and then decrease with increasing extrinsicality. Moreover, as shown in Figure J.33 the estimated standard errors seem to be generally higher in the Asian subgroup and lowest in the White subgroup, which can be explained by the difference in number of subjects for each of these (Figure H.23).

**Figure J.30:**
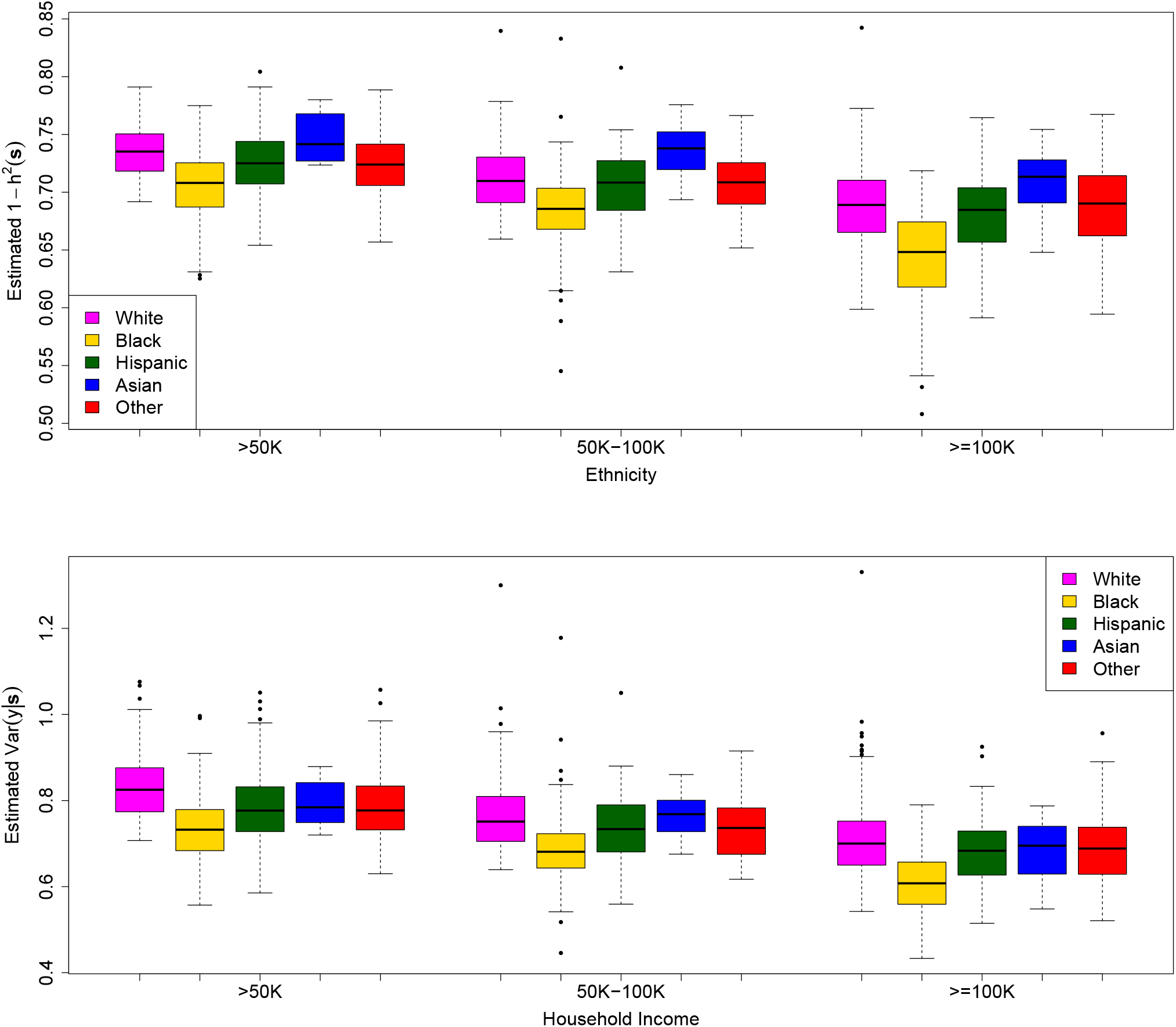
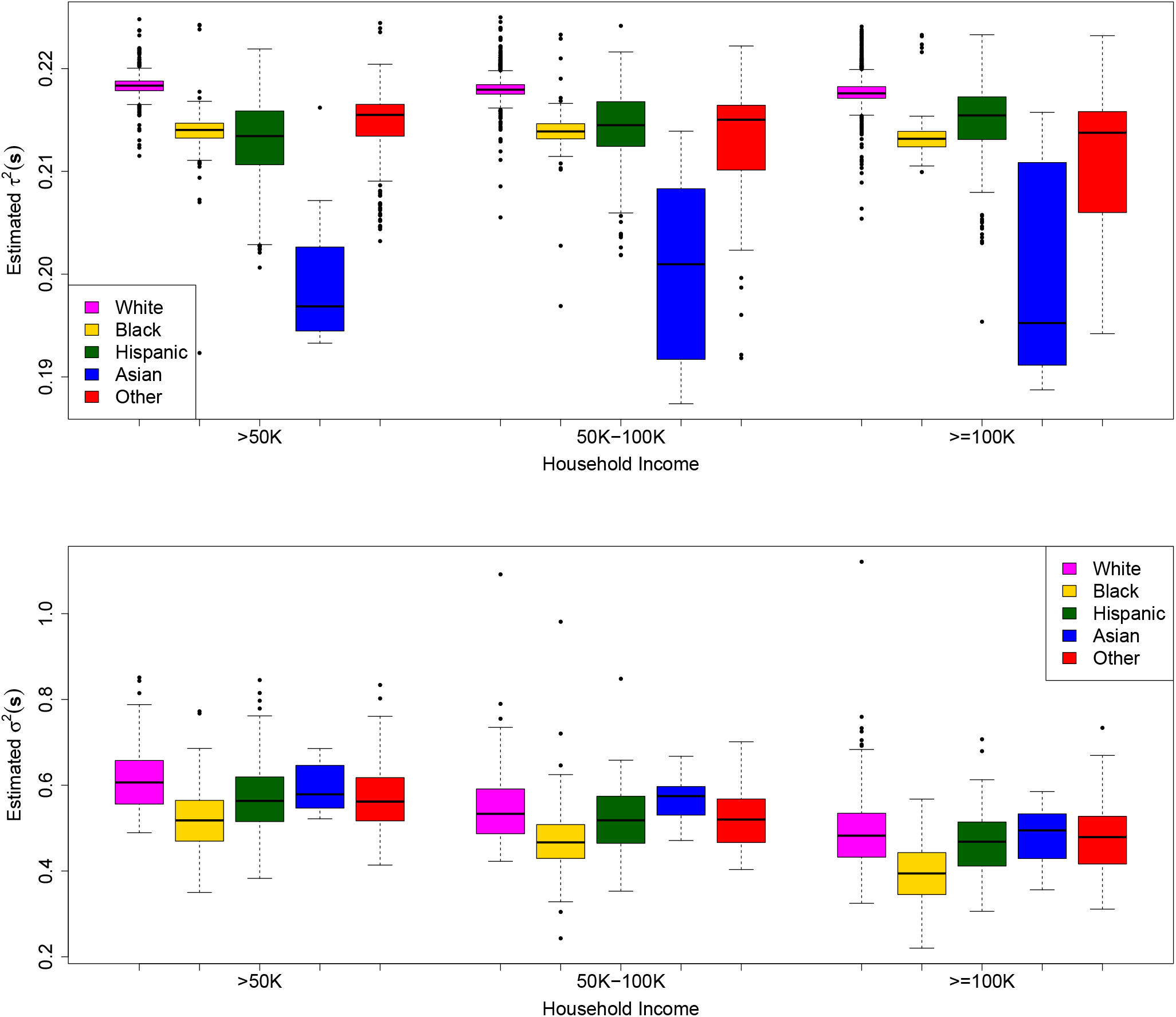
Continues on next page. Boxplots of estimated 1 −*h*^2^(***s***), Var(*y*| ***s***), *τ* ^2^(***s***) and *σ*^2^(***s***) for IQ scores in the ABCD dataset. We group the estimates by income and ethnicity for presentation purposes. We observe an inverse trend between phenotypic variance Var(*y*|***s***) and income, driven by the non-genetic variance *σ*^2^(***s***). This explains the inverse trend between the estimated extrinsicality 1 − *h*^2^(***s***) and income. Boxplots of estimated 1 −*h*^2^(***s***), Var(*y* |***s***), *τ* ^2^(***s***) and *σ*^2^(***s***) for IQ scores in the ABCD dataset. We group the estimates by income and ethnicity for presentation purposes. We observe an inverse trend between phenotypic variance Var(*y*|***s***) and income, driven by the non-genetic variance *σ*^2^(***s***). This explains the inverse trend between the estimated extrinsicality 1 − *h*^2^(***s***) and income.

**Figure J.31:**
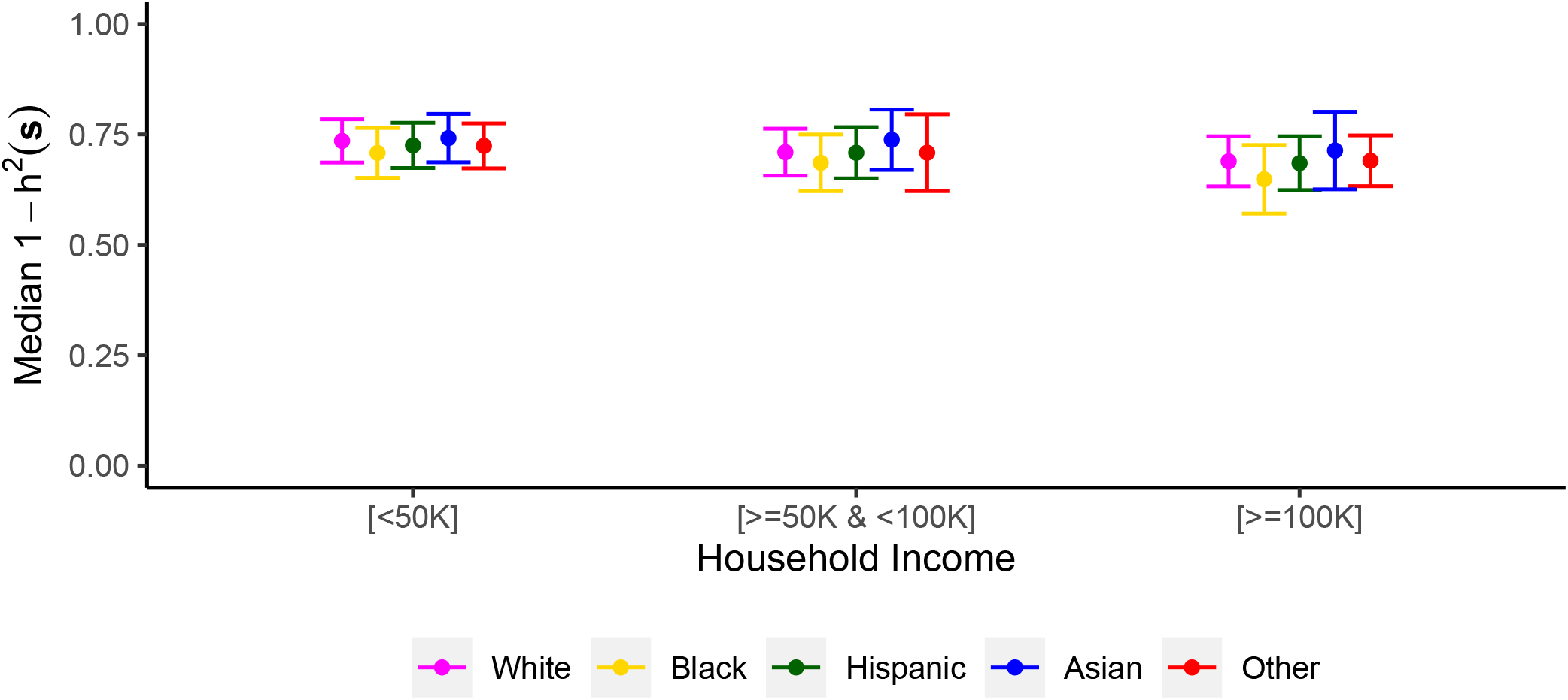
Plot of median estimated extrinsicality in the ABCD data analysis with the corresponding estimated standard error by income and ethnicity. No noticeable differences are observed across ethnicities.

**Figure J.32:**
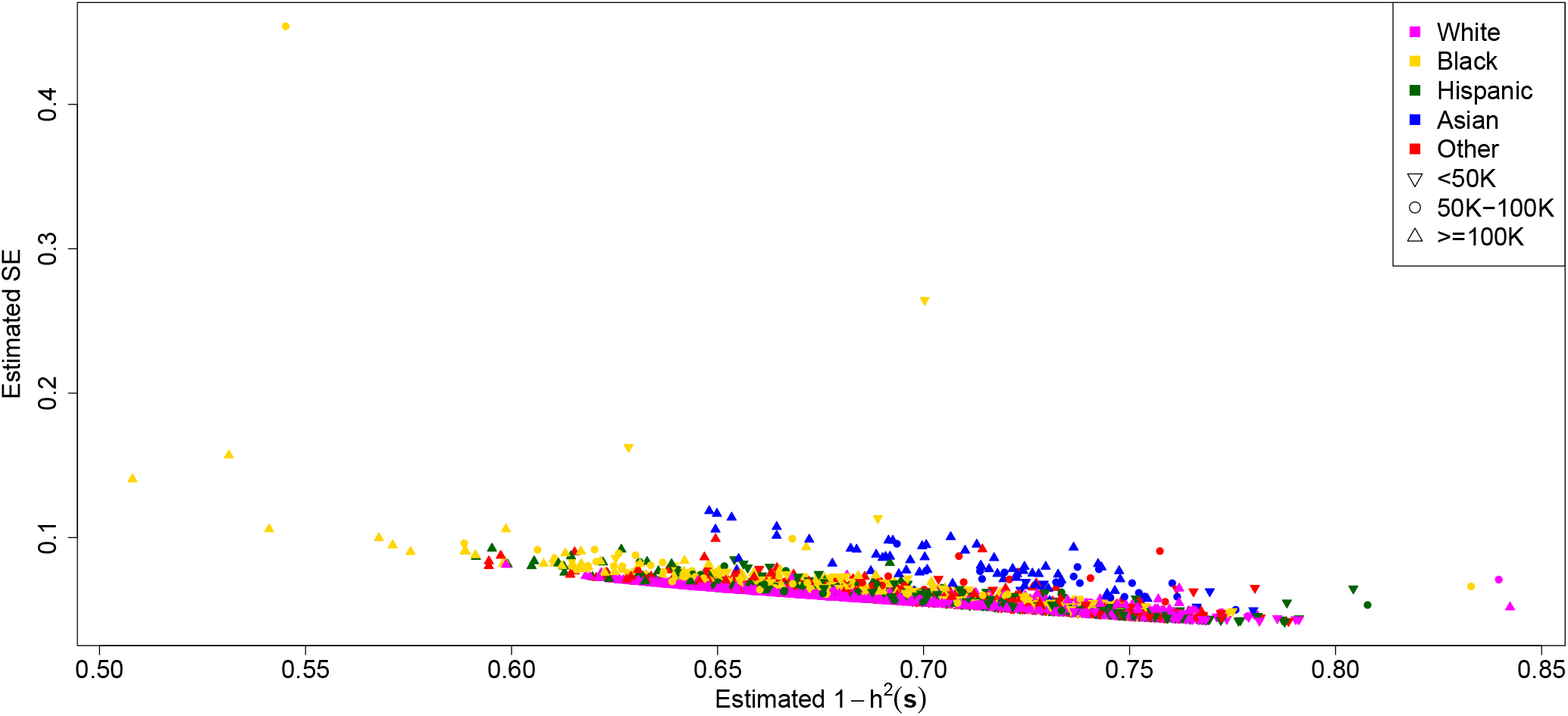
Comparison of estimated extrinsicalities and their corresponding estimated standard errors. Standard errors are highest when the estimated extrinsicality is close to 0.5 and then decrease with increasing extrinsicality.

**Figure J.33:**
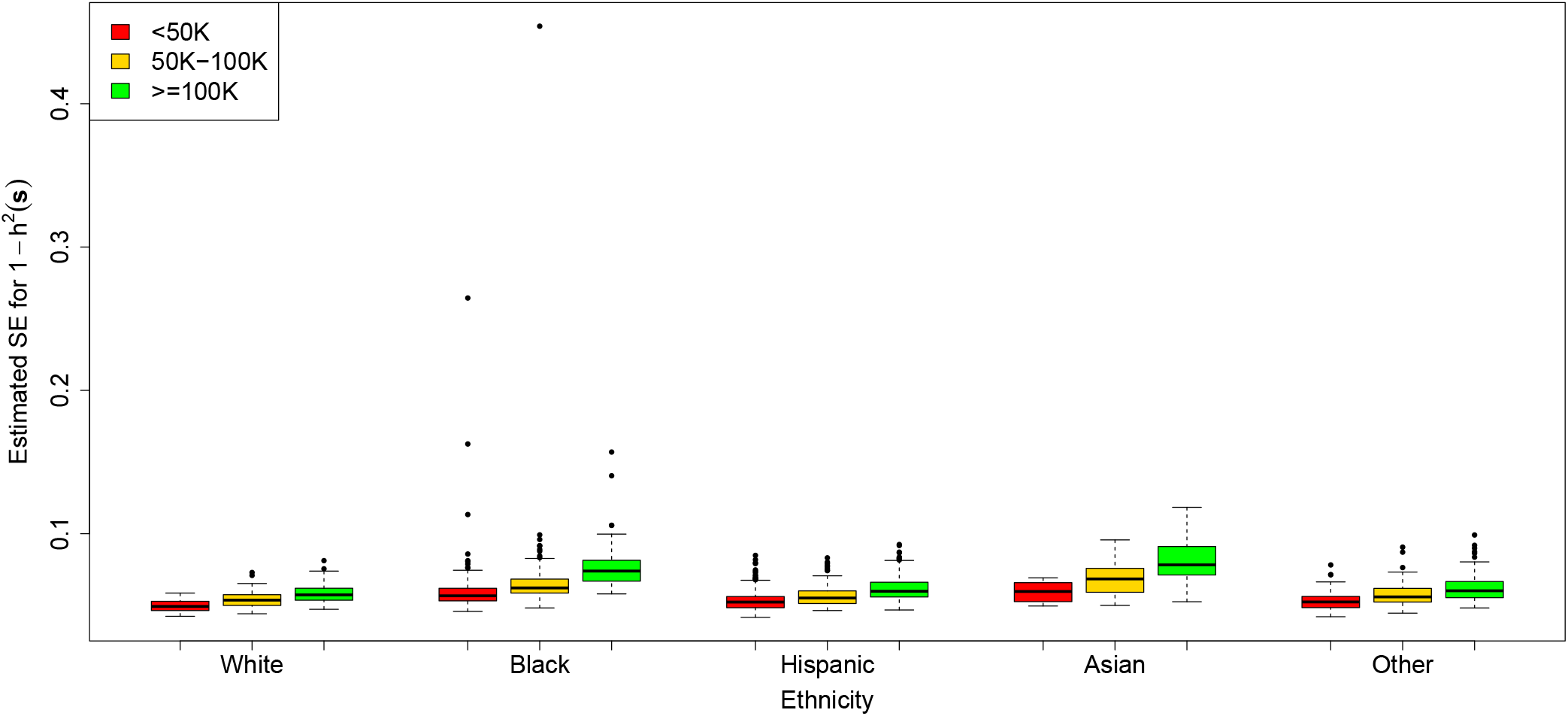
Boxplots of estimated standard errors grouped by income and and ethnicity. The estimated standard errors seem to be generally higher in the Asian subgroup and lowest in the White subgroup, which can be explained by the difference in number of subjects for each of these.

